# Eight Unexpected Selenoprotein Families in ABC transport, in Organometallic Biochemistry in *Clostridium difficile* and other anaerobes, and in Methylmercury Biosynthesis

**DOI:** 10.1101/2022.07.06.499078

**Authors:** Daniel H. Haft, Marc Gwadz

## Abstract

A novel protein family related to mercury resistance protein MerB, which cleaves Hg-C bonds of organomercurial compounds, is a newly recognized selenoprotein, typically seen truncated in sequence databases at CU (cysteine-selenocysteine) dipeptide sites fifty residues before the true C-terminus. Inspection shows this protein occurs in a nine-gene neighborhood conserved in more than fifty bacterial species, taxonomically diverse but exclusively anaerobic, including spirochetes, deltaproteobacteria, and Gram-positive spore-formers *Clostridium difficile* and *C. botulinum*. Three included families are novel selenoproteins in most instances, including two ABC transporter subunits, one a substrate-binding protein with another CU motif, the other a permease subunit with selenocysteine at the substrate-gating position. Phylogenetic profiling shows a strong pattern of co-occurrence with Stickland metabolism selenoproteins, but an even closer link to a group of 8Fe-9S cofactor-type double-cubane proteins. These 8Fe-9S enzymes vary in count and in genome location but frequently sit next to the nine-gene locus. We have named the locus SAO, because of the Selenocysteine-Assisted Organometallic (SAO) biochemistry implied by an uptake ABC transporter with apparent metal-binding selenocysteines, complementary metal efflux pump SaoE, the MerB-like cytosolic enzyme now called SaoL, and comparative genomics signatures suggesting energy metabolism rather than metal resistance. Hypothesizing cycles of formation and dismutation of organometallic compounds involved in fermentative metabolism, we examined methylmercury formation proteins, and discovered most HgcA proteins are selenoproteins as well, with a CU motif N-terminal to the previously predicted start. Seeking additional rare and overlooked selenoproteins, tricky because of their rarity, could help reveal more candidate cryptic biochemical processes.

## Introduction

*Clostridioides* (*Clostridium*) *difficile*, an important intestinal pathogen, can become prevalent after patients are exposed to certain antibiotics, and then maintain itself in a diarrheal disease-causing dysbiosis for extended periods. Several important principles of *C. difficile’s* survival as an obligate anaerobe in the intestinal milieu are well described (1–3). Much of its energy generation depends on the extraction of energy substrates refractory to anaerobic oxidation by most other bacteria. *C. difficile* is a specialist in finding respiratory sinks, and several pathways it contains have been examined in detail. For one, it performs acetogenesis, converting carbon dioxide (CO_2_), via carbon monoxide (CO), plus hydrogen (H2), to acetyl-CoA through the Wood-Ljundahl pathway (WLP) (1). From acetyl-CoA, *C. difficile* does not simply export acetic acid, but instead performs further transformations, generating both ethanol and butyrate (2).*C. difficile* also performs Stickland-type fermentations of a wide variety of amino acids (1,4), coupling some transformations to H+/Na+ transport, which creates gradients that it uses to generate ATP.

Several key proteins used in Stickland reactions invariably require selenium, with key selenoproteins including glycine reductase subunits GrdA and GrdB, sarcosine reductase subunit GrdF, and proline reductase subunit PrdB. Selenium in the form of selenocysteine is used in *C. difficile* also in a formate dehydrogenase and the selenophosphate synthase enzyme SelD, two of the most widespread selenoproteins in bacteria (5). In addition to selenocyteine-containing enzymes, *C. difficile* has the YqeB and YqeC gene pair for preparing the labile selenium-containing molybdenum cofactor (Se-cofactor) for enzymes referred to collectively as selenium-dependent molybdenum hydroxylases (SDMH) (6,7). Characterized SDMH enzymes of this type include nicotinic acid hydroxylase from *Clostridium barkeri* (8), purine hydroxylase from *Clostridium purinolyticum* (9), and xanthine dehydrogenase EF2570 from *Enterococcus faecalis (10)*. The protein CAJ70074.1 from *C. difficile* is 59 % identical to the *E. faecalis* enzyme and annotated as functionally equivalent. However, that enzyme is just one of several candidate SDMH in the proteome, suggesting that several additional selenium-dependent pathways remain to be catalogued in clostridial species.

In light of the broad interest in the metabolic capabilities of *C. difficile*, in the relationships between microbiome-mediated fermentations in the gut and metabolites appearing in host serum, and in new protein family discoveries that expand the selenoproteome (5,11), we were surprised to encounter an apparently overlooked system of nine proteins, six from families whose memberships include selenoproteins some to most of the time, and to find the system closely associated with a still only partly characterized metalloenzyme family that features the highly unusual 8Fe-9S cofactor. The system’s overabundance of Cys-rich and Sec-containing motifs, its MerB-related putative organometallic lyase, its putative metal import and export transporters, and its correlations with anaerobic metabolism, Stickland fermentation, and 8Fe-9S cluster proteins suggest something novel. It suggests a system that exists for metabolic throughput and energy generation rather than heavy metal resistance, one that transports metal-containing compounds or ions both inwards and outwards, a still-undescribed Selenocysteine-Assisted Organometallic (SAO) system whose pathways may involve transient formation and later dismutation of organometallic compounds.

## Methods

### Discovery of novel selenoproteins

Groups of proteins with sequence similarity were detected by BLAST (12) or HMMER (13) searches and aligned using Clustal-W (14) or muscle (15). Multiple sequence alignments were examined, made non-redundant, and trimmed in belvu (16). Any sequence with an unexpected N-terminal or C-terminal truncation, such that the missing region covers what would otherwise be a strongly conserved cysteine (Cys or C) or selenocysteine (Sec or U) residue was considered a candidate selenoprotein. Review of such families looked for 1) at least ten candidate selenoprotein examples in each family, 2) universal presence of TGA (100%) in the DNA encoding the putative selenocysteine site, 3) strong protein-protein sequence similarity running through the putative U residue, showing conservative substitution of amino acid residues and consistent average protein lengths for selenoprotein vs. non-selenoprotein members of the same family. In the cases of a fifth (SaoC) and six (SaoX) selenoprotein in an operon, requirements were relaxed somewhat, but still reported. For SaoC, continued alignment past the Sec residue is not required because the identified Sec residue is the C-terminal residue.

### SECIS element verification

Messenger RNA sequence starting from the UGA codon position were examined manually for regions capable of forming RNA structures that could serve as selenocysteine insertion (SECIS) elements, using dotter (17) for visualization. Searches for SECIS elements were attempted at http://gladyshevlab.org/SelenoproteinPredictionServer/ (18) with a variety of settings but were unsuccessful for both the novel selenoproteins described here and for previously several previously described selenoproteins. Server-based confirmation of putative novel selenoproteins was therefore not used as a criterion for selenoprotein identification.

### Reference data for comparative genomics studies

Three collections of representative genomes were used. The larger collection is the full set of PGAP-annotated prokaryotic assemblies that are in the RefSeq collection (references) and identified as representative, that is, passing a variety of quality assurance metrics on the quality of structural and functional annotation and then chosen as the single representative of that species. This collection contained 15,511 genomes as of May 5, 2022. We will refer to the snapshot from that date, a static set, as “the calibration set”. It is used by NCBI biocurators during the construction, curation, review, and testing of protein family models being developed for PGAP (19).

A collection very similar to the calibration set is available for BLAST searching as “RefSeq Select proteins (refseq_select)” at NCBI’s Web BLAST page https://blast.ncbi.nlm.nih.gov/Blast.cgi?PROGRAM=blastp. Only bacterial assemblies within that collection are relevant to this study. We will refer to this larger set of assemblies as “RefSeq Select.”

Lastly, a collection of 3217 assemblies was built in July 2018 for studies using the algorithm Partial Phylogenetic Profiling (PPP), a data-mining algorithm that looks for statistical evidence that one protein family regularly co-occurs with another, while performing on-the-fly optimizations of protein family size that best reveal what patterns of co-occurrence may be inherent in the genomic data (20,21). We refer to this collection of prokaryotic proteomes as “the PPP reference set.”

### Protein Family Definition

Each protein family definition was created by first building a multiple sequence alignment of representative sequences, then trimming off poorly conserved end regions and eliminating highly redundant sequences. Protein profile hidden Markov models (HMMs) were constructed from these curated alignments, using HMMER3 (13). Curator examination of well-conserved genomic regions was used to guide construction of HMMs that provide sharp separation between members of each family and additional homologs outside the family, if any exist, and then to set cutoffs that would limit or eliminate both false-positive and false-negative scoring of potential member proteins of each family. For families in which previously missed identifications of selenocysteine-containing full-length versions of protein sequences complicated calibration, we first determined what the corrected sequences should be, and then used the corrected sequences to set cutoffs.

### Partial Phylogenetic Profiling (PPP)

PPP was performed in order to see what proteins show the most consistent patterns of co-occurrence with the MerB-like selenoprotein family (SaoL) and with the rest of our newly built protein families from its conserved gene neighborhood. PPP was performed by using a query profile of 33 species encoding the CysCys-COOH protein (SaoC), among the 3217 of the PPP reference set.

BLAST search results needed for PPP were generated for all proteins from each of five taxonomically diverse RefSeq-annotated assemblies: GCF_000009205.1 (*Clostridioides difficile* 630), GCF_000325665.1 (*Brachyspira pilosicoli* P43/6/78), GCF_001754075.1 (*Merdimonas faecis*), GCF_001940565.1 (*Tissierella creatinophila* DSM 6911), and GCF_000014965 (*Syntrophobacter fumaroxidans* MPOB). The PPP software uses the algorithm previously described (20,21) but is a custom implementation that works within NCBI infrastructure, with dependencies that include use of in-house compute farms and relational databases.

The PPP algorithm examines how consistently two protein families co-occur, being both present or both absent, across large numbers of other genomes. The first family is represented by its phylogenetic profile, that is, by the list of all genomes found to contain the family. This profile provides the reference standard used when PPP searches the proteome for any protein that can be seen as belonging to a second family occurring in a similar set of genomes, so the profile functions as a query. PPP examines every protein in a proteome, one at a time, and finds the best possible score for that protein. To find that optimized score, it scans down the descending list of best BLAST hits, finds the genome of origin for every homolog, and computes odds according to the binomial distribution that such an overrepresentation of YES genomes could have been seen purely by chance. For each protein in the genome, the reported score comes from the depth in the scan down the BLAST hits list where the negative of the log of the computed odds hits the highest possible score.

PPP scores are computed for all proteins annotated in a genome, sorted in order from the highest (most significant) PPP score to lowest. Because proteomes from related species, or inhabiting related environments that require similar adaptations, may have considerable overlap in protein family content, no assumption of statistical independence can be made for the distribution of two different protein families among the different proteomes in the test set. Consequently, PPP scores as they are computed, uncorrected, have a high background, and must be used only by comparison to scores for other proteins from the same proteome. The top tier of proteins, those scoring well above background, are reviewed under the assumption that joint appearances across large numbers of genomes may point to a mechanistic connection between those proteins and the one used as the basis for the query profile. Results are interpreted by investigating proteins that score far above this background, after which we build new HMM-based protein family definitions as necessary, with cutoffs set manually only after review of gene neighborhoods and molecular phylogenetic trees.

The assemblies used for PPP have structural annotation that predates the discovery reported here of six new selenoprotein-containing protein families, and are used without any special correction. PPP therefore performs computation on part-length versions of selenoproteins, missing regions either N-terminal or C-terminal to the selenocysteine-encoding UGA codon.

A standard warning applies for all use of PPP, namely, problems that make BLAST search results less reliable make PPP results less reliable. Concerns include suboptimal structural annotation (failure to find all coding regions with correct start sites), repetitive sequences or low-complexity sequence in proteins, or that the order of best hits as computed by BLAST somehow fails to perform as well as desired to separate members of one branch of a large protein family from other branches. For this reason, the BLAST distance-based definitions of protein families that PPP weighs on the fly during computation should be replaced by curator-constructed and validated protein family definitions for downstream uses such as genome annotation and comparative genomics.

### Structure Prediction

3D Protein structure predictions were obtained via Alpha Fold v2 installed on the NIH Biowulf supercomputing cluster, using all available sequence and template databases. The highest confidence 3D model of each protein prediction was submitted to VAST Search (https://www.ncbi.nlm.nih.gov/Structure/VAST/vastsearch.html) (22), which compares the 3D coordinates of a submitted protein structure (including those modeled by AlphaFold) to all solved 3D structures in NCBI’s Molecular Modeling Database, to help identify structure neighbors of interest. 3D visualization was performed using iCn3D for most proteins (23,24), while the structural alignment and overlay visualization of SaoP with the MetI dimer used ChimeraX (25).

## Results

### MerB-like selenoprotein (SaoL)

Pfam (26) model PF03243 identifies proteins of the MerB family. Characterized members of that family described in the literature (27) are organomercurial lyases (EC 4.99.1.2) able to cleave substrates such as methyl-mercury. MerB-like proteins WP_004454659.1 from *Clostridioides difficile* (Firmicutes), WP_041443257.1 from *Syntrophobacter fumaroxidans* (Deltaproteobacteria), and WP_228369505.1 from *Brachyspira pilosicoli* (Spirochaetes) predicted previously by PGAP (19) represented a distinct branch within the alignment of MerB-like proteins. All predicted proteins in this branch were somewhat shorter than the typical MerB, averaging 197 residues in length, and all ended in a perfectly aligned Cys. Since some selenoproteins contain a CU (Cys-Sec) or UC (Sec-Cys) dipeptide motif, we investigated the possibility that all were, in fact, previously unrecognized and therefore improperly truncated selenoproteins. In every case, the original stop codon was UGA, which can encode selenocysteine, followed by a resumption of the open reading frame. Translations with a single UGA codon interpreted as selenocysteine produced alignments with homology extending to the C-terminus of MerB-like proteins such as WP_008520765.1 of *Jonquetella anthropic*, WP_208119251.1 of *Spirabiliibacterium pneumoniae*, and WP_151052138.1 of *Neisseria zalophi*.

By examining the genomic regions surrounding WP_004454659.1, WP_041443257.1, WP_228369505.1, and their closest homologs, we discovered that the sequences can be treated as the founding members of a novel protein family that serves as a molecular marker for a large conserved gene neighborhood. The neighborhood occurs 52 times in a collection 15,511 representative prokaryotic genome assemblies. This is just 0.3 % of species, all anaerobic, skewed heavily toward the Firmicutes but broadly distributed. Examination of neighborhoods showed that the sequence mentioned above from *Jonquetella anthropici*, WP_008520765.1, shares the same gene neighborhood and is the lone member of the SaoL family that that has a CC rather than CU motif. *Jonquetella anthropici* does have a selenocysteine incorporation system for production of other selenoproteins.

Surprisingly, a member of this MerB-like (seleno)protein is found in only 45 of the 52 conserved regions. It occurs as a selenoprotein in 44 of the 45 (98%). The full list of hits in RefSeq can be found by searching Protein Family Models at https://www.ncbi.nlm.nih.gov/protfam for hidden Markov model NF040728. The nine novel protein families of the neighborhood, with suggested gene symbols, protein names, accession numbers of HMMs identifying protein family members, and other notes, are listed in **Table 1**. All members of all nine family are found exclusively in bacteria. Note that we have added corrected full-length sequences that span the selenocysteine insertion site as reference information for PGAP, so the pipeline now correctly produces full-length proteins when PGAP annotates new prokaryotic genome assemblies entering RefSeq or when it does its regular reannotations of existing assemblies.

**Table 1.**

| Protein Accession | HMM | gene | provisional protein name | U/C+U |
| --- | --- | --- | --- | --- |
| WP_012048219 | NF040730 | saoT | thioredoxin-like protein SaoT | 34/52 |
| WP_012048218 | NF040736 | saoE | ArsP-related export pemease | 0/52 |
| WP_003357554 | NF040734 | saoC | Cys-Cys-COOH protein SaoC | 3/52 |
| WP_012048217 | NF040735 | saoX | ABC transporter substrate-binding protein SaoX | 2/53 |
| WP_240004094 | NF040728 | saoL | MerB-related organometallo-lyase SaoL | 44/45 |
| WP_242649186 | NF040727 | saoB | ABC transporter substrate-binding protein SaoB | 31/52 |
| WP_242649185 | NF040733 | saoP | ABC transporter permease subunit SaoP | 44/52 |
| WP_012048215 | NF040729 | saoA | ABC transporter ATP-binding subunit SaoA | 0/52 |
| WP_003357559 | NF040732 | saoD | DsrE-related protein SaoD | 0/35 |

The MerB-like selenoprotein turned out to be the first novel selenoprotein we found in its conserved gene neighborhood. It serves as a molecular marker for the presence of an extended locus at least nine genes in length, several others of which also encode selenoproteins, as will be described. We infer that the system on the whole acts on organometallic (probably organomercurial) compounds, as MerB itself does, and suggest the term ***SAO*** (**S**elenocysteine-**A**ssisted **O**rganometallic) as a shorthand name for the system on the whole. For this MerB-like protein, we suggest the name “SaoL”, for “**S**elenocysteine-**A**ssisted **O**rganometallic Lyase.”

### 3D structural prediction for SaoL

The corrected, full-length, selenocysteine-containing protein SaoL of *Clostridioides difficile* 630 is WP_238476746.1, 213 amino acids long with a CU motif at position 161-162. The U in WP_238476746.1 was changed to C, producing a sequence we designate SaoL_Cdiff_Sec162Cys, in order to avoid having AlphaFold treat the selenocysteine as X. **Figure 1** shows structures for SaoL_Cdiff_Sec162Cys, as predicted by AlphaFold2, and for MerB from *Escherichia coli* plasmid R831b, solved crystallographically with a bound mercury atom as PDB structure 3F0P. The two sequences, although homologous and both recognized by Pfam model PF03243, share less than 15% pairwise identity in amino acid sequence. However, homology is readily detected by BLAST search with SaoL_Cdiff_ or HMM NF040728 vs. sequences represented in PDB, while VAST (Vector Alignment Search Tool) (28) finds the similarity of the two 3D structures.

**Figure 1:**
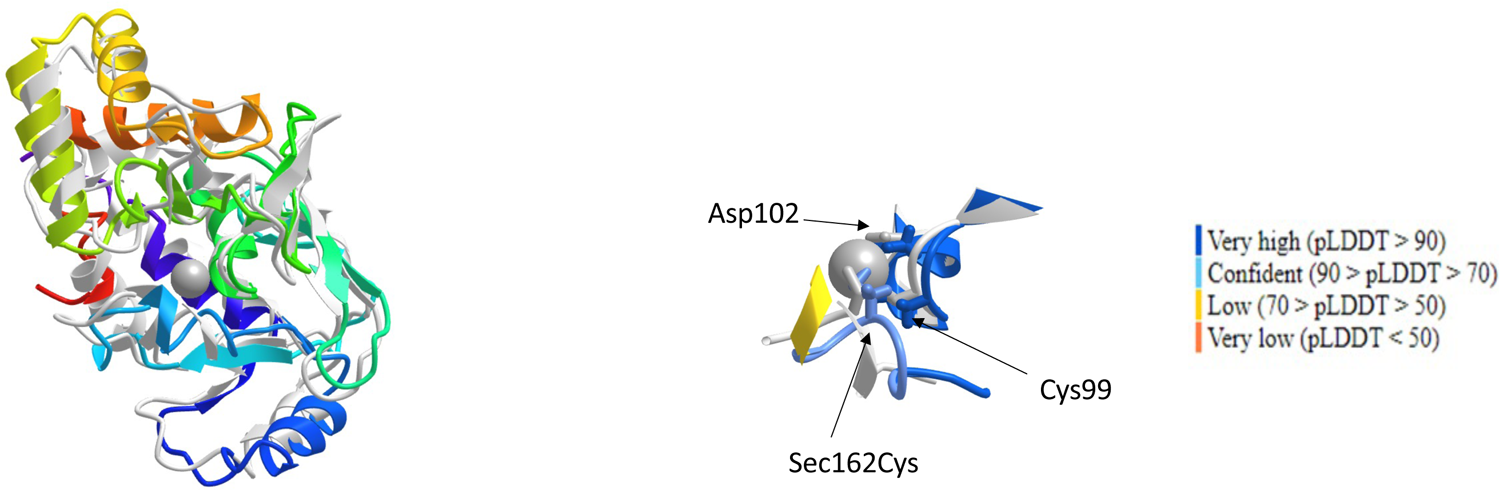
AlphaFold2 structure of SaoM_Cdiff_Sec162Cys with structural alignment to MerB structure 3F0P. The left panel shows alignment of the complete structures, with SaoM rainbow-colored from N-terminus (red) to C-terminus (purple), while MerB is colored gray. The right panel shows closeups of the Hg atom bound by MerB from the same superposition of structures. In this panel, MerB and the Hg atom are gray, while SaoM_Cdiff_Sec162Cys is colored by AlphaFold2 pLDDT scores of local structural prediction confidence (see key). SaoM residues Cys99, Asp102, and Sec162Cys (indicated by arrows) align to MerB’s active site residues Cys96, Asp99, and Cys159.

MerB enzymes acts as organomercurial lyases for methylmercury and a broad variety of related organomercurial substrates. MerB’s binding to and positioning of active site residues at the mercury atom is paramount. It is notable that MerB will bind other heavy metal atoms where it binds Hg (29), a single amino acid change at the metal-binding site can convert MerB to a copper-binding enzyme (29), and the ability of heavy metal atoms for bond covalently to carbon is not limited to mercury, but includes also, for example, arsenic and cobalt. The replacement of Cys by Sec at a site contacting the heavy metal atom may be a profound change, so it increases suspicion that the physiologically relevant substrate profile may not be restricted to organomercurial compounds. Indeed, mercury resistance operons frequently are encoded on plasmids, and for well-studied human pathogens tend be present only sporadically. By contrast, the SAO system in *C. difficile* strains is chromosomal, surrounded by housekeeping-like rather than mobilome-like genes, and preserved in 3279 of 3343 genome assemblies (98%) for that species annotated by PGAP for the RefSeq collection.

### CD2361 family ABC transporter ATP-binding protein (SaoA)

Homologs of CD2361 (WP_003430772.1) are the ATP-binding subunit of an adjacent ABC transporter of unknown substrate. None of the members of this family are selenoproteins, and that is expected. The ATP subunit energizes high-affinity transport by its effects on other subunits, and it does not interact directly with substrate. High sequence similarity to other nitrate/sulfonate/taurine group ABC transporter subunits helps determine that adjacent genes likely also encode subunits of an ABC transporter, although the substrate remains unknown. We suggest the name SaoA (**SAO** system **A**TP-binding protein).

### ABC transporter permease selenoprotein (SaoP)

The adjacent ABC transporter permease subunit, originally predicted to be 209 amino acids in C. difficile (WP_003430774.1), was corrected to a length of 268 by translation through the selenocysteine-encoding UGA codon. Members of the family are mutually more than 45% identical to each other. In a few members of the family (WP_008522719.1, WP_015556508.1), Cys is present rather than Sec, but the majority of family members are selenoproteins. We suggest the name SaoP (Selenocysteine-Assisted Organometallic transport Permease).

Pairwise sequence alignment of the sequence to proteins with a solved crystal structure, such as the MetI subunit of the MetNI methionine ABC transporter shows the highly conserved selenocysteine in WP_003430774.1 aligning within a residue of Met-163 of MetI. MetI forms a homodimer. In the solved conformation, open toward the cytosol and closed on the extracellular face, side chains of the Met-163 residues to the two MetI monomers pack together, as that residue performs a gating function (30).

The positioning of SaoP’s selenocysteine side chain in a gating position becomes even more vivid after structure prediction. Because SaoP is the only ATP transporter permease in the SAO locus, it too is expected to form a homodimer. The top-scoring AlphaFold prediction for the SaoP homodimer, as seen in **Figure 2**, puts the permease in an open-inward (toward the cytosol) configuration. The panel on the left shows excellent superposition with the solved structure 3DHW of MetI. The alignment of SaoP to MetI based on their structures aligns Met-163 of MetI in PDB crystallographic structure 3DHW to Cys-210 (standing in for Sec-210 for software compatibility) in the AlphaFold structure of SaoP. The right panel shows a top-down view of the SaoP dimer. The selenocysteine side chains of the two SaoP monomers appear close enough to form a diselenide bond, or to have both contact the same putative metal atom of a possible organometallic transporter substrate.

**Figure 2:**
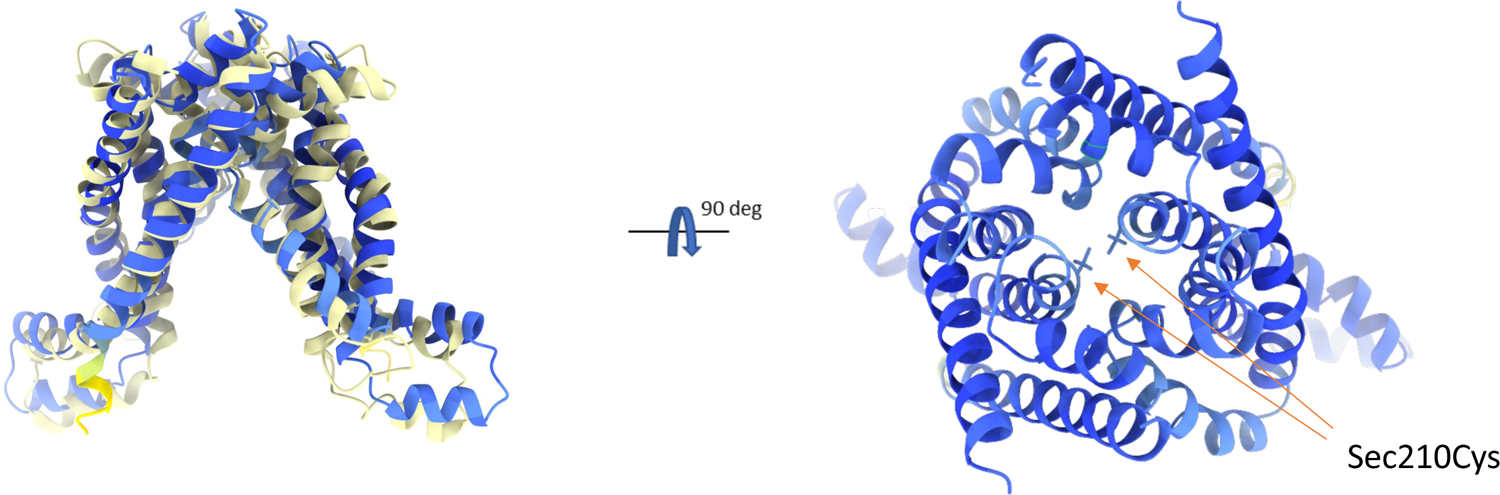
AlphaFold2 prediction for SaoP_Cdiff_Sec210Cys as a homodimer and structural alignment to structure 3HDW for the Escherichia coli ABC transporter permease subunit MetI in the open-inwards conformation. The left panel shows a side view, with the bottom facing the cytosol. The right panel shows a view of homodimeric SaoP_Cdiff_Sec210Cys as seen top-down from outside the plasma membrane after rotation 90^0^. Amino acid side chains are shown only for the Cys residue that represents Sec210, which occupies that same position in the center of the image as the permease-gating residue Met-163 in the MetI solved structure.

**Figure 3.**
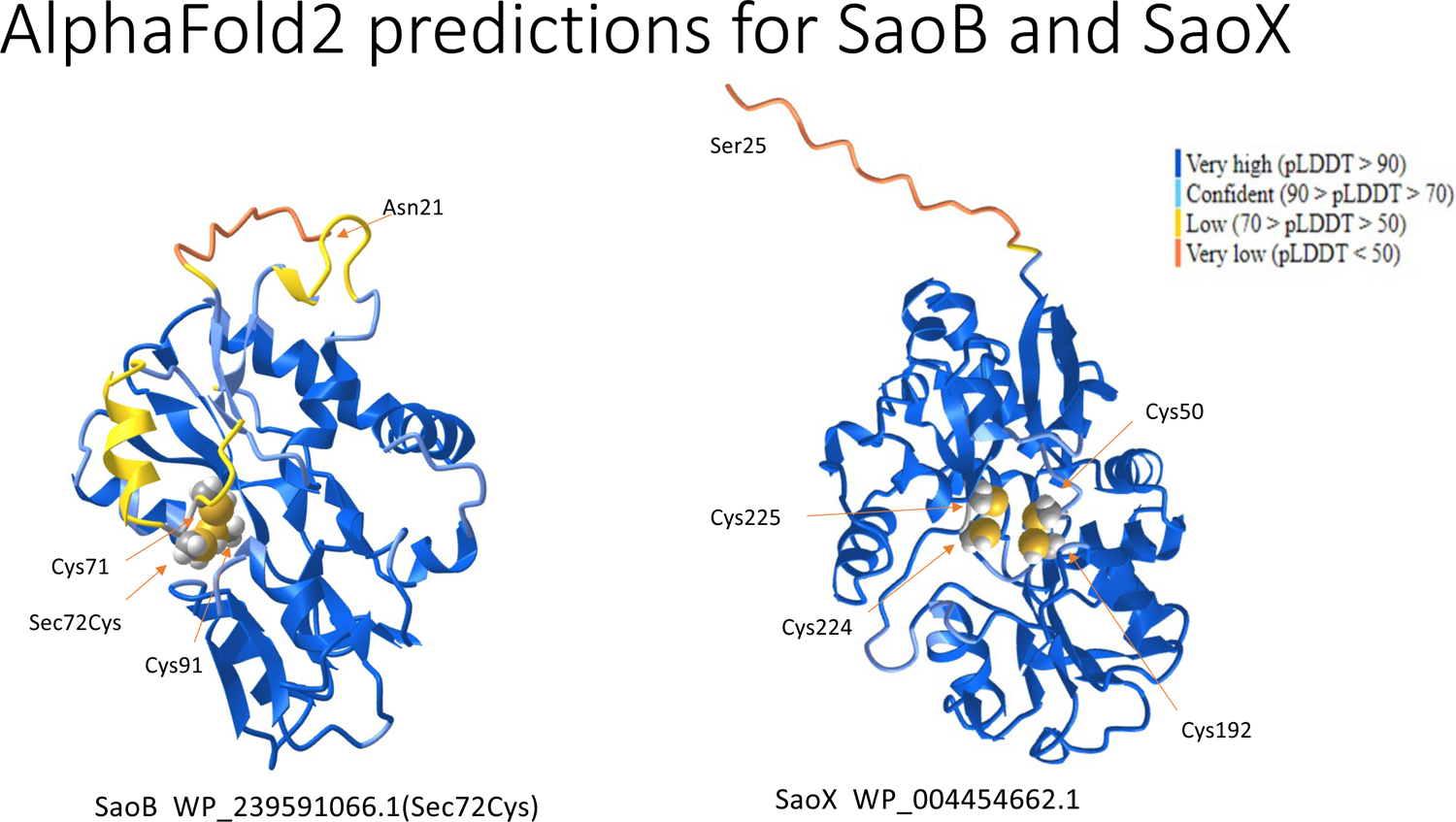
AlphaFold2 predicted structures for putative ABC transporter organometallic substrate-binding proteins SaoB and SaoX. Coloring reflects the pLDDT metric of confidence in the local structure. The left panel shows a prediction based on *C. difficile’s* SaoB selenoprotein WP_239591066.1, after trimming an N-terminal signal sequence and replacing Sec72 with Cys. Locations are shown for Cys/Sec residues conserved across the SaoB family, found at positions 71, 72, and 91. All S/Se atoms are near each other and may help form a metal-binding site in a substrate-binding pocket or cleft. A fourth Cys residue is not shown because it is not well conserved in a multiple alignment of SaoB sequences. The right panel shows the lipoprotein SaoX of *C. difficile*, WP_004454662.1, after trimming the predicted lipoprotein signal peptide. An additional N-terminal region is predicted with low confidence (reddish-orange color) and not made part of the globular structure. The rest packs in a structure highly similar to ABC transporter substrate-binding protein PDB structures, such as 4NMY and 2X26. Four conserved Cys residues are found at positions 50, 192, 224, and 225, and their side chains conserve to form what again appears to be a metal-binding site. In about 40% of SaoX proteins, the residue preceding Cys-192 is cysteine as well, forming a CC motif.

Selenocysteine residues are extremely uncommon in any type of transporter. We know of no previous example of an ABC transporter subunit containing a selenocysteine residue. The presence of two selenocysteine residues, positioned so they can form a diselenide bond, or both can serve as ligands to the same moiety in whatever substrate is being imported, is striking. This positioning of selenocysteine at such a critical site suggest that chemistry involving the pair of selenium atoms is critical to both gating and transport by this novel permease.

### ABC transporter substrate-binding protein (SaoB)

Upstream of the permease subunit is a protein we judge to be the substrate-binding protein (SBP) of the ABC transporter whose other subunits are SaoA and SaoT. The two previously predicted proteins in *C. difficile*, WP_009897548.1 and WP_009897547.1, are actually N-terminal and C-terminal regions within WP_239591066.1, which we were able to make full length after recognizing the selenocysteine-containing nature of the protein. We designate this protein SaoB (Selenocysteine-Assisted Organometallic transport substrate-Binding protein). Homology between SaoB and previously reported ABC transporter SBPs is remote. Two rounds of PSI-BLAST (12) of WP_239591066.1 vs. RefSeq_Select, using an inclusion threshold of 1e-8, followed by multiple sequence alignment of homologs with E-values 0.1 or better, yielded 85 sequences, which became 49 after removing fragmentary sequences and redundant protein sequences more than 80% identical to others. The resulting multiple alignment showed distant but convincing sequence similarity across the full length of the multiple sequence alignment (data not shown), suggesting homologous origin and similar 3D structure, to proteins such as WP_048312536.1, which in turn has very strong sequence similarity (E-value < 1e-43) to SBP with solved structures, such as PDB:3VVE and PDB:4ZV2.

The selenocysteine-containing motif in SaoB family protein WP_009897548 is CU, echoing the CU motif seen in the MerB family selenoproteins. As with the permease subunit, in which two Sec or Cys residues come together by dimerization, the substrate-binding protein also seems able to provide two (seleno)cys ligands to a single metal atom. More than 60 % of the members of the SaoB family are selenoproteins. As with ABC transporter permease subunits, we know of no previous example of any ABC transporter SBP containing selenocysteine.

We performed structural predictions for SaoB_Cdiff_Sec72Cys, a sequence derived from the *C. difficile* selenoprotein WP_239591066.1. A long N-terminal region including the signal sequence was predicted at low confidence and apart from the main globular structure, so the signal sequence was removed and fold prediction repeated. **Figure 2** shows to the left a structural prediction for SaoB_Cdiff_Sec72Cys with the signal sequence removed, and the right, a prediction for SaoX protein WP_004454662.1 with its lipoprotein signal sequence removed. A VAST search for PDB structures similar to the prediction made for SaoB_Cdiff_Sec72Cys shows that the most closely related structures belong to other ABC transporter substate-binding proteins, confirming observations made by sequence comparisons. In SaoB_Cdiff_Sec72Cys, cysteine residues occur at positions 71, 72, 91, and 174. The first three Cys residues are well-conserved in a multiple sequence alignment of SaoB family proteins, and their side chain sulfur atoms are shown to be close to each other in the fold prediction, packing around a probable substrate-binding site. Cys-174 is not conserved in multiple alignments, and is therefore not highlighted in Figure 2, but comparison to other solved crystal structures suggests it also is a likely close contact to substrate. The companion image in Figure 2, from SaoX (described below), which also has conspicuous conserved Cys residues, shows a similar convergence of Cys residue side chains in the substrate-binding pocket.

### CD2365 family substrate-binding (lipo)protein (SaoX)

Members of the family of CD2365 from *C. difficile* 630 (CAJ69250.1) nearly always are lipoproteins. The family is related to a number of ABC transporter substrate-binding proteins, which would make it an extra SBP in a system with only one ABC transporter permease subunit and only one ATP-binding protein. The protein is designated SaoX (SAO system eXtra binding-protein). Because SaoX is a lipoprotein, it was studied as part of the *C. difficile* lipoproteome. It was found to be expressed in vegetative cells and at even higher levels in spores (31,32). This lipoprotein may serve, fortuitously, as a marker for expression of the SAO system. It seems to suggest that the whole operon is transcriptionally active and expressed as protein, and that the system has a meaningful role in host cell metabolism, even during spore outgrowth. SaoX has a conserved CC motif at positions 224-225. Cys-192 has an adjacent Cys at 191 in about half of SaoX sequences. SaoB and SaoX therefore differ from any of 27 other annotated ABC transporter SBP in the *Clostridioides difficile* 630 genome in having a CC or CU motifs. Double-cysteine motifs appear to be uncommon in SBP, and do not occur in the 38 ABC transporter SBP of *Escherichia coli* K-12.

Of the 52 annotated bacterial assemblies in the calibration data set that contain the nine-gene system, a single species, *Negativibacillus massiliensis*, appeared to lack a substrate-binding lipoprotein from this family as annotated. What it had instead as originally annotated is two consecutive tandem pairs of proteins, WP_022132697 and WP_022132698, followed by WP_022132699 and WP_022132700, with the upstream protein terminating in a TGA codon that aligns to the second Cys of the more N-terminal CC motif in the CD2365. Correcting the structural annotation in *N. massiliensis* gives two novel putative selenoproteins, WP_248000900 and WP_248000901, full-length homologs of CD2365, with TU at 191-192 where many homologs have CC.

### *CD2367 family efflux transporter* (SaoE)

CD2367, encoded two genes further upstream, appears to be a transmembrane transporter, but not an ABC transporter. According to TCDB (33), members of that family, such as WP_022618526.1 from *C. difficile*, belong to the Organo-Arsenical Exporter (ArsP) family. While members of CD2367’s family do not include selenoproteins, the most closely related described family among characterized proteins, the SO_0444 family Cu/Zn efflux transporters, does contain selenoprotein examples. Intriguingly, the SO_0444 family is one of only two prokaryotic transporter family we were able find previously through our combination of literature review and legacy data review and validate for inclusion in reference data that NCBI’s Prokarotic Genome Annotation Pipeline (PGAP) uses to detect new occurrences of known selenoproteins (34). The other is a small set of proteins related to the mercury resistance transporter MerT. We suggest the name SaoE (SAO system metal Efflux protein) for CD2367-like proteins.

The tripeptide Cys-Ser-Cys (CSC), which is the single motif most strongly conserved between the SO_0444 and SaoE families, occasionally is replaced by a selenocysteine version, Sec-Ser-Cys (USC) in the SO_0444 family. Selenoprotein examples from the SO_0444 family include WP_014809615.1 from *Desulfomonile tiedjei* and all of its closest homologs. We suggest that SaoE family permeases, while lacking selenium in all species we examined, are involved in efflux of a potentially toxic heavy metal. The absence of selenocysteine from this family means even a total lack of selenium would have no effect on translation of this protein and would not block its ability to remove heavy metal (such as mercury) ions from the cytosol.

Although it seems highly likely that SaoE is involved the export of either some metal-containing substrate, it is unclear whether its primary physiological substrates are inorganic compounds, or organometallic. The exporter SaoE is never found unless import proteins SaoP, SaoA, SaoB, and SaoX also occur, suggesting the need for its metal efflux activities depends on the presence of the ABC transporter.

### CD2366 family CC-COOH protein (SaoC)

Homologs of CD2366 were previously protein bioinformatics “dark matter”, with no available crystal structure, no relevant literature detectable by PaperBLAST (27), no protein family models defined by major protein family databases such as Pfam, NCBIFAMs (which includes TIGRFAMs), CDD, or InterPro. PSI-BLAST searches do not reveal any homologs beyond those found in the conserved gene neighborhood described in this paper. All have signal peptides, most of which appear to be lipoprotein signal peptides. The average length is 158. The protein is designated SaoC (SAO system CysCys-COOH protein), as nearly all end with a double-cysteine motif.

In the set of SaoC proteins from 52 representative species, just three terminate with a single Cys residue instead of a CC pair. The reported coding regions for all three of these exceptional members, WP_068474724.1, WP_055078435.1, and WP_070088375.1, end with a double stop codon, UGA-UAG. Because the UGA codon aligns to what is otherwise an invariant Cys residue in an invariant Cys-Cys motif, a selenocysteine insertion site is suspected. The 43 base nucleotide sequence starting from the UGA codon of WP_055078435.1, UGAUAGUAUUUUU **UUACUUG** GGU **CAAGUAA** GGAGAGAUUAUGA, shows a perfectly based paired stem of length seven (in boldface), with a G-containing loop GGU, as expected for a SECIS (selenocysteine insertion sequence) element in Gram-positive species (35). Thus, even in the absence of a continuation of homology C-terminal to the proposed selenocysteine, and with the relatively small number of proposed selenoproteins seen in this family, we consider the confidence that SaoC family proteins include selenoproteins to be high, as there is good precedence for a Cys-Sec-COOH motifs in selenoproteins. Two small DUF466 family proteins are encoded in Escherichia coli K-12, YbdD and YjiX, each ending Arg-Cys-Cys-COOH. Each is encoded next to a pyruvate uptake transporter, either CstA or its paralog (36). A related DUF466 family selenoprotein occurs as a selenoprotein in *Campylobacter* and some *Helicobacter* species, with the ending Lys-Cys-Sec-COOH (37). While the SaoC and Lys-Cys-Sec-COOH family selenoproteins exhibit no detectable homology, their shared proximity to transporters is intriguing.

SaoC family selenoproteins are the fourth set in this system with a CU motif, in addition to the permease subunit in which U is flanked by residues other than Cys, but which dimerizes so that the selenocysteine site chains of the two different monomers could be just as close. The repetition of CC and CU motifs in so many of the protein families in this nine-gene system is highly suggestive.

### CD2368 family thioredoxin-like protein (SaoT)

The 61-amino acid predicted protein WP_003432954.1 from *C. difficile* is incomplete, as it is yet another selenocysteine-containing protein family. WP_012100993.1 from *Clostridium botulinum* and WP_011700724.1 from *Syntrophobacter fumaroxidans* are full-length (about 85 residues) and lack selenocysteine residues, but corrected selenoprotein sequences include WP_240067706.1 from *C. difficile*, WP_240067707.1 from *Peptacetobacter hiranonis*, and WP_240067709.1 from *Negativibacillus massiliensis*. A search with the HMM we built to detect members of this family, NF040730, detects sequence similarity, and apparent homology, to MJ0307, from *Methanococcus jannaschii*, which has a thioredoxin-like activities (38) and a solved structure (39). However, variations of the typical CXXC motif of thioredoxins and glutaredoxins that are seen in CD2368 family proteins include UPTC, CAUC, CPTC, and also CPCC. The last of these forms, CPCC, overloads the CXXC with a third Cys residue, and is the most common form across the family. By contrast, in the MJ0307-like family and among all other prokaryotic proteins in the RefSeq collection whose names contain the words “thioredoxin” or “disulfide”, a CXXC motif also never has an addition Cys in the second or third position. Less than 0.5% of CXXC motifs in such proteins have the form CCXC or CXCC. The contrast suggests that while the fold may be similar, SaoT may be undergone a change in function relative to more typical thioredoxin-like ancestral sequences, perhaps now involving direct interaction with the same family of metal-containing species as other members of the SAO system.

### CD2369 family DsrE-like protein (SaoD)

Homologs of the C. difficile protein CD2369 are never detected as selenoproteins. They have an invariant CxxC motif that sometimes takes the form CxxCC, and another invariant Cys residue. The family is related to DsrE proteins identified by Pfam model PF02635, and therefore designated SaoD (SAO system DsrE-like protein). Characterized members of the DsrE family have sulfur-transfer abilities used in processes such as tRNA modification, in the cases of TusA, TusC, and TusD in E. coli, and in sulfur oxidation.

In contrast to other proteins in the SAO system, the SaoD family has a number of fairly closely related homologs found in species lacking the system, although members of the SAO system can be separated from non-members in molecular phylogenetic trees made from a multiple sequence alignment. For member proteins of the related branch found outside the set of species with the SAO system, the most frequently observed neighboring genes include homologs of metal import ABC transporter substrate-binding proteins such as AztC, metallochaperones such as AztD, and ABC transporter permease subunits for various metals. This observation further supports the notion that the SAO system interacts, at least partly, with inorganic substrates or organometallic compounds.

SaoD, like the putative organometallic lyase SaoL, is one of two proteins in the system not universally present across the 52 representative species.

**Table S1**, in Supplemental Materials, shows proteins identified by the nine new HMMs, providing identities of 52 species with the conserved gene neighborhood, accession numbers for all member proteins found in the SAO locus in those organism, and identities of proteins from additional correlated protein families such as SelD, GrdX, and double-cubane proteins identified by phylogenetic profiling and gene neighborhood analyses.

### Partial Phylogenetic Profiling results

A phylogenetic profile was constructed by use of the new NCBIFAMs model NF040734, which finds members of the SaoC family of CysCys-COOH proteins. The profile included 33 YES genomes, from 3217 genomes in the PPP genomes, for a probability *p* of about 0.01. PPP was run with the YES frequency parameter elevated to the manually set value of 0.03. Increasing this parameter, used in binomial distribution computations by PPP, compensates for an intrinsic bias problem in comparative genomics studies from the fact that species from the same lineage share larger numbers of closely related protein pairs, simply because of recent common ancestry.

Partial Phylogenetic Profiling (PPP) studies were performed on the proteomes of assemblies GCF_000009205.1 *(Clostridioides difficile)*, GCF_000014965.1 *(Syntrophobacter fumaroxidans)*, GCF_000325665.1 *(Brachyspira pilosicoli)*, GCF_001754075.1 *(Merdimonas faecis)*, and GCF_001940565.1 *(Tissierella creatinophila)*. **Table S2** in Supplementary materials lists the 40 top-scoring proteins from each of five species, showing the PPP score, RefSeq protein accession number, and functional name. All sets of results showed members of the SAO system’s nine core protein families of this system as the top-scoring proteins, although with gaps occurring for those proteins where previous treatment of selenocysteine-encoding TGA codons as stop codons blocked proper structural annotation. The high scores from PPP reflected not only that these proteins were all universal across species with the system (within the limits of structural annotation accuracy at the time the PPP data set was built), but also that strong BLAST score matches to proteins in these families do not occur to proteins in genomes lacking the system.

All proteins with PPP scores higher than 50 (that is, with computed odds of such a high rate of co-occurrence just by chance being 1e-50 or less) from any of the five genomes belongs to the SAO locus. That is, we detected no additional components of the SAO system outside the set of nine proteins families described. High scores were achieved for these because all homologs in the PPP reference set were being found in species with the marker SaoC. Example scores in *C. difficile* include 115.7 for WP_070088344.1 (SaoC itself, hitting 33 YES genomes in the top 33 hits), 107.2 for WP_009897547.1 (SaoB fragment C-terminal to the selenocysteine, with 30 YES genomes out of 30 hit), and 53.3 for WP_004454661.1 (SaoB fragment N-terminal to the selenocysteine. 16 YES genomes from the top 17 hits).

The next tier, all scoring below 40, includes proteins such as WP_084191679.1 from *Tissierella creatinophila*, a glycine reductase complex selenoprotein B (GrdB). The top 148 sequences hit occur in 144 different genomes, including 27 that have the SAO system, for a score of 37.1. Proteins appearing in this tier can include both those with direct connections, as when the end product of one pathway is a substrate for another, and those frequently co-occuring because of sharing biological similarities such niche and ancestry. GrdB may be correlated because 1) its species distribution, though broad, skews strongly toward the taxonomic class *Clostridia*, 2) it functions only in anaerobic environments, just as the SAO system seems to, and 3) it requires a selenium insertion proteins SelA, SelB, and SelD. For similar reasons, proteins associated with selenium-dependent molybdopterin hydroxylases occur in this second tier, including predicted selenium cofactor biosynthesis proteins YqeB and YqeC (6,7).

### Associated Double-Cubane proteins

Supplemental file **SAO_system_PPP_top_hits.xIsx**, in supplemental materials, shows the top 40 hits from PPP for five phylogetically diverse species: *Brachyspira pilosicoli, Clostridioides difficile, Merdimonas faecisa, Syntrophobacter fumaroxidansa*, and *Tissierella creatinophila*. The surprise family of protein at the top of the second tier of PPP scores is the family of 8Fe-9S cofactor-containing “double-cubane” proteins. As can be seen for all five species, at least one of the top five proteins is a double-cubane protein. That includes two species in which it is the top such hit, and two more where it is second. In double-cubane enzymes, two 4Fe-4S iron-sulfur clusters make contact through an additional bridging sulfur atom, creating an unusual 8Fe-9S metallocenter (40,41). The side chains of seven conserved Cys residues, forming the motif 7-Cys motif CX(19)CX(18-19)CX(29)CX(166-170)CX(32-34)CX(32)C, provide ligands to the cluster. These proteins are not known to have a connection to selenium metabolism. 8Fe-9S-type double-cubane proteins are described as having the ability to reduce refractory small molecules such as acetylene and hydrazine (40), but in general, physiologically relevant substrates are not known. Their closest homologs among more completely characterized proteins include beta subunits of enzymes such as (R)-phenyllactyl-CoA dehydratase, (R)-2-hydroxyisocaproyl-CoA dehydratase, and (R)-2-hydroxyglutaryl-CoA dehydratase. These enzymes require an ATP-hydrolyzing activase, an extremely oxygen-sensitive enzyme that transfers a single electron to its target, after which the dehydratase can perform large numbers of turnovers (42).

We built a new HMM, accession number NF040772, to describe double-cubane cluster-containing putative anaerobic reductases. Most were annotated, at the beginning of this study, as “2-hydroxyacyl-CoA dehydratase” or as “2-hydroxyacyl-CoA dehydratase family protein.” Of the 15,511 genomes in the calibration set, just 735 genomes (4.7 %) have a double-cubane protein as detected by hidden Markov model NF040730. But quite remarkably, 11.8 % of all double-cubane proteins occur in the 0.3 % of representative genomes that have the SAO system. Among genomes with at least one double-cubane protein, the count per genome is higher in species with the SAO cassette (2.2 per genome) than in species without (1.2 per genome). Among the 52 species with SAO cassette, 21 (40%) have at least one double-cubane protein encoded within 5000 nucleotides of the cassette, and often more than one. Examples of genomes with double-cubane proteins next to the SAO cassette are *Clostridium botulinum* (see Figure 4), *Clostridium sporogenes, Desulfopila aestuarii, Gottschalkia purinilytica*, and *Syntrophorhabdus aromaticivorans*.

**Figure 4:**
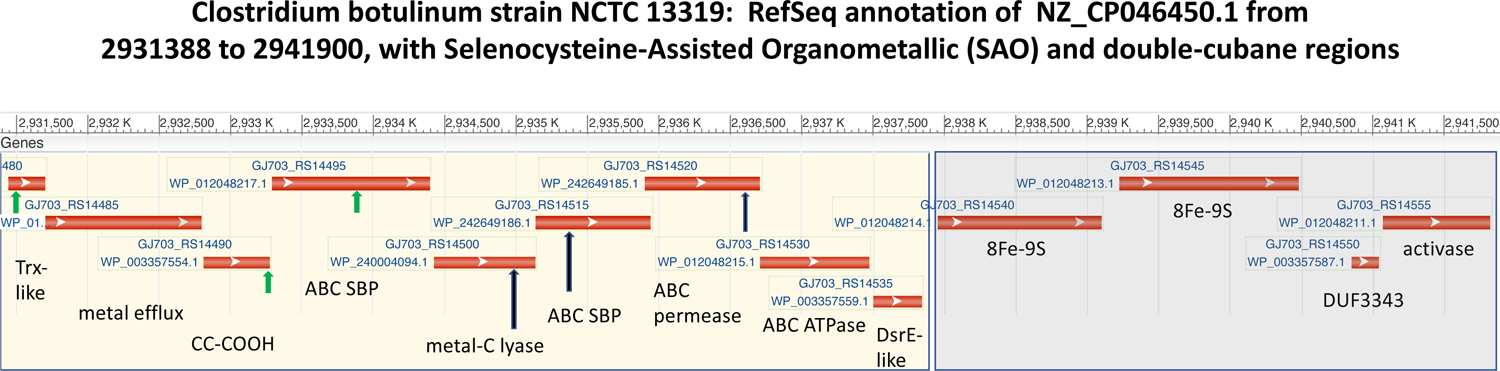
PGAP annotation of the 9-gene *sao* (Selenocysteine-Assisted Organometallic) operon and adjoining double-cubane locus in *Clostridium botulinum* strain NCTC 13319. The region shown is from RefSeq nucleotide sequence NZ_CP046450.1 (2931388-2941900). The SAO (selenocysteine-assisted organometallic) locus (yellow box) gene order is *saoTECXLBPAD*. Black vertical arrows show selenocysteine (Sec) insertion codons in *C. botulinum*. Short green vertical arrows show locations where Sec occurs in orthologs from other species. C. botulinum was selected from among 21 *sao* locus-containing representatives with nearby double cubane proteins and their activase (gray box). A comparable re-annotated *sao* region in *C. difficile*, NZ_JAGIVI010000014.1(60051-66700), has four selenoproteins instead of three, with the genes in a different order, *saoDTECXBPAL* (not shown). The lone 8Fe-9S double-cubane protein in that genome, WP_004454390, is located more than 500,000 base pairs away.

The NF040772 family of double-cubane enzymes includes many examples from genomes that lack the SAO system, including the uncharacterized enzyme YjiM from *Escherichia coli* K-12. However, we were unable to find sequence signatures we could use to separate double-cubane from genomes with the SAO system from their counterparts in genomes without. The comparative genomics therefore seems to suggest the SAO system requires the presence of at least one double-cubane protein, but not the other way around. Substances becoming available in the cytosol as a consequence of the SAO system may enter a pool of compounds that double-cubane proteins can act on, and not be chemically distinguishable from substrates obtained in other ways.

### Double-cubane proteins and selenium

Every species with SAO system has both selenocysteine and one or more members of the NF040772 branch of the family of double-cubane proteins. We therefore looked at the relationship between those double-cubane proteins and selenium. SelD is the selenophosphate biosynthesis enzyme needed for selenocysteine, selenouridine, and selenium-molybdenum cofactor biosynthesis. It is more common in anaerobes than aerobes, and is found in just 38 percent of representative genomes overall. However, 955 of 970 of double-cubane proteins that score > 400 bits to hidden Markov model NF040772 are encoded in SelD-containing assemblies. Of the 15 exceptions, three are from bacteria that encode selenocysteine biosynthesis protein SelA, suggesting that SelD’s apparent absence may be artifactual, and nine others are from methanogenic archaea. Of over 900 examples of bacterial double-cubane proteins identified in representative assemblies, only three are from assemblies lacking SelD.

This remarkably close association of the NF040772 and SelD families could be explained by a role for double-cubane enzyme in processing selenium-containing compounds, or in pathways that have obligatory selenium-dependent steps either before or after. Explanations of this kind for the double-cubane enzyme to selenium linkage are plausible and may be seen as more likely than a previously unsuspected novel chemistry. But we list as a formal possibility the explanation that the structure of the cofactor itself is what links the selenium donor protein SelD to the NF040772 family of double-cubane proteins. The correlation we detect may signify biosynthesis of selenium-bridged analogs of the recently described 8Fe-9S cofactor in a subset of double-cubane proteins. If so, then the active site cluster in double-cubane proteins and selenocysteine sidechains in SAO system proteins may share abilities to bind a similar class of compound through a selenium atom. The product or an intermediate of one system may be a substrate acted upon by the other.

### Revisiting methylmercury formation proteins HgcA and HgcB

Proteins DND132_1056 (EGB14269.1) and DND132_1057 (EGB14270.1) in *Desulfovibrio desulfuricans* ND132, and similar pair GSU1440 (AAR34814.1) and GSU1441 in *Geobacter sulfurreducens* PCA, were described in by Parks, et al. in 2013 as proteins HgcA and HgcB of a two protein cluster needed for methylmercury formation (43).

Unlike DND132_1056 and GSU1440, the founding members of HgcA family described by Parks, et al., occasional homologs differ by having N-terminal extensions that include CC motifs. One such proteins was examined in a site-directed mutagenesis study (44). We examined the coding capacity of 5’-extended versions of the coding region for DND132_1056, GSU1440, and 18 additional homologs of similar length. In all 20 cases, we found that the coding region could be extended upstream, to a new length between 363 and 400 amino acids, by allowing translation of a single UGA codon as selenocysteine. The minimal required N-terminal extensions are MAPPDVPARDDAPC**U**GPRPDPRAGVFEKPGYAVEPFV for DND132_1056, and MSQPGPDVDRPPC**U**GPKTPAVSGTINENVPGFLRWLDTPAGR for GSU1440. Across all 20 proteins, the pentapeptide motif surrounding the selenocysteine residue is perfectly conserved, PCUGP, and all species encode the proteins required for selenocysteine incorporation. The CU motif resembles that from several selenoproteins of the SAO system, and lends further support to the notion that the SAO might be involved in organometallic compound formation, as well as dismutation.

Examination of HgcB showed that it too contains selenoproteins, 17 of 266 examples (6.4%). While the HgcB, like SaoC, has a Cys-Cys dipeptide as the final two residues for nearly every family member, the site of selenocysteine incorporation in HgcB differs, occurring instead about 25 amino acids from the C-terminus, at residue 73, the 9^th^ of HgcB’s invariant Cys residue positions.

**Figure 5** shows the similarity CU motifs found in this study for in several key proteins of the SAO locus and in methylmercury formation protein HgcA. The CU motif is relatively rare in selenoproteins. We see it in the uncharacterized CUAEP/CCAEP-tail radical SAM protein (NF040546) and at the C-terminus of the Campylobacter/Helicobacter lineage KCU-star (NF033934), both of whose structures and functions are not well known. It occurs in a small group of mercuric transporter MerT family selenoproteins such as WP_250634603.1, and in redoxin domain-containing proteins in the family related to WP_012872293.1, but in no other selenoprotein we know of outside of the SAO and HgcAB systems. We find the three examples in SAO system remarkable, and the sharing of the CU motif with HgcA proteins to be highly suggestive of a shared interaction with a heavy metal atom. The CU in the SAO system occurs in protein exposed to the extracellular milieu (SaoC and SaoB), but also in the cytosol (SaoL). We can add to this SaoT, a thioredoxin-like protein, which has two different forms of selenocysteine-containing motifs, UXXC and CXUC, in addition to two non-selenoprotein forms, and cysteine-overload CXCC form (extremely rare in other thioredoxin like proteins) and the CXXC form expected in thioredoxins. It seems quite likely that the multiple selenoproteins of the SAO system, or their Cys-overloaded non-selenoprotein homologs, bind a mercury-like heavy metal-containing part of some compound, or class of compound, as it is trafficked from outside the cell to inside, most likely undergoing covalent changes in the process. While possible roles for the SAO system could include making mercury less toxic for itself, or more toxic for competing organisms, we speculate that an alternative role could be the catalytic use of a trace element present in extremely small quantities for sa new type of selenium-assisted energy metabolism.

**Figure 5:**
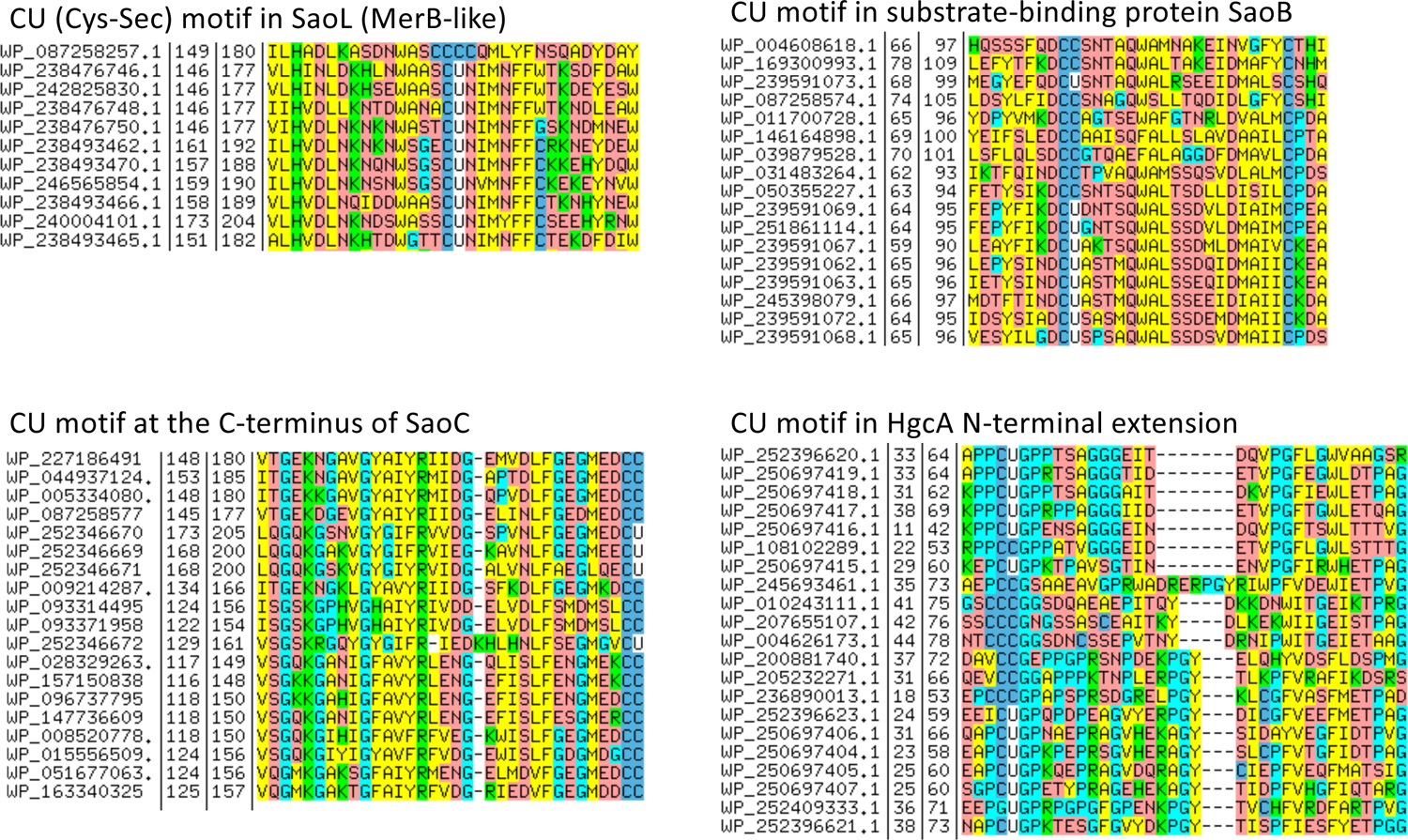
Alignments of selenoprotein and non-selenoprotein versions of proteins SaoL (top left), SaoB (top right), SaoC (bottom left), and HgcA (bottom right), for short regions that include the site where the selenocysteine may occur, are show as displayed in belvu (16), colored by residue type. For SaoL, homolog of the organomercurial lysase MerB, the top sequence (WP_087258257.1) was exceptional for having Cys (C) rather than Sec (U), and the site is “packed”, with four consecutive Cys residues. Similar packing with extra Cys residues is seen in methyl mercury formation protein HgcA region. Full-length multiple sequence alignments for all eight selenoprotein families described in this paper can be found in Supplemental Materials.

### Rolling reannotation of RefSeq genomes

RefSeq regularly subjects prokaryotic assemblies to reannotation by the PGAP prokaryotic genome annotation pipeline (19) so that both structural and functional annotation can be kept up-to-date. Reference data that enables proper prediction of all novel selenoproteins described in this paper has been added to PGAP, so the rolling reannotation is replacing fragmented, incomplete, or missing selenoproteins with corrected versions containing U, the single letter code for selenocysteine, in the amino acid sequence. The exception is SaoC, as PGAP currently cannot correctly predict C-terminal selenocysteine residues. Sequences belonging to any of the HMM-defined protein families listed in Table 1 and currently annotated on any RefSeq genome can be retrieved by query with the HMM’s accession number at https://www.ncbi.nlm.nih.gov/protfam. Genomic regions that now have correctly annotated selenoproteins can be visualized, such as https://www.ncbi.nlm.nih.gov/nuccore/NZ_CP046450.1?report=graph&from=2931388&to=2941900, as seen in Figure 4, or https://www.ncbi.nlm.nih.gov/nuccore/NZ_JAGIVI010000014.1?report=graph&from=60051&to=66700&strand=true for a *Clostridioides difficile* strain JV65 033.

Figures S1 through S8 in supplemental materials show full-length or minimally trimmed alignments for the eight selenoprotein-containing families discussed in this paper. Sequences in each alignment were made non-redundant by removing the shorter protein from any pair above a certain percent identity. These thresholds were 80% for SaoL (Figure S1), SaoB (Figure S2), SaoP (figure S3), and SaoT (figure S4). The thresholds used were 85% for SaoX (Figure S5), 95% for SaoC (figure S6), and 90% for HgaB (figure S8), families in which selenoproteins are relatively rare. The alignment for HgcA (figure S7) shows selected sequences from the family, non-redundant to below 75% identity, with some manual editing to correct small errors made during progressive alignment by Clustal W. Alignments for SaoL, SaoB, SaoP, SaoT, HgcB, but not SaoC, SaoX, or HgcA, include sequences that have not yet reached by RefSeq’s rolling reannotation and that therefore appear in truncated form, either N-terminally or C-terminally, caused by improper treatment of the selenocysteine-encoding UGA codon as a stop codon.

## Discussion

### The SAO system contains six novel selenoprotein families

We have described here a rare but taxonomically diverse conserved gene neighborhood that is exceptional in a number of ways. In a collection of representative prokaryotic genomes, one per species for 15,511 species, the system occurs in just 52 genomes, broadly distributed taxonomically, but focused on anaerobes that show various adaptations for performing biochemically challenging fermentations in anaerobic environments. The system is a nine-gene operon, encoding four previously unknown protein families that are selenocysteine-containing a majority of the time (98%, 85%, 65%, and 60% for SaoL, SaoP, SaoT, and SaoB, respectively). Two additional families contain small numbers of selenoproteins, bringing the set of novel selenoproteins up to six new families.

### CC and CU motifs suggest interaction with heavy metal atoms

The putative organometallic lyase SaoL has a CU motif that aligns to a mercury-binding Cys at MerB’s active site. The putative metal efflux transporter SaoE has a CSC motif that aligns to the motif USC in selenoprotein homologs in the SO_0444 family of Cu/Zn efflux transporters. Additional signature motifs in the SAO system include CU in the predicted substrate-binding pocket of SaoB, CC in the substrate-binding cleft of SaoX, U at the gating position of the homodimer-forming permease SaoP, CXUC and variations at the redox disulfide bond-like site in SaoT, and a C-terminal CC motif that sometimes is CU in the dark matter protein SaoC. This is a remarkable repetition of the selenocysteine or double-cysteine theme across the SAO, within each system suggesting multiple proteins interacting with the same metal or its adducts at multiple cellular sites.

### Selenocysteine is unknown in characterized mercury resistance protein families, but is discovered in methylmercury formation

Our previous extensive efforts to catalog authenticated prokaryotic selenoproteins as structural reference sequences for NCBI’s PGAP prokaryotic genome annotation pipeline resulted in no examples of a selenoprotein involved in mercury resistance. This made us curious about the need for selenocysteine in the MerB homolog SaoL and in the rest of the SAO operon. Our suspicion that the system might catalyze for the formation and the dismutation of organometallic compounds sparked our interest in a previously described pair of proteins known to participate in organomercurial compound formation, HgcA and HgcB. Examination of the HgcA and HgcB proteins revealed that both can be selenoproteins, HgcA in the great majority of cases, with a previously unknown N-terminal extension, and HgcB occasionally. The motifs are CU in the newly discovered N-terminal extension of HgcA selenoproteins for methylmercury formation, and CC at the C-terminus of HgcB, echoing what is seen in the SAO system. We found these strong parallels between the SAO and HgcAB systems, in the use of selenocysteine and the prevalence of CU and CC motifs, only after we started to hypothesize that the SAO system could form organometallic compounds as pathway intermediates, and consider our observations about selenocysteine’s role in methylmercury formation to be significant support.

### Co-occurring proteins suggest adaptation to anaerobic environments by selenocysteine-enabled fermentation of normally refractory substrates

We described above a special relationship between the SAO system and 8Fe-9S cofactor-containing double-cubane proteins, showing that double-cubane proteins are universally present in SAO-containing genomes, extra abundant in those genomes, and encoded adjacent to SAO proteins in almost half of those genomes. Correlations with *grd* (Glycine Reductase) genes are similarly informative. These genes of Stickland metabolism include grdB (glycine reductase complex selenoprotein B), grdH (betaine reductase selenoprotein B), and grdF (sarcosine reductase complex component B subunit beta) all are selenoproteins that help enable anaerobic fermentations most bacteria cannot do. In Stickland fermentations, oxidation of one amino acid is coupled to reduction of another (or of sarcosine or betaine). The reductive metabolism leads to ATP formation (45). SAO operons are present in 0.3% of bacterial species, but in 13.4% of those with GrdX family proteins. GrdX is present in just 2.2% of genomes, but in 87% of those with SAO. Both SAO and Stickland metabolism require selenocysteine, but selenoprotein families involved in processes other than Stickland metabolism do not show such strong patterns of co-occurrence with SAO. For example, Clostridium difficile has a selenocysteine-containing formate dehydrogenase, but its Partial Phylogenetic Profiling score does not place it in the 1500 proteins best correlated with the SAO system.

### The SAO system’s conservation, context, and patterns of design suggest metabolism, not resistance

The SAO system is sporadically distributed, as are the markers of Stickland fermentation, found in bacteria that are taxonomically highly divergent, while occupying similarly anaerobic environments. Examples of the operon we have viewed in extended local context show no signs of inhabiting mobile islands now or having undergone recent horizontal gene transfer. All genomic contexts we viewed showed housekeeping-type genes as neighbors rather than apparent markers of mobility such as transposases or phage or plasmid proteins.

Conservation within lineages similarly suggests a central metabolic role. The *C. difficile* genome is known to be heavily invaded by mobile elements and mosaic in structure (46), so substantial variability of genome content is expected from one strain to another. Genes that help confer resistance to trace metals such as mercury or arsenic, such as *merA*, typically frequently appear in plasmids, or in genomic islands flanked by markers of mobility. We find *merA*, encoding mercury(II) reductase, in 5469 of 34051 RefSeq assemblies (16%) of *Escherichia coli*. By contrast, we find the SAO system in *C. difficile* in 3279 of 3343 assemblies (98%). All genomic contexts we viewed showed housekeeping-type genes as neighbors rather than apparent markers of mobility such as transposases or phage or plasmid proteins.

The SAO system’s *pattern of design* (47) seems to argue against a resistance function, and therefore to further support a metabolic role such as energy metabolism. The system occurs exclusively in bacteria that grow anaerobically, and mostly in members of the Firmicutes, bacteria that lack an outer membrane. The MerT and MerP proteins in Gram-negative transport the highly toxic mercuric ion, Hg(II), from the periplasmic space to the cytosol, where MerA can convert it to the less toxic Hg atom. But for species that lack an outer membrane to enclose a periplasmic space, it seems an unfavorable design to import mercury ions or compounds from the whole of the extracellular milieu in order to detoxify them, imposing energy costs for substrate import, metal efflux, and expression of nine different proteins. Similarly, relying on selenoproteins seems like a strange design with and inherent liability if the purpose of the SAO system is mercury resistance. A lack of selenium would block protein translation and prevent the function of proteins needed for resistance if selenium is in short supply. Having such limitation is plausible, of course, but we have encountered no examples.

### Catalysis vs. binding affinity

Selenocysteine in prokaryotes is well known for replacing Cys in enzymes at the active site, giving the enzyme dramatically higher catalytic efficiency than its Cys analog would have. Selenocysteine is far less common in proteins without enzymatic or redox function. But selenoproteins SaoB and SaoP are binding and permease subunits of an ABC transporter, normally expected to have a binding but not a catalytic role. Their selenocysteine sidechains raise the prospect of a dual role for the transporter. Selenium atoms in contact with the transporter’s substrate might effect some covalent change. Two different SBPs work together with a single permease that has a pair of active site-like selenocysteine sidechains meeting at the crystallography-defined substrate-gating site. Any number of different scenarios could be proposed in which the SAO system’s ABC transporter assembles a substrate molecule before transporting it. Catalysis by transporters has ample precedent, including sugar import tied to phosphorylation in PTS transporters. On the other hand, there is an extraordinarily high binding affinity between selenium and mercury (48). Multiple steps in the handling of a single substrate found in the extracellular milieu, involving binding rather than catalysis, could be the reason so many selenoproteins are found in the SAO system.

### Is mercury itself the most likely metal for the SAO system to traffic?

Multiple lines of evidence suggest the binding to, transport of, and likely the formation and/or dismutation of organometallic compounds. But is not clear that mercury is the only or even principal metal involved. MerB family lyases are known to accommodate metals other than mercury at the mercury-binding site, even if physiological substrates other than organomercurial compounds are not known. The CU motif at the heavy metal binding site of SaoL is non-canonical for the MerB family, so it cannot be assumed automatically that organomercurials are appropriate substrates.

Most components of the SAO system are invariably present, but SaoL itself occasionally is not detected, missing in 9 of 52 loci. How the system could include in most cases such a vividly suggestive component, yet dispense with it in some bacteria, is unclear. But these variant forms of the system may provide some clues to the function of the system. *Desulfopila aestuarii* has a replacement for SaoL, an apparently authentic MerB protein, WP_073613903, encoded in the middle of the SAO operon. This replacement protein has much higher levels of sequence identity to homologs found in the middle of mercury resistance loci than it does to any SaoL protein.

Our best guess for the reason that SaoL can be dispensable is that the system can act on a variety of metals, and the system may not require Hg to be present. We have suggested that organometallic compounds are formed for transient existence, as they would be consumed later in a pathway that is part of fermentation for energy generation. The organic portion of the molecule may undergo redox changes, with the attached metal atom facilitating otherwise biochemically difficult transformations. The final step, liberating the metal moiety from the fully reduced end product, might proceed non-enzymatically at a sufficient rate, for metals other than mercury, that SaoL is not required.

### Model for the biological process carried by proteins of the SAO system

We suggest that the selenoprotein-loaded SAO system provides a novel mechanism for *C. difficile*, and other bacteria that share the same rare operon, to make use of certain normally refractory organic substrates in organically rich but oxidant-poor environments. More specifically, we suggest that the SAO system manages the import of a food source that most bacteria cannot readily utilize in anaerobic conditions, and furthermore, that at some point during the process of import and catabolism, enzymes and/or transporters are acting on an organometallic compound. Said organometallic compound might typically have a short half-life in solution, but is chaperoned before, during, and after transport by selenoproteins that have multiple Cys or Sec residues in contact with the metal atom. Following import, the compound is reduced by an 8Fe-9S (or possibly a 2(4Fe-4S)-Se) cluster-containing double-cubane protein, making the compound an oxidant that enables ATP generation by other cellular machinery, such as the energy-transducing Rnf complex (45). Once the compound can no longer be reduced further, the carbon-metal bond may be cleaved by SaoL, chaperoned to the efflux transporter SaoE, and extruded. It is further possible that the metal, when extruded, is captured for reuse in the next turn of the cycle. In this scenario, even trace amounts of a heavy metal with the right properties could serve a catalytic role to provide a competitive advantage in energy metabolism to bacteria with the SAO system.

Although our model is admittedly highly speculative, it is built on a number of potentially important underlying observations about various protein families and their unexpected structures and relationships. Quite a number of experiments to confirm or falsify part of the model immediately suggest themselves, including tests for uptake of and sensitivity to mercury and other heavy metals, ability to metabolize various compounds with or without certain metals present, ability to form or cleave various organomercurial compounds, and how these results change if certain genes within the SAO system are knocked out.

Our discovery of eight novel selenoprotein families overall, six in the SAO system and two in methylmercury biosynthesis, suggests there may be many others left to find, with potential importance in understanding conversions of heavy metals to more or less toxic forms in gut or environmental microbiomes, or for bioengineering applications. We suggest that selenocysteine side chains may potentiate organometallic compound biosynthesis in a wide variety of individually rare pathways that remain to be discovered. Their relative rarity, plus the likely scarcity of non-selenoprotein homologs to assist with their discovery, has likely made such systems difficult to discover in the past. However, the potential for finding more such systems now seems clear. Families of novel organometallics may have transient existences in anaerobic metabolism as pathway intermediates, including in gut microbiomes, undergoing formation, transport, transformation, and dismutation, all in pathways that remain to be discovered, in part because key markers of those systems are still-undiscovered selenoproteins.

This research was supported by the Intramural Research Program of the National Library of Medicine at the NIH. This work utilized the computational resources of the NIH HPC Biowulf cluster (http://hpc.nih.gov). The authors would like to thank Eric Boyd and Ian Krout for helpful discussions.

