## Supplemental Figure 4 for "Eight Unexpected Selenoprotein Families in ABC transport, in Organometallic Biochemistry in *Clostridium difficile* and other anaerobes, and in Methylmercury Biosynthesis"

thioredoxin-like protein SaoT (with proteins truncated at selenocysteine UGA codons)

```
WP_003483328.1   1 MKKILVEFVNTCPCCDEYSEFVKEVCSKYSDEVECKIYYAGKDIDYIKKYGMILKGTLIVNEKKKIDKLSKEIIESEIEKAIC
WP_206170201.1   1 ---------------------MQAAAAKYKDKVEARIYYAGKDITYIKKYGPIFKATLIINEKKKIQRISKDIIEKEIEMAVI
WP_246579167.1   1 MSKVLVEFINTUPSCDGYTRAVEVAAAKYKDKVDVKIYIAGKDMSYVKKYGVIFRGTLIINEKKKIQRISKNIIENEIALAVS
WP_051590281.1   1 -------------------MLVKKIGDKYIDKVEVKLYQAGKDFSYIKKYGMITKGTLIINQKKKYDRLSKEIIEKAILEAII
WP_242825833.1   1 MAKVLIEFISTUPSCDEHGGLIKDLGKKYSDKLEVKLYTAGKDFSYVKKYGLITKGTMIIGQKKKYDRLNKKSIEKAILEAIC
WP_240067708.1   1 MAKVLVEFINTUPTCDEHGMLIKRLGEVYKEKLDIKLYQAGKDFTYLKKYGIITKGTMIINQRKKYDRLSKDVIEKAIIEAIN
WP_243159878.1   1 MSKILVEFINTUPSCDEHGVFVKKMSEKYADKLEIKLYQAGKDFSYIKKYGIITKGTLIINQKKKYDRLNKDTIERAIVEAIN
WP_008520781.1   1 MTK--VEFINTCPCCDGYGDVLRDLQKRFPDKIDLKIYYVGKDFDYLPKYGPITRGTMIINGTDRYEDLSRRSIEKIISEAIC
WP_013244969.1   1 MSKVLVEFINTCPTCDGYSSYLKELQETYKDKIEVKLYYAGKDFDYLAKYGMVDRGTLIINEKNRYDVLSKKLIEQEIKKAII
WP_147736611.1   1 MSKILVEFINTCPTCDSYSEYLNELQKTYEDKMELRIYYAGKDFDYIQKYGIIDRGTLIINGKNRYEILNKKLIEKEIKAAII
WP_011700724.1   1 MSKVKVEFINTCSCCEEYGRIIRSVASGHGDKVEVKIYIAGKDFDYLKKYGPVTRGTMIINGEKRLDTLSREIIEKAVTDALA
WP_190239395.1   1 MSKTLVEFINTCSCCDEHIQTIKNTAQKHGDQVEVKIYYAGKDVDYLKKYGMVTKGTMIIDGRKKIDNLSKTIIEKAIEDALI
WP_012031740.1   1 MARVQVEFINTCPCCEEHVINIKKAAARYGDEVVVKIYHAGKDFDYIKKYGAVSKGTMIINGRKKLDNLSRSIIEKAIEEAVI
WP_243153044.1   1 MAKVQVEFINTUPSCDEYARVIRHAAAKYGDVVEVFIYYAGKDMGYLKKYGMVTRGTMIVNSRKKYDILSRAVIEKAIEEAVI
WP_028893773.1   1 MAKVKVEFINTCSSCVEIGHTIQDVAAGYGDDVDVKLYVAGRDFDYLKKYGMVCKGTMIINGKTKYDNLNREIIEKAIDKAVI
WP_073613904.1   1 MSKVTVEFINTCACCDEHGESIKEIAAKYGDAVDVSLYYAGKDFDYLKKYGMISKGTMIINGTRKYETLSKDIIEKAIGEAMQ
WP_107740078.1   1 MAKVQIEFINTCPCCDEHGATIQDIAARYGDDIDVFIYNAGKDFGYLKKYGMISKGTMIINGVKRYENLSKEIIEQAISQAMA
WP_073089016.1   1 MGKVEIEFINTCSCCASYEEMIKKAAAPYKDQVNLKFYAAGKDMDYIRKYGMVSKGTMIINGKKKYDQLNQEIIERAIKEAVI
WP_093371960.1   1 MAKVLVEFINTCPTCVGYEEMIRKAAKEKGDQVEVKIYYAGKDYDYVRKYGMVTKGTMIINEKKKYDRLNQKTIEDAISNALI
WP_031483268.1   1 MSKTLVEFINTCPCCDEYVNIIEKTTSQYGSSIDLKVYYAGKDFDYVRKYGQVNKGTMIINERKKYDSLTTTVIENAIAEAVA
WP_163340323.1   1 MVS--VEFINTCPCCDEYVTIIRDYIKSKEGKVSLKVYYAGKDFDYIRKYGPVNKGTMIIDGKKKYDSLTTPVIHNAIDEAMS
WP_005542134.1   1 --------------------MVQSSAERYGDKIELKMYRTGKDMDYIRKYGMIYVGTMIINEKIFIKNLNRKSIDKAIDDAIY
WP_068474708.1   1 -----------------MGRNVKDVAKEFGDKVEVKLYSQGKDIEALKKYGMVFQGTMIINEKKKVTRLSVKNIRKEIAQAVI
WP_114642955.1   1 ---------------------MKDYAKKKG--ITAKIYKAGKDFGYLKKYGAVMKSILIINETKKYQTLSEEIIKKAIDEAV-
WP_240067710.1   1 MVNILVEFVNTCAUCNKYAVFVEDEVKKYNGKVKLKIYRVGKDFDYIKKYGMVTKSMIVIDEKKKIQNLSEGVISRAIEEAVI
WP_242842469.1   1 MGKILVEFINTCAUCDKYGAFVESLAKQHEDKLELVIYRAGKDFGYIKKYGMVTKSMIVIGEKKKIQNLSEKSITEAIEEAVI
WP_242943826.1   1 MEKVLVEFINTCAUCDHFGGFVEDLEIKYPGKVEVVIYRAGKDFGYIKKYGMVTRSMIVINERTKVQNLSEASITKAVEEAVM
WP_252342676.1   1 -MFVFVEFINACAUCNKYEGFVKALSDEFDN-VKVKVYNVGKDFDYIKKYGMITKSTVIINEEKRLNDLSETNILNAVKEEVI
WP_009214285.1   1 MKKPCLEFILACACCAELAKYTEELAKDYP-EVETKVYTAGKDTDYIEKYGALTKSLLVVNEEKAYRRLSRAVVKEAFEEALI
WP_072832008.1   1 --------------------MVQELAKKYEDLVEVKVYKAGKDFDYIRKYGLMSKSVLIIDESQVVENLNKKVIELAFSQLMC
WP_240067711.1   1 MKKATLEFINACAUCNVYTSLVEGLGEEYRDKVQVKIYKVGKDFEYIRKYGPVMKSILIINEKKKIENVNRDTIRKAFQEAAI
WP_242948466.1   1 MKKGFLEFINACAUCNVYTSFVEGLAKEYRDRVNVKIYKVGKDFDYIKKYGPVMKSILIINERKKIDNISKETIRKAFEEAII
WP_242838409.1   1 MKKNLIEFINVCAUCDTLTPFVEELGKKYKDHVDVKIYKAGKDFDYIRKYGMFNKSVLIVDEKKVIDNVNKSTVEKTFMSIVC
WP_242942359.1   1 MQKPLLEFINVCSUCDIFGSLVKTLGERYKDKVDVKIYKAGKDFEYVKKYGMFTKSVLVLNETKIIDKVNKATIAEAFKELAC
WP_087258584.1   1 --------------------MVQSLAVEYRGRVDVKIYKSGVDTEYVAKYGAISKSMLIINEAKAIAKLSKSAVRNAFEEALI
WP_089610263.1   1 ---------------------MKSLAEQYQGQIKVTIYKVGEDFDYIKKYGPTTKSMLIINESRVVTKLSKESIRKAFEEVLI
WP_119205921.1   1 ------------------MGFVESLRKEYEGRLEVVIYKTGKDFDYIKKYGAVTKSMLVINESKAIKKLSRESITAAFEEAIC
WP_240067709.1   1 MKKPCVEYICACAUCEVYVGFVESLKKEYEGRLEVVIYKAGKDFGYIKKYGATTKSMLVINESKAVRNINKETIVAAFEEAIN
WP_252200387.1   1 MEKPSIEYISACAUCEVFEGLVQALEKEYEGKVHVKIYKAGKDFDYIPKYGAVTKSMVVINEKKAVTKLTKPAIRKAFEEALI
```
