## Supplemental Figure 8 for "Eight Unexpected Selenoprotein Families in ABC transport, in Organometallic Biochemistry in *Clostridium difficile* and other anaerobes, and in Methylmercury Biosynthesis"

mercury methylation ferredoxin HgcB (with proteins truncated at selenocysteine UGA codons)

```
WP_250697400.1     1 --MKNLRYLDDVSSLVLDRDLCIGCGLCTQVCPQEVFEMEE---NKASIVDFNACMECGACVNNCPSGALQVSP--GVGUASYIIQVWIK
WP_031449992.1     1 --MKELRYLDDVSTLVLNKEKCIGCSLCTEVCPHAVFEMRH---NKARIVDFNACMECGACVNNCPSEAISVSP--GVG-----------
WP_092232719.1     1 --MKELRYLDDVSTLALNDEKCIGCSLCTQVCPHAVFKMRN---DKAHIVDFNACMECGACVNNCPSGAIAVSP--GVG-----------
WP_250697390.1     1 --MQDFRYLEDVATLVLDEQKCIGCGLCTEVCPHAVFEIQE---KKAQIVDFNACMECGACVSNCPADAISVSP--GVGUATYIIQVWIK
WP_250697391.1     1 --MRDLRYIEDVSTLVIDQEKCIGCSLCTVVCPHGVFEMKE---RKAVIVDLDGCMECGACVNNCPVYALNVTP--GVGUAAYIIQVWIK
WP_250697393.1     1 --MQNLRYLDGVSTLKLSQKLCVGCGLCIEVCPHGVFVLDE---HKAHIQDLDGCMECGACVNNCPTSAISVTP--GVGUAAYIIQVWIK
WP_045674113.1     1 --MKTLRYLDNVVTLTLNAQKCIGCGACTQVCPHTVLAMQS---RKAYILDMNACMECGACANNCPTEALTVTP--GVGCASYIIQSWIV
WP_092212500.1     1 --MKDLRYIDNMATLELHTDLCVGCGLCTTVCPHGVFAQEG---PKALIADGNGCMECGACARNCPVAAISVNP--GVGCAAYIISGWIK
WP_236890014.1     1 --MKDLRYIDNMATLTLHSDLCVGCGLCTTVCPHGVFAKEG---PKVRIDDLDGCMECGACARNCPVAAISVDP--GVGCAAYIISGWIK
WP_155317485.1     1 --MKRFRYLPGVTTLEYDKGLCVGCGSCERVCPHGVFTMNG---KKVQLTDKDGCMECGACALNCPTSAIRVTP--GVGCAAYIISGWIH
WP_213183181.1     1 --MRDLRYLRGVTTLAYENERCIGCGNCILVCPHSVFARDGE--RKVRLADPDGCMECGACALNCPTEAIRVTP--GVGCAAYIISGWIH
WP_144305632.1     1 --MKNFRHYSGVATLKLDRDACVGCGLCPQLCPHGVLAMEQ---KRARIVDPDGCMECGACAKNCPSGALSVSP--GVGCASYIINKWLR
WP_144301671.1     1 --MRDFRHYDGVATLNLNEDACVGCGLCPQLCPHGVLAMDG---RKAHIVDLDGCMECGACARNCPAQALSVKP--GVGCAAYIIQVWLR
WP_163302676.1     1 --MRDMRYLSGVASISLDVRRCVGCGLCEAVCPHGVLALTE---GKAGIVDRDGCMECGACVTNCPTGAVAVTP--GVGCAAYILQVWLK
WP_176630762.1     1 --MRGMRYLDGVATIGLDRERCVGCGMCAVVCPRGVLAMDG---GRAVLVDRDGCMECGACVTNCPSGAVSVTP--GVGCAAYILQVWLR
WP_027186353.1     1 --MNDFRYLDNVSTLVLDPAACVGCGMCEIVCPHGVLRVED---RMAVIVDHDACMECGACALNCPTHALTVTP--GVGCASYLIQTWFK
WP_037564333.1     1 --MESFRYLSGVTTLQLDPEACVGCGMCENVCPHGVFLLEG---KKAVVRDIDGCMECGACALNCPTDAIHVTP--GVGCASYIIQTWIK
WP_126379058.1     1 --MKDFRYLDGVATLKYDAEACVGCGSCTTVCPHGVFELQG---KKAVIVDGDGCMECGACAMNCAPGAITVTP--GVGCAAYIIQTWLP
WP_022661037.1     1 --MKTFRYLEGVVTLALDPERCIGCGACGTVCPHGVLRLEGE-PRRAHIADRDGCIECGACAVNCPTSAVSVTP--GVGCAALILASWLG
WP_158950321.1     1 --MKDFRYIDGVSTLTLDRDTCVGCGLCTEVCPQRVLRVRE---GKAEIIDFDACMECGACVTNCPVSALSVHP--GVGCASHILSVWLS
WP_066802544.1     1 --MKDFRYLDKVATLKLDTDTCIGCGACAQVCPHRILEVRD---KKAAILDFDACMECGACWRNCPVEAITVTP--GVGCASYLISVWLH
WP_229594619.1     1 --MKDFRYIDNIATLKLDEATCIGCGMCAVVCPHRILEMRE---KKAVILDFNACMECGACAKNCPVEAITVTP--GVGCASYLISVWLY
WP_071544227.1     1 --MKNFRYLDGVATLALDTDACVGCGSCVEVCPHRILEMRG---KKAAILDPDACMECGACATNCPVRAVTVTP--GVGCASYLISVWLH
WP_014321698.1     1 --MKDFRYIDGVSSLALDTDKCVGCGSCVDVCPHRILAVRE---RKTTILDFDACMECGACARNCPVEAITVTP--GTGCAAYLVSVWLH
WP_128329673.1     1 --MKEFRYIDDISTLKIDTGKCIGCGLCVTVCPHRVLTTQD---KKADVVDFNACMECGACAQNCPTAAITVTP--GVGCASYIISLWLH
WP_013515565.1     1 --MTEYRYINGVATLKLDQDACVGCGMCATVCPHRVFAVSE---RKAGIVDFDACMECGACARNCPASAIAVRP--GVGCASYIMAVWLA
WP_020875938.1     1 --MKDFRYIGGVATLELDTGRCVGCGMCETVCPHRIFTVRE---KQAVIRDIDACMECGACARNCPTKAIAVTP--GVGCAAYIISTWIH
WP_097012721.1     1 --MKSFRYIPDVATLKLNRDECVGCGMCHAVCPHRIFQLRE---KKAEIVDLDACIECGACAKNCPTAAISVNP--GVGCASLIMSIWWH
WP_045222548.1     1 --MKRFHHLQNVATLAYDRDKCVGCGLCVAVCPHRVFVLKKD---KADIQDREACMECGACALNCPTEAVAVTP--GVGCASYIIQTWLP
WP_031386617.1     1 --MKRFHHLQNVATLTYDRDKCVGCGLCVTVCPHRVFTLQND---KAHVQDREACMECGACALNCPTQAVIVTP--GVGCASYIIQTWLP
WP_028570972.1     1 --MKQFRHLQNVATLAYDQEKCVGCGLCVAVCPHRVFTLKND---KAHVQDREACMECGACALNCPTEAMSVTP--GVGCAAYIIQTWLP
WP_205240453.1     1 --MKRYRHLENVATLAYDREKCVGCGLCATVCPHRIFAVRDG---KAEVLDREACMECGACALNCPTEAVTVTP--GVGCAQYIIQTWLP
WP_012805587.1     1 --MKIFRHLKNVATLAYEEDKCVGCGICATVCPHRIFAVRNG---KAHVLDREACMECGACALNCPTAAITVTP--GVGCAQYIIQTWLP
WP_092118529.1     1 --MKRYRHLQNVATLLYNEDKCVGCGLCATVCPHRIFVLCNG---KAAVMDREACMECGACALNCPTEAITVTP--GVGCASYIIQTWLP
WP_054649311.1     1 --MKEFRYIEGKATLEYDRDKCVGCGMCAAVCPHGVFIMNER---KAEVVDHGACMECGACSMNCPVEAVSVTP--GVGCAALIIRRWLG
WP_018125557.1     1 --MKDFRYLEGMATLSLDTQKCVGCGMCLAVCPHEVFRLEDG---KVVLADHGACMECGACALNCPVDAVSVTP--GVGCASLIIKRWLG
WP_205232270.1     1 --MKGLRYIENVASLTLDKDRCVGCSTCVTVCPHRVFQLQES---KAVMVDKNGCMECGACAVNCPVDAIRVDP--GVGCASLIINRWLK
WP_174408776.1     1 --MQGFRYIENVVTIGLDAEKCVGCGLCTEVCPHLVFRMQDG---KAGFADREACMECGACARNCPVGALSVNP--GVGCASLMIKRWLH
WP_174404739.1     1 --MKDFRYIDGVASIALEAKTCVGCGVCVQVCPHQVFTVQKG---KAVFADRNACIECGACAKNCPVDAILVTP--GVGCASLMIKRWLH
WP_230996100.1     1 --MKGFRYLDGVATVRLDPSACVGCTRCVEVCPHQVFAVSD---RKAHIVDKDACMECGACAKNCPTAAIEVEA--GVGCATWVIDRWFG
WP_175641722.1     1 --MKEFRYLPDVTTLKLDRELCIGCGICHLVCPHGVFEPDG---NVMEIVDLNGCMECGACSHNCPVDAISVNP--GVGCASLIINTWLK
WP_163335649.1     1 --MINFRYIDNTATLKLDVGRCIGCGNCLTVCPHRIFELVD---KKAVIQDLDACMECGACSVNCPVEAISVKP--GVGCAEAYINLWVE
WP_084813749.1     1 --MQTMRYIDNTTTLKLNVDVCVGCGNCILVCPHRVFEIKQ---KKATICDLNACMECGACANNCPVDAISVNP--GVGCVEAYINRWLG
WP_045701157.1     1 --MQDFRYLDDTAILRLTAEKCVGCGNCALVCPHRIFRVRE---QKAEILDPNGCMECGACAINCPVGAIYVNPDDGCGCAAYIINSWIA
WP_136816054.1     1 --MKGFRYLDNTAILRLSEEKCIGCGSCELVCPHRIFRVRH---GKASITDHNRCMECGACAKNCPADAIYVNPDDGCGCAAYIINSWLA
WP_092224830.1     1 --MKTFRYLDNTAILHLTQEQCIGCGNCEIVCPHRIFKVVE---KKARITDKNSCMECGACARNCPVGAIFVNPDDGCGCAAYIINSWWN
WP_136873888.1     1 --MRTFRYLDNVAILTLAREKCIGCGNCEMVCPHRVFKVTG---GKADIKDPDGCMECGACARNCPVDAIYVNPDDGCGCAIYILNSWLG
WP_183351577.1     1 --MKDFRYIDGAAILRLDREACIGCGNCVTVCPHRIFRLED---NKAAVIDPDGCMECGACATNCPVAAIYVNPDDGCGCAALIINRWLS
WP_073612885.1     1 --MKDFRYLEETIQLRLDENLCTGCGNCVTVCPHRVFTMQD---HKAKIDDRGGCMECGACSLNCPVAAIFVNPDEGCGCATLLIHSWLQ
WP_136795961.1     1 --MKEFRYLENVAELTLDVDTCIGCGMCVTTCPHRIFTLKG---KKAEIVDFGACMECGACALNCPVNAIHVNPDDKCGCATLIIKSWIA
WP_136806099.1     1 --MKKLRYLEKLPILSLDTERCIGCGMCETTCPHRIFSIEN---GKAKLIDKGACMECGACATNCPVNAITVNEEEGCGCATLIISSWIS
WP_224981223.1     1 --MKGFRYLEGVATLELDGALCIGCGRCVEVCPHQVFCMEG---KRAVIVDRDACMECGACALNCPAAALKVDA--GVGCASGMIHEWLR
WP_216500699.1     1 --MKGFRYLQGVATLELSTPLCIGCGRCVEVCPHQVFRMEG---KRAVMVDRDACMECGACALNCPAAALKVDA--GVGCASGMIHEWLR
WP_216799213.1     1 --MKGFRYLEGVATLELAAPLCIGCGRCVEVCPHQVFRIEG---EKALVVDRDACMECGACALNCPAAALKVDA--GVGCASGMIHEWLR
WP_216511558.1     1 --MKGFRYLAGVATLELESSLCVGCGRCVEVCPHQVFRMAQ---KRAVVADRDACMECGACALNCPAAALKVDA--GVGCASGLIHEWLR
WP_015719368.1     1 --MKGFRYLEGVATLELERSLCVGCGRCTEVCPHQVFRIEQ---KKAVLADRDACMECGACALNCPAAALKVDA--GVGCASGMINEWLR
WP_185242963.1     1 --MKGFRYLEGVVTLELNRELCVGCGRCIEVCPHQVFRLEE---KRALLADRDACMECGACALNCPAAALKVDA--GVGCASGLINEWLR
WP_129127896.1     1 --MKGFRYLASVVTLELNAALCVGCGRCVEVCPHQVFRIEE---KKAALADRDACMECGACALNCPTSAIKVDA--GVGCASGLINEWLR
WP_151128529.1     1 --MKGFRYLKDVVTLELDQAACIGCGRCLEVCPHQVFSLAE---KKASMADRDACMECGACALNCPVKAITVDA--GVGCASGMINEWLQ
WP_223923564.1     1 --MEGFRYLRDVVTLELDRAACIGCGRCLEVCPHQVFSLAE---KKAALRDRDACMECGACALNCPVKAITVDA--GVGCASGMINEWFQ
WP_214300419.1     1 --MKGFRYLKDVATLNLDQPTCIGCGRCLEVCPHQVFSLVG---KQASIVDRDACMECGACALNCPVKAITVDC--GVGCASGMINEWLQ
WP_214169658.1     1 --MKGFRYLSSVVTLELDQAACIGCGRCQEVCPHQVFAVTA---KKAVMQDRDACMECGACALNCPVKAIKVDA--GVGCASGMIHEWLR
WP_088535263.1     1 --MKGYRYLKDATTLELREPLCIGCGRCIEVCPHQVFRLSE---KKAAVADRDACMECGACARNCPVNAITVDA--GVGCASGLINEWLQ
WP_173200239.1     1 --MQGFRYLENVVTLDFDPSLCIGCGRCLQVCPHQVFSPDG---KKVTLSDRDACMECGACAFNCPVKAISVDS--GVGCASGMINEWLR
WP_155876367.1     1 --MKGFRYLKDVVTLDLDRSACIGCGRCGEVCPHHVFRIAD---RKASIVDRDACMECGACASNCPVVAISVEA--GVGCAIGLFNEWLR
WP_199386911.1     1 --MKGFRYLEDVTTLQFDPARCTGCGRCVEVCPHQVFAVQG---RKAELMDRDACMECGACAKNCAARAITVDA--GVGCASGLINEWLQ
WP_145023550.1     1 --MKGYRYLSGVTTLALRQEVCVGCGRCLEVCPHRVFALDG---TEARIIDRDACMECGACARNCPAAAITVDS--GVGCASGLINEWLQ
WP_108102290.1     1 --MKGFRYLENVATLRFASDRCTGCGRCVEVCPHQVFEPAG---EIVAVVDRDACMECGACRMNCPHDAIQVDA--GVGCASGLLHEWLR
WP_041972229.1     1 --MKGFRYLNNVATLELDEAACIGCGRCLEVCPHQVFSVEN---RKAALIDKDACMECGACARNCPVKAIRVEA--GVGCASGIINEWLR
WP_039740952.1     1 --MRGFSHLKNVATLELDRAACIGCGRCPEVCPHGVFWLDG---GKAALADRDLCMECGACARNCPVAAIRVDA--GVGCASGMITEWLR
WP_011937421.1     1 --MKDFRYLRNVATLELRESACIGCGRCVEVCPHKVFSLAE---KRSRIVDFDGCMECGACAKNCPTAAIKVDA--GVGCASGLITEWLR
WP_054698307.1     1 --MKGFRYLQNATTLELNESACIGCGRCLEVCPHQVFTLAD---KRARLADRDACMECGACQRNCPTAAIRVDA--GVGCASGIINEWLK
WP_183356731.1     1 --MKGFRYLSNVATLELDQKRCVGCGRCLEVCPHQVFALEA---KLARIVDRDACMECGACAGNCPVRAITVDS--GVGCATGLINEWLR
WP_183360605.1     1 --MKGFRYLSNVATLELDQGRCIGCGRCLEVCPHQVFVLEA---RRACIGDRDACMECGACAGNCPVDAISVDS--GVGCATGIINEWLR
WP_250677262.1     1 --MKGFAYLKNVATLELNREKCVGCGRCLEVCPHQVFSLAA---RTAAIEKKDACMECGACAKNCPVGAIAVDA--GVGCASGMINEWLR
WP_243372457.1     1 --MKNFAYLRKVTTLKLERELCSGCGRCVEVCPHQVFSLTD---GISAVANLDACMECGACARNCPAGATEVDA--GVGCASGLIHEWLR
WP_223911758.1     1 --MKGFSYLKNVATLELDKDTCIGCGMCTVVCPHQVFALEG---EKAVIRAFDACMECGACALNCPVKAIRVDA--GVGCAAGMINEWLD
WP_214187022.1     1 --MKGFSYLTNVATLELNREACIGCGMCVAVCPHGVFALQG---TKAEIRDFDACMECGACAVNCSVGAIEVDS--GVGCASGMINEWLA
WP_207162600.1     1 --MKGFAYLPNVATLELSREICIGCGMCVTVCPHEVFVLQG---AKAEIINFDACIECGACSVNCPVEAIRVDS--GVGCASGLINEWIS
WP_039645259.1     1 --MIGFQYLANVATLKLAPERCIGCGMCVEVCPHQVFRLAG---KKAEIGTRDACMECGACAVNCPAAAIHVDA--GVGCASGMINEWLA
WP_119332422.1     1 --MIGFQYLANVATLEFIGEKCVGCGMCVEVCPHQVFRFEG---SKAKVVARDDCMECGACAVNCPASAIRVDA--GVGCASGMINEWLA
WP_029914610.1     1 --MDNFRYIKNVATLSLDAEHCIGCGRCVEVCPHAVLVLES---GRARIVDLDACMECGACARNCPVAVLSVDS--GVGCASGLLSEWLE
WP_100393475.1     1 --MKAFRYIDGVATLKLNRELCIGCGRCVVVCPQRILAMGE---RCVEVVDGDACMECGACAFNCSAGAITVNP--GVGCTSYIIAVWLS
WP_073378894.1     1 --MKDYRYIPDTATLRLDTETCIGCGLCQTVCPHRIFVVAD---KKARIEDRDACMECGACARNCPVEAITVTP--GVGCAVEIIARWVN
WP_015723166.1     1 --MHGYRYIDDTATLEVDFDRCIGCGNCTVVCPHRIFALEG---EKLKVGDRDLCMECGACARNCPVQALTVTP--GVGCAVAILAAWIN
WP_028584583.1     1 --MQAFRYIDGTVTLRLDRQDCIGCGRCSLVCPHRILRLEG---NRAVIVDRDRCMECGACQTNCPTRAITVSP--GVGCAAHIISQWLN
WP_205223032.1     1 --MDTFRYIEGAVTLQLDQEACIGCGSCVVVCPHRILYLEQ---RKAYITNVDLCMECGACQANCPTEAITVDP--GVGCAAYIIAKWVN
WP_205228011.1     1 --MRGYRYLQEVSTLSYTQERCIGCGLCVQVCPHQILTLSG---KQVHIRDIDLCMECGACAHNCPTEALSVRP--GVGCAVAILNGWKN
WP_028586334.1     1 --MKEFRYLPGVVSLELDAAACVGCGMCVTVCPHRVFALEG---GRAVLRDRDACIECGACARNCAPGAVRVSP--GTGCATLIIQSWLR
WP_167940925.1     1 --MEGFRYIPGVVSLEFDEKACVGCGLCIQVCPHRVFELVA---GRARMRDRDACIECGACALNCPPEAIRVTP--GVGCAAHIIKSWLA
WP_078683520.1     1 --MEHYRYIPGVVSLNFDEEACTGCGLCVDVCPHRVFEMIN---GKARLRDRDGCVECGACAKNCEGRAIRVDP--GVGCAVCVIKSWLW
WP_029894243.1     1 --MSELRYLENVVSLDLDREACIGCGMCLTVCPRGVFRLKD---GKAEYADRDACIECGGCAQNCPVEAIAVKP--GAGCATLILNGWLG
WP_041228877.1     1 ----MLRYLENVVTLRLNTEACNGCGMCAIVCSHGVFGIED---RKAVIIDQDACIECGACAINCQVNAIRVQT--GAGCATGVLLGALG
WP_008871052.1     1 --MQDFRYLEDVLTLEYDPGLCMGCGSCVQVCPHRVFDVGEQ---KAVLLDPGGCMECGACALNCPAGAIKVRP--GVGCAQAILNSWLA
WP_020887724.1     1 --MRDFRHLENIVTLDYDREKCIGCGACPTVCPHEVFALRGD---KAELVDHGGCMECGACALNCPSGAITVRP--GVGCAQAILYGWLS
WP_020880624.1     1 --MREFRHFENVTTLALDRAACVGCGACVAVCPHAVLALDETG--KARLADPGGCMECGACATNCAAAAITVRP--GVGCAQAILNGWLS
WP_028575525.1     1 --MKTFRHLPDVATLCFDKNLCIGCGNCLDVCPHQVFALQDG---KALILDRDACMECGACMTNCPVEAVYVRP--GVGCAQAILYGWLV
WP_168888863.1     1 --MKDYRYLPDVATLTLNHDACVGCGLCVTVCPHRALIMKE---GKAHIADLNGCIECGACMRNCPTEAVNVNP--GTGCATLIMDNWLY
WP_053551250.1     1 -----MRYLEEVSTLKLDRQVCIGCGLCAMVCPHGVFSVEE---KKARILDLDRCMECGACARNCPVAALAVKA--GVGCASAILYGWLT
```

```
WP_020887724.1    1 --MRDFRHLENIVTLDYDREKCIGCGACPTVCPHEVFALRGD---KAELVDHGGCMECGACALNCPSGAITVRP--GVGCAQAILYGWLS
WP_020880624.1    1 --MREFRHFENVTTLALDRAACVGCGACVAVCPHAVLALDETG--KARLADPGGCMECGACATNCAAAAITVRP--GVGCAQAILNGWLS
WP_028575525.1    1 --MKTFRHLPDVATLCFDKNLCIGCGNCLDVCPHQVFALQDG---KALILDRDACMECGACMTNCPVEAVYVRP--GVGCAQAILYGWLV
WP_168888863.1    1 --MKDYRYLPDVATLTLNHDACVGCGLCVTVCPHRALIMKE---GKAHIADLNGCIECGACMRNCPTEAVNVNP--GTGCATLIMDNWLY
WP_053551250.1    1 -----MRYLEEVSTLKLDRQVCIGCGLCAMVCPHGVFSVEE---KKARILDLDRCMECGACARNCPVAALAVKA--GVGCASAIIYGWLT
WP_066728217.1    1 -MAEKLVYLEGVSTLKLDRERCIGCGLCAVVCPHAVFLVEA---GKAEITALDRCMECGACARNCPVTALQVKA--GVGCASAIIHSWLT
WP_221249322.1    1 --MKELRYLQGVSTLTLDVPTCIGCGMCSLVCPHAVFLVEQ---KKALITDRDRCMECGACARNCPVSAISVKA--GVGCASAIIKGWLT
WP_200889300.1    1 --MKELHYLENVATLHLDVETCIGCGLCASVCPHGVFEIIA---QKAQILERNRCMECGACARNCPVNALAVKA--GVGCASAIIHSWIT
WP_163299160.1    1 ---MEMRYLEGVATLALDEAACIGCGRCTEVCPHAVFEVAH---RKARIRDRDRCMECGACARNCPAGALRVQA--GVGCAAAIVMGWLT
WP_025322949.1    1 --MEGFRYLDGVATLELHADRCVGCGLCETVCPHGVLAVSD---GKARITDRDACMECGACARNCPVEALSVSA--GVGCAGAIIKSWLT
WP_027367780.1    1 --MQELRYLSGVASLELNRDKCAGCGMCTVVCPHRVFEFRE---GKAEAVDRDACMECGACALNCPTGAIRVSP--GVGCAQAIIKGWLT
WP_173083722.1    1 -MN-ALRYIEGVATLRLDESLCTGCGTCREVCPHAVFGHAP---GKARLADPGACMECGACALNCPEGALTVSQ--GVGCARAIIHGWIT
WP_243359980.1    1 -MA-GLKYIEGVATLRLDEEACTGCGMCREVCPHGVFGTEP---GKARIENLDACMECGACALNCREGALSVSQ--GVGCARAIIHGWIT
WP_027189825.1    1 -MK-EMRYIDGVATLELDRDACTGCGTCREVCPHGVFGHGT---GIAAIEDLDACMECGACALNCREGAITVSQ--GVGCARAIIHGWLT
WP_243312050.1    1 -MSGCCRYIPGVVTLRMDPHRCGGCGMCAQVCPHGVFVVEG---GTARIADRDACMECGACALNCSSGAIEVEQ--GVGCARAIIHGWIT
WP_085053428.1    1 -----MIYLAGVSTLQFDAERCTGCGMCAEVCPHGVFIIAS---AKASITNKDKCMECGACENNCAFDAIRVSS--GVGCAAAIINGMIT
WP_203472301.1    1 -----MKYLTNVASLQLSPEKCTGCGRCIDVCPHGVFVINN---KKVSIADKDICMECGACALNCEFGAISVNS--GVGCAAAIINSMIT
WP_103116131.1    1 --MREHAYLNDTVSLELAEEKCTGCGLCTLVCPHAVLRVED---GKARIDARERCMECGACAKNCAFDALKVEA--GVGCASAIIYGWLT
WP_085011807.1    1 --MREYAYLNEYVSLQLDSGKCTGCGLCQQVCPHAVLRVED---GKARIDARERCMECGACAKNCAFDALKVEA--GVGCASAIIYGWLT
WP_020676021.1    1 --MAHQKYLKNVVTLKLEQEKCVGCGLCAIVCPRAVFKIED---DLAWITALDNCMECGACVGNCPVGAITVHA--GVGCASAIIHSWLT
WP_035294481.1    1 -----MQYLQNVVSLKLNKEECIGCKMCINVCPHEVFTIED---KKAVIMRKDSCMECGACAQNCPSGAIEVKK--AVG----------
WP_250697392.1    1 -----MEYLKNVASLKLNEEKCTGCKMCINVCPHEVFVMED---KKAVIIRKDSCMECGACAQNCPSGAIEVKK--AVGUVLAILKGILA
WP_250697396.1    1 ---MAIAYIKNVAQIKLDREKCIGCRMCLNVCPHTVFEMQE---GKAKFKNKDACIECGACDRNCPVEAIEVKS--GVGUAYAIIKGWLT
WP_250697399.1    1 ---MKMNYLKNVAAIRLDQDKCIGCGICLEVCPHGVFEMLQ---GKANFHDKDLCIECGACDRNCPVKAIEVKS--GVGUAYAILKGWLT
WP_132848058.1    1 ---MEHKYLKNVVTLSLDVNQCIGCNMCINVCPHEVFTTSD---KKVQIHNKDKCMECGACAKNCPVAAIDVKS--GVGCAYAIIVGKIT
WP_250697397.1    1 ---MRHKYLKNVATIELDIEKCIGCKMCINVCPHQVLTIKD---KKAFIQNKDYCIECGACDKNCPVDAIMVKS--GVGUAYAILKGKLT
WP_250697398.1    1 ---MKMKYLKNVASIVFYQEKCIACGMCIKVCPHEVFIFEN---DKIKLDKRDLCMECGACSKNCPVNAIEVKQ--GVGUAYAILKGMLT
WP_068703362.1    1 --MKTLQYLKNVVTLKLKPDLCTGCGMCTVVCPHAVFEMAN---GKARIVDIDDCMECGACANNCRFGAISVKA--GVGCAAGILNGILR
WP_227016804.1    1 -----MRYLKNVVTLKLESERCIGCGMCTDVCPHGIFSIEK---GKAKVNNRDYCMECGACEKNCPAAAITVKT--GVGCAAAVINGLLR
WP_036935625.1    1 ---MKNRYLKNVSTLKLKTDKCTGCGRCLEVCPHEVFELEN---GKSHIIDRDACMECGACAKNCPFNAIDVKP--GVGCAYAIIMGWLT
WP_045660524.1    1 ---MNHKYLKNVTTLILKADKCTGCGRCIQVCPHKVFDLKN---GKSEIIDKDGCIECGACAKNCLFNAIEVKP--GVGCAYAIIMGWLR
WP_073538590.1    1 ---MKNKYLKNVCTLKVDSDKCVGCGRCIEVCPHKVFNLNK---SKIELVNKDACMECGACAKNCAFNAIEVKS--GVGCAYAVIMGWLL
WP_106064375.1    1 ---MKNKYLKNVCTLKLNSDKCVGCGRCIEVCPHKVFDLNQ---GKVELINKDGCMECGACAKNCAFNAIEVNS--GVGCAYAVIMGWLL
WP_131929766.1    1 ---MKHCYLKNVSTLTLKSEKCTGCGRCIEVCPHQVLGLKC---KKAEILDIDLCMECGACARNCPFEAINVKS--GVGCASAVITGWLT
WP_027629139.1    1 ---MRHRYLKNVATLKLDREKCKGCGKCIEVCPHKVFALRA---DKAEIMDRDGCMECGACAGNCPFGALEVKP--GVGCAAAIIMGWLT
WP_010243113.1    1 ---MKHRYLKNVVSLKLNTDKCKGCGRCAEVCPHRVFDLKN---SKAVIMDKDSCMECSACARNCPFGAIEVKT--GVGCAAAVITGWLT
WP_006718597.1    1 ---MRNTYLKNVSTLKIESDKCIGCGRCLEVCPHHVFILSE---RKSVIIDKDHCMECGACVRNCPVNALEVKP--GVGCASAIIKGWLT
WP_135380082.1    1 ---MRHQYLKNVTTLKLKSDRCKGCGKCLEVCPHKVFSLNN---GKSDIVDIDRCMECGACVKNCPFNALEVKP--GVGCAFAIIMGWLT
WP_068964682.1    1 ---MTNRYLKNVATLKLKSDQCTGCGRCLEVCPHNVFLIED---RKSVIVDKDRCMECGACVKNCPFNALEVKP--GVGCAFAIIMGWLT
WP_073031434.1    1 ---MKNRYLKNVATLKLNSDQCKGCGRCLEVCPHNVFVIED---RKAVILDKDRCMECGACVKNCPFNAIEVKP--GVGCAYAIIMGWLT
WP_007786009.1    1 ---MKNRYLKNVATLKINSDLCTGCGRCLEVCPHNVFLLKD---KKSVIVDKDQCMECGACVKNCPFDALEVKP--GVGCAFAVIMGWLT
WP_034140029.1    1 ---MRHQYLKNVSTLKMKPDQCTGCGRCLEVCPHNVFLIKD---KKSVIVDKDRCMECGACVKNCPSNAIEVKP--GVGCAYAIIMGWLT
WP_135551677.1    1 ---MKNLYLKNVATLKLKSELCTGCGKCLEVCPHNVFLIKD---RKSVIIDKDRCMECGACAKNCSFKAIEVKQ--GVGCAFAIIMGWLT
WP_106801060.1    1 ---MRNQYLKNVATIKLKSDQCTGCGRCIEVCPHNVFQMED---KRSVIVGKDRCMECGACVKNCPFNAIEVKQ--GVGCAYAIIMGWLT
WP_034614517.1    1 ---MRNRYFKNVSTLNLKSEQCTGCGRCIEVCPHNVFLIEN---RKTVIFDKDGCMECGACAKNCPFNALEVKP--GVGCAAGIIKGWLT
WP_014185008.1    1 ---MKHKYLKNVSTLKMESDKCTGCGKCLEVCPHHVFVMRE---GKSIIADKDSCIECGACVKNCPFKALEVNP--GVGCAAAIIKGWLT
WP_088227281.1    1 ---MKHSYLKNVSTLRMDSGKCTGCGMCLEVCPHHVFVMSK---GKSIIVDKDRCIECGACVKNCPFKALEVKP--GVGCAAAIIKGWLT
WP_014826631.1    1 ---MKNRYLKNVSTLKMEPEKCTGCGKCLEVCPHNVFVMTN---GKSVITEKDNCIECGACVKNCPFNALEVKP--GVGCASAIIKGWLT
WP_045573189.1    1 ---MKNRYLKNVATLKMKSDKCTGCGKCIEVCPHNVLSISN---GRAIIVAIDRCIECGACVKNCPFNALEVKP--GVGCASAVLKGWLT
WP_015261307.1    1 ---MKSQYLKNVATLKLEQSRCTGCGKCFEVCPHQVFAIMN---SKATIVEKDRCIECGACVKNCPFHALEVKP--GVGCASAIIKGWLT
WP_014794562.1    1 ---MKNQYLRNVSTLKLEQSLCTGCGKCIEVCSHQVFAIVN---SKCSVVNKDGCIECGACVKNCPFLALEVKP--GVGCASAIIKGWLT
WP_073278284.1    1 ---MKLKYLQNVATLKLNREKCVGCGMCLNVCPHGVFELTK---GKAQVVNLDNCMECGACLKNCAFSAITVSP--GVGCASAIIKGFLT
WP_073591208.1    1 ---MKHKYLKNVATLQLNKEKCVGCGRCKDVCPHGVFVLDG---GKTQIVDLDSCMECGACLKNCAFSAITVSP--GVGCASAIIKGLLT
WP_034838824.1    1 ---MKYQYLKNVATLKLDSEKCVGCGMCVNVCPHNVFELFA---GKAKIIDLDSCMECGACAKNCAFSAITVTP--GVGCAAAIIKKFLT
WP_186428912.1    1 ---MKNKYLRNVATLKLYEEKCTGCGMCLNVCPHDVFEIRE---RKSHIIDKDSCMECGACAKNCGFSAIEVDS--GVGCAYAIIVGKLT
WP_069194727.1    1 ---MKHRYLKNVATLRLSAEKCIGCGRCTEVCPHGVFSVNE---NKAKIKDKDRCMECGACSKNCPTNAIAVDS--GVGCAAAVIIGWLK
WP_038290751.1    1 ---MKHRYLKNVATLRLSAEKCIGCGRCTEVCPHGVFSVNE---KKAKIEDKDFCMECGACAKNCPSNAITVDA--GVGCATAVIMGWLT
WP_204614197.1    1 ---MKHRYLKNVATLRLSAEKCIGCGRCAEVCPHRVFSVNE---KKAQIENKDHCMECGACAKNCPANAITVDA--GVGCAFAVIMGWLT
WP_069194638.1    1 ---MKHRYLKNVATLGLSAEKCIGCGRCTEVCPHGVFSMNE---NKAQVEDKDSCMECGACAKNCPATAITVDA--GVGCASAVIMGWLT
WP_013625503.1    1 ---MKHRYLKNVATLNLLVEKCVGCERCTEVCPHGVFGIAD---KRAKLKDRDSCMECGACAMNCPVNAISVEA--SVGCAAAIIRGWFT
WP_080064583.1    1 ---MRYKYLKNVTTLCLSAEKCICCGRCTEVCPHGVFGIHD---KKARIKDKDWCMGCGACAINCPVKAIDVDV--GVGCAAAVIIGWLT
WP_207857747.1    1 --MIRHTYLKNVVTLTLNANKCIGCGMCLNVCPHKVFGLNS---GKAWISDRDRCMECGACVRNCPVEALAVKA--GVGCAAAIIQGKLK
WP_045505539.1    1 --MKQMSYLQNGQTLRFNAERCVGCGRCVEVCPHQVFALEQ---RKASIMHREWCMECGACQKNCPVNAIAVQA--GVGCAAAIIQGYLK
WP_132779871.1    1 ---MEFRYILAGQSLTLDTEKCIGCLMCCEVCPHNVFSFAE---GKARIADRGRCMECGACARNCPVSALTVTS--GVGCTAAVIGGLLK
WP_250697394.1    1 ---MGLTYLKNVTTLKLDTDKCSGCRLCTEVCPHQVFAMEN---KKSRIANKEACMECGACARNCPTEAITVHS--GVGUAYAIILGKLT
WP_250697395.1    1 ---MSREYIKNVVTLNLDRDKCIGCKNCETVCPYRIFKIEE---RKAKIINKDLCIECGACMNNCPSEAITVRK--GVGUVLAIVKGVLT
WP_084055567.1    1 -MRGNLVYLKDVVSLQYEASRCVGCGMCSLVCPHGVFVQEN---GKARITDRDACMECGACARNCPTEAIRVQT--GVGCAQAVINSVLG
WP_073040146.1    1 -MGPNLVYLKDVVTLRYEASRCVGCGMCSIVCPHGVFVQEN---GKARITERDACMECGACARNCPAEAIWVQT--GVGCAQAVINSALG
WP_123289891.1    1 -MG-RLVYLKDVVTLQLDAAKCIGCGECTLVCPHGVFVKEN---GVARIAFRDACMECGACARNCPADAIAVKT--GVGCAQAVLNTMIG
WP_041585151.1    1 -ME-TLMYLDNVVTLRLDEEKCIGCGMCLEVCPRTVLSLEK---GRARISNRDACIECGACSRNCPVDAFAVGT--GPGCATAVINSMLG
WP_037464304.1    1 -MG-NLIYLKNVATLNMDEKKCTGCGMCMEVCPHHVFKMNG---KSALIIDRDACMECGACSQNCPAGAISVQS--GVGCAAAVINSLLG
WP_039659148.1    1 -MK-NPVYLKDVVTLQLDENKCTGCGMCLDVCPHEVFRMNS---KYVTIQNRDACMECGACSLNCPANAISVQS--GVGCAAAVINSMLG
WP_228853893.1    1 -MT-QFCYLEDVVTLACNIDKCIGCGMCTLVCPHGVFTIKG---KKAVISARNYCMECGACMRNCPVSAISVEA--GEGCVRGVINELLG
WP_049676292.1    1 --MRPL-YLHGVSTLVLDPEKCLGCGSCLPVCPHQILYLAN---GRIACSERDACMECGACAMNCPVGAVAVEA--GVGCAAAVLASAFR
WP_211213743.1    1 MPMRKLVYLHDVVTLNLDETRCSGCGMCLQVCPHGVLAKHN---GSVEIVNRDACMECGACAKNCPTAALSVKT--GVGCATAVINYALG
WP_006425356.1    1 --MEKFTYLRDVVTLALDQEKCVGCGMCLMVCPQEVLSMNN---GSAGILNRDRCMECGACARNCPTEAITVKV--GVGCAAALINAALG
WP_181550703.1    1 MGTTSFTYLKNVTTLELNPELCTGCGMCTMVCPREVMALAD---GKAEIVSRDDCMECGACAQNCPTAAITVDA--GVGCAAAVINSMLG
WP_106818687.1    1 -MS-KLVYLKDVVTLQLNSESCSGCAMCATVCPRGVFEISH---RKAKVADRDACIECGACAQNCPEGAVSVRV--GVGCASAVINSMLG
WP_051408865.1    1 -MQ-SMVYLKDVVTLAHKPEMCNGCGICLTVCPRRVLRRSN---GKVEVAQRDACIECGACQRNCPQGAVTVRT--GVGCASALINQMLG
WP_069894202.1    1 -----MTYLKNITTISVNAEDCTGCGRCQEVCPRGVLVREG---KKVAVADRDACIECGACMRNCPYAAVTVEA--GVGCAAAFINGALS
WP_044350425.1    1 -MSRSPLYIKNVSTLLLFSDLCIGCGTCLSVCPHQVLKLVK---GKVALKCLDACMECGACALNCPAKAITVNS--GVGCASSIILTALG
WP_116482024.1    1 -MFNS----YASNTLKYDPELCINCGMCSTVCPHRVFSPDE---KIAQLTNPKACMECGACQLNCPTKAIKVDS--GVGCASAMMLAAIK
WP_012617702.1    1 MIFDS----YSENTLAYDSEQCVNCGACSTVCPHRVFTPGK---KAAILTTPAACMECGACQVNCPTGAITVES--GVGCASALIRVALT
WP_019177912.1    1 MVFVEDIVQNSLNTLRYLPGRCVNCGRCSEVCPHGVFEAGE---KRARLAAPERCMECGGCQMNCPAGAIGVES--GVGCAAAMIKAALT
WP_230740017.1    1 MESLS-----IINTLHFDELLCINCGMCSIVCPHGVFSRGE---RKAVPVSPDSCMECGACQLNCPVDAIVVDS--GVGCASAMIKAALT
WP_012899469.1    1 -MFNS----YKVNTLKFDPEECTGCGMCTKVCPHAVFAYQD---KKVVLANKEACMECGACFLNCPTTAIEVES--GVGCAAAMIKAALT
WP_167816154.1    1 -MFDS----YTETSLSFDPALCINCRRCTQVCPHAVFTAGQ---TRVILDRPSACMECGACALNCPVQAITVQS--GVGCAWGMIGAALR
WP_015284971.1    1 -MFNS----YTETSLELHADRCINCKRCMQVCPHAVFTEGR---EHVILSRPAACMECGACAKNCPVQAIEVQS--GVGCAWAMIGAALR
WP_015052521.1    1 -MFDS----YLENTLLYYPQKCINCLRCTQVCPHGVFAEGK---EHAELMQPAACMECGACVRNCPVQAIEVQS--GVGCAWAMISAALR
WP_015324054.1    1 -MFNS----YIENTLQYYPEKCINCLRCTHVCPHEVFGEGK---GHVELMRPAACMECGACARNCPVQAIEVQS--GVGCAWAMIGAALR
WP_094228742.1    1 -MFDS----YRENTLQYYPDRCINCLRCTQVCPRGVFTEGK---EHVELTQPAACMECGACARNCPVLAIEVQS--GVGCAWAMIGATIR
WP_209616048.1    1 -MLNS----YIENTLQFYQEKCINCLICTQVCPHGVFTEGK---EHVELTKPSACMECGACAVNCPVEAIDVQS--GVGCAWAMIGAALK
WP_091933276.1    1 -MFNS----YNENTLQFHPEKCINCLICTQVCPHGVFTEGK---GHVELTSPEKCMECGACAGNCPVEAIEVQS--GVGCAWAMINAALK
WP_233083665.1    1 -MFNS----YLENTLQYYPEKCINCLMCIQVCPHGVFTGGK---EHVELKSPASCMECGACAGNCPVQAIEVES--GVGCAWAMISAALK
WP_023846499.1    1 -MFNS----YKENTLKYYPEKCINCLMCTNVCPHGVFTDGE---KQVMLTKPELCMECGACAGNCPVEAIEVQS--GVGCAWAMISAALK
WP_176965778.1    1 -MFDS----YNENTLQYYPEKCINCLMCTQVCPHGVFTGGG---EHVELKRPSSCMECGACAMNCPVQAIEVQS--GVGCAWAMIRAALR
WP_109967936.1    1 -MFNS----YTENTLNFHQDKCINCRRCTEVCPHGVFEEGM---KSVELKNPVRCMECGACALNCPTQAIEVQS--GVGCAWAMIGAAIR
WP_214046877.1    1 -MFNS----YQENTLHYHEEKCINCRRCTEVCPHGVFAEGA---DHVTLVYPTRCMECGACALNCPVQAIEVES--GVGCAWAMISAALR
WP_214418654.1    1 -MFNS----YIENTLQFHEEKCINCKRCTEVCPHGVFAEAG---GRVNLSEPNRCMECGACALNCPVQAIEVES--GVGCAWAMISAALR
WP_011447904.1    1 -MFHS----YLDNSLKFYKNRCINCKRCTEVCPHGVFSAGK---SHVNLVYQVRCMECGACALNCPVQAIEVES--GVGCAWAMISAALR
WP_209674662.1    1 -MFDS----YVETTLRYYPERCINCRRCTQVCPHGVFSEGP---ERAVLRHPPDCMECGACALNCPVQAIEVQS--GVGCAWAMISAALR
WP_221056473.1    1 -MFDS----YRENTLRYNPERCFGCLRCTEVCPHGVFAEGN---RRVEVVRPEECMECGACARNCPVQAIAVQS--GVGCAWAMIGAALR
WP_004039396.1    1 -MFYS----YGENTLRYHPERCFNCRRCIEVCPHGVFAAGD---RRVACVSPERCMECGACARNCPVQAIEVES--GVGCAWAMIGAALR
WP_011991428.1    1 -MPDS----YAENTLRFYAEKCINCRRCTEVCPHGVFAEGE---KTAVLGNPRACMECGACAKNCPVQAISVQS--GVGCASAMIRAALK

WP_250697400.1   84 GKENASCGGDAG---------CC-------------------- 97
[illegible]
```

```
WP_221056473.1    1 -MFDS----YRENTLRYNPERCFGCLRCTEVCPHGVFAEGN---RRVEVVRPEECMECGACARNCPVQATAVQS--GVGCAWAMIGAALR
WP_004039396.1    1 -MFYS----YGENTLRYHPERCFNCRRCIEVCPHGVFAAGD---RRVACVSPERCMECGACARNCPVQAIEVES--GVGCAWAMIGAALR
WP_011991428.1    1 -MPDS----YAENTLRFYAEKCINCRRCTEVCPHGVFAEGE---KTAVLGNPRACMECGACAKNCPVQAISVQS--GVGCASAMIRAALK

WP_250697400.1   84 GKENASCGGDAG---------CC-------------------- 97
WP_250697390.1   84 GKENASCG--DG---------CC-------------------- 95
WP_250697391.1   84 GKEKASCG-SGG---------CC-------------------- 96
WP_250697393.1   84 GKENASCG-IGG---------CC-------------------- 96
WP_045674113.1   84 GKEKATCG--PG---------CC-------------------- 95
WP_092212500.1   84 GKDNASCGGG-----------CC-------------------- 95
WP_236890014.1   84 GKENASCGGGEG---------CC-------------------- 97
WP_155317485.1   84 GKENASCGGEGG---------CC-------------------- 97
WP_213183181.1   85 GKENASCGGPDG---------CC-------------------- 98
WP_144305632.1   84 GSKAAQAK-N-----------CC-------------------- 94
WP_144301671.1   84 GTRYADAK-FS----------CC-------------------- 95
WP_163302676.1   84 G----KAK-AS----------CC-------------------- 91
WP_176630762.1   84 G----QAR-AS----------CC-------------------- 91
WP_027186353.1   84 -KDNSKG---TA---------CC-------------------- 93
WP_037564333.1   84 GKDKAACG-SAE---------CC-------------------- 96
WP_126379058.1   84 WTKNAGG--------------CC-------------------- 92
WP_022661037.1   86 -------RRAKG---------CC-------------------- 92
WP_158950321.1   84 ---RLTGRKVTS---------CC-------------------- 94
WP_066802544.1   84 ---KMTGKQVDT---------GCC------------------- 95
WP_229594619.1   84 ---QLTGKEMDS---------GCC------------------- 95
WP_071544227.1   84 ---KLTGRKVDS---------ACC------------------- 95
WP_014321698.1   84 ---RLTGRKIDA---------ACC------------------- 95
WP_128329673.1   84 ---KLTEKKVST---------GCC------------------- 95
WP_013515565.1   84 ---RLTGNKVKT---------TCC------------------- 95
WP_020875938.1   84 ---RIKG-SAGT---------RCC------------------- 94
WP_097012721.1   84 ---QLTGKEIKN---------TCC------------------- 95
WP_045222548.1   84 KS-------TTP---------QCC------------------- 91
WP_031386617.1   84 KS-------SAP---------QCC------------------- 91
WP_028570972.1   84 KS-------SAP---------QCC------------------- 91
WP_205240453.1   84 GK-------RGT---------SCC------------------- 91
WP_012805587.1   84 GK-------RGT---------SCC------------------- 91
WP_092118529.1   84 RK-------RSK---------GC-------------------- 90
WP_054649311.1   84 SRNEAI---------------NCC------------------- 92
WP_018125557.1   84 IRS-SG---------------GCC------------------- 91
WP_205232270.1   84 GKNLP--SG------------SCC------------------- 93
WP_174408776.1   84 EKGLPVRMG--N---------GCC------------------- 96
WP_174404739.1   84 EKGIPVRLGGGT---------GCC------------------- 98
WP_230996100.1   84 G-KKFGGTGS------------CCG------------------ 95
WP_175641722.1   84 DK-RITKSSGGG---------CCC------------------- 97
WP_163335649.1   84 ---RLTGRKGGG---------SCC------------------- 95
WP_084813749.1   84 ---RLFKDRGRS---------SCC------------------- 95
WP_045701157.1   86 ---RMK----TG---------CGC------------------- 93
WP_136816054.1   86 ---GIRGKPPSG---------CGC------------------- 97
WP_092224830.1   86 ---RLRGKSNSP---------CGC------------------- 97
WP_136873888.1   86 ---KITGKRSSD---------CGC------------------- 97
WP_183351577.1   86 ---RITGKTVSG---------CNC------------------- 97
WP_073612885.1   86 ---RITGRKSSS---------ACC------------------- 97
WP_136795961.1   86 ---KYTGKTLGG---------GCC------------------- 97
WP_136806099.1   86 ---KFTGKPSSG---------SCC------------------- 97
WP_224981223.1   84 E-KKLPGLNGGG----------CCG------------------ 97
WP_216500699.1   84 E-RKIPGFGGGG----------CCG------------------ 97
WP_216799213.1   84 E-KKIPGISSGG----------CCG------------------ 97
WP_216511558.1   84 E-RKIPGFNGGG---------GCCG------------------ 98
WP_015719368.1   84 E-KKFPGFGGAG---------GCG------------------- 97
WP_185242963.1   84 E-KRIPGFGGSG----------CCG------------------ 97
WP_129127896.1   84 E-KKLPGLKGSG----------CCD------------------ 97
WP_151128529.1   84 E-HKL-RAPSGG---------CCS------------------- 96
WP_223923564.1   84 E-RKL-RLPGGN---------CCS------------------- 96
WP_214300419.1   84 E-RKLGKLGGGN---------CCS------------------- 97
WP_214169658.1   84 E-HKLEKLAG-D---------CCS------------------- 96
WP_088535263.1   84 E-RKL-RIPGGG---------CCS------------------- 96
WP_173200239.1   84 E-RKLVKTGGGG---------C--------------------- 95
WP_155876367.1   84 E-RKLYKPKEC-------------------------------- 93
WP_199386911.1   84 E-RKLLKGIKTG---------GCCQ------------------ 98
WP_145023550.1   84 E-RKLRRVGSG-----------CCS------------------ 96
WP_108102290.1   84 E-RNLSSSNRS-----------CCS------------------ 96
WP_041972229.1   84 E-RNLRASGGE-----------CCS------------------ 96
WP_039740952.1   84 E-RNIRWSGAD-----------GC------------------- 95
WP_011937421.1   84 E-RNIRGVGGG-----------CCS------------------ 96
WP_054698307.1   84 D-HNIRLGGSG-----------CC------------------- 95
WP_183356731.1   84 E-KKIPGVGGGG-----------CG------------------ 96
WP_183360605.1   84 E-KKIPGLGGGS-----------CG------------------ 96
WP_250677262.1   84 E-HNIRGRSGE-----------C-------------------- 94
WP_243372457.1   84 E-RNIRGWGGE-----------CCG------------------ 96
WP_223911758.1   84 S-LNLRR-RGDG---------CC-------------------- 95
WP_214187022.1   84 S-IGARKTRGGG---------CC-------------------- 96
WP_207162600.1   84 S-ITGRKTGGGG---------CC-------------------- 96
WP_039645259.1   84 G-VVGNK--GGG---------CC-------------------- 94
WP_119332422.1   84 G-VVGSR--GGR---------CC-------------------- 94
WP_029914610.1   84 E-KFSLRIKGG-----------C-------------------- 94
WP_100393475.1   84 ---RFRRRKVTS---------ACC------------------- 95
WP_073378894.1   84 ---GITGRTVMK---------GCC------------------- 95
WP_015723166.1   84 ---RLLGRKLLS---------GCC------------------- 95
WP_028584583.1   84 ---TLLGRQVFK---------GCCE------------------ 96
WP_205223032.1   84 ---RLLGREIVK---------GCC------------------- 95
WP_205228011.1   84 ---RLLGTTKFG---------DCC------------------- 95
WP_028586334.1   84 GL-GLAGRSAGG---------CCGG------------------ 98
WP_167940925.1   84 TL-GLC-RSSSS---------CC-------------------- 95
WP_078683520.1   84 SM-GLR-VSRPS---------CR-------------------- 95
WP_029894243.1   84 -------RSTKD---------CC-------------------- 90
WP_008871052.1   84 GMPVLRRFFRRD---------SCCT------------------ 99
WP_020887724.1   84 RVPLLRRVVAPD---------ACCS------------------ 99
WP_020880624.1   85 RLPFLRRLSGAD---------SCCS------------------ 100
WP_028575525.1   84 KVPILRKVLSPD---------SCCP------------------ 99
WP_168888863.1   84 EM-SGG-RLRKK---------CC-------------------- 95
WP_053551250.1   81 GKEP--SCDCG--------GESCC------------------- 94
WP_066728217.1   85 GEEP--SCDCG--------GSSC-------------------- 97
WP_221249322.1   84 GSEP--NCDCG--------G----------------------- 93
WP_200889300.1   84 GEEP--SCDCGTG----QSGSGC-------------------- 100
WP_163299160.1   83 GRTP--ACGCGSDG--------CC------------------- 96
WP_025322949.1   84 GAPP--TCGCEG-------GGGC-------------------- 97
WP_027367780.1   84 GGPP--NCDCGSD---------CC------------------- 96
WP_173083722.1   84 GGPP--SCDCGG-----DGSSTCC------------------- 100
```

```
WP_200889300.1   84 GEEP--SCDCGTG----QSGSGC-------------------- 100
WP_163299160.1   83 GRTP--ACGCGSDG--------CC------------------- 96
WP_025322949.1   84 GAPP--TCGCEG-------GGGC-------------------- 97
WP_027367780.1   84 GGPP--NCDCGSD---------CC------------------- 96
WP_173083722.1   84 GGPP--SCDCGG-----DGSSTCC------------------- 100
WP_243359980.1   84 GGPP--SCDCGG-----DGGSTCC------------------- 100
WP_027189825.1   84 GGPP--SCDCGG-----DGGSTCC------------------- 100
WP_243312050.1   85 GGPP--SCDCGG-----DGGSTCC------------------- 101
WP_085053428.1   81 GGEP--SCGCS------SEGGTCC------------------- 96
WP_203472301.1   81 GGPP--SCDCGGSSS-QSASSNCC------------------- 101
WP_103116131.1   84 GKEA--SCDCG-------GG--C-------------------- 95
WP_085011807.1   84 GKEA--SCDCS-------GG--C-------------------- 95
WP_020676021.1   84 GDEP--SCDCG-------GGGNC-------------------- 97
WP_250697392.1   81 GCRK--------------------------------------- 84
WP_250697396.1   83 GSRP------------------CC------------------- 88
WP_250697399.1   83 GSRP------------------C-------------------- 87
WP_132848058.1   83 GREPDCGCSDEI---------DCC------------------- 97
WP_250697397.1   83 GSEP-----------------NCC------------------- 89
WP_250697398.1   83 GSEP------------------RC------------------- 88
WP_068703362.1   84 GTEPSCDCSTGSKA--------CC------------------- 99
WP_227016804.1   81 GTEPSCDCSENSNS--------CC------------------- 96
WP_036935625.1   83 GTEPSCDCSGGSSS-------GCC------------------- 99
WP_045660524.1   83 GTEPSCDCSGSSSSS------GCC------------------- 100
WP_073538590.1   83 GTEPTCDCSGDNSS-------GCC------------------- 99
WP_106064375.1   83 GTEPTCDCSGGSSG-------GCC------------------- 99
WP_131929766.1   83 GTEPSCDCSGGSSG-------GCC------------------- 99
WP_027629139.1   83 GSEPSCDCSSGDSSS------GCC------------------- 100
WP_010243113.1   83 GTEPSCGCADDNNSG------GCC------------------- 100
WP_006718597.1   83 GTEPSCDCSE---G-------GCC------------------- 96
WP_135380082.1   83 GTEPSCDCSGDNGG-------SCC------------------- 99
WP_068964682.1   83 GTEPSCDCSGGSGG-------GCC------------------- 99
WP_073031434.1   83 GTEPSCDCSGGSGD-------NCC------------------- 99
WP_007786009.1   83 GTEPSCDCSSGSGG-------SCC------------------- 99
WP_034140029.1   83 GTEPSCDCSPDSGD-------SCC------------------- 99
WP_135551677.1   83 GTEPNCDCSGSSGG-------SCC------------------- 99
WP_106801060.1   83 GKEPSCDCSDSSSG-------NCC------------------- 99
WP_034614517.1   83 GTEPSCGCSDGGGD-------SCC------------------- 99
WP_014185008.1   83 GTEPSCDCSD---G-------SCC------------------- 96
WP_088227281.1   83 GTEPSCDCSD---G-------SCC------------------- 96
WP_014826631.1   83 GTEPSCDCSD---G-------SCC------------------- 96
WP_045573189.1   83 GTEPSCDCSDNSGG-------GCC------------------- 99
WP_015261307.1   83 GTEPSCDCSPDS-G-------GCC------------------- 98
WP_014794562.1   83 GTEPSCDCSSDS-G-------SCC------------------- 98
WP_073278284.1   83 GSEPSCDCSGGE----------SC------------------- 96
WP_073591208.1   83 GSEPSCDCSGGE----------CC------------------- 96
WP_034838824.1   83 GGEPSCDCSNSD----------CC------------------- 96
WP_186428912.1   83 GTEPNCGCSGENQG--------CC------------------- 98
WP_069194727.1   83 GSEPSCDCSDSE---------GCC------------------- 97
WP_038290751.1   83 GKEPSCDCSSSE---------ECC------------------- 97
WP_204614197.1   83 GSEPSCGCSSGE---------ECC------------------- 97
WP_069194638.1   83 GSEPSCDCSSGE---------ECC------------------- 97
WP_013625503.1   83 GSEPSCDCSGSG---------DCC------------------- 97
WP_080064583.1   83 RSEPSCDFQASE---------NAAKK----------------- 99
WP_207857747.1   84 GTAPCCGGSEGNK-S------SCCG------------------ 101
WP_045505539.1   84 KTAPQCGCSTSS--S------GCCG------------------ 100
WP_132779871.1   83 GGEPNCDCDCGTAKS---KKVRCC------------------- 103
WP_250697394.1   83 GREPTCGSRDDG---------CGC------------------- 97
WP_250697395.1   83 GRGKKA------------------------------------- 88
WP_084055567.1   85 RTG-DSCCSLDSG---------CC------------------- 98
WP_073040146.1   85 RKG-DSCCSLDSG---------CC------------------- 98
WP_123289891.1   84 RSG-DGCCSLDSG---------CC------------------- 97
WP_041585151.1   84 RRG-ECCCSLDDGDEKGGGSSSCCC------------------ 107
WP_037464304.1   84 RNGGECCCGPQTDETGARKSGGCCS------------------ 108
WP_039659148.1   84 KDG-ECCCGPDTGGN-ASGNKSCC------------------- 105
WP_228853893.1   84 IKG--PCC----------------------------------- 89
WP_049676292.1   83 RGESGCC------------------------------------ 89
WP_211213743.1   86 RNSSSCCCVIEPDAENSNQAPTQSSGCC--------------- 113
WP_006425356.1   84 RDGSSCCCVIET--ENSGTFASGNPGCC--------------- 109
WP_181550703.1   86 RTGDSCCCVVEP---DGKSRPECGPGCC--------------- 110
WP_106818687.1   84 INNSGCCSMDQYENDKAAAKGTGC------------------- 107
WP_051408865.1   84 RKK-ACCVADDAPG-KTIASNTAR------------------- 105
WP_069894202.1   81 GGEPNCGCG---------SGGCCG------------------- 95
WP_044350425.1   85 LGSQKVCC----------------------------------- 92
WP_116482024.1   81 GQKEATCG--CDSP---------C------------------- 93
WP_012617702.1   82 GKGEETCD--CCSERELADERSYCEGGCSETGCR--------- 113
WP_019177912.1   86 G-KEATCGPDCCR------------------------------ 97
WP_230740017.1   81 GSKKVTCG--CDDDECEAEEAPCCCCEK--------------- 106
WP_012899469.1   81 G-GPETCDCGCDETD---KPPGCC------------------- 100
WP_167816154.1   81 GKDMDT-ESCCG------EGSSCCGSSAPKQKALICDCGKKER 116
WP_015284971.1   81 GKDMDT-ECSCGG-----EDGSCCGNSST-------------- 103
WP_015052521.1   81 GKGMDSGECGCGG-----EQMSCCGGGAKESFSNREEMSTPED 118
WP_015324054.1   81 GKDMDSSECCCGG-----EDGSCC------------------- 99
WP_094228742.1   81 GKDMDSADCCCGG-----DTGSCCGG----SVKNKRDQ----- 109
WP_209616048.1   81 GKDMDSDACCCG------EDGCC-------------------- 97
WP_091933276.1   81 GKDMDSDACCCG------DDGCC-------------------- 97
WP_233083665.1   81 GKDMDSDACCCGE-----DDGCCQ------------------- 99
WP_023846499.1   81 GKDMDSEACCCG------EDGCCQ------------------- 98
WP_176965778.1   81 GKDMDSADCCCN------GDSC--------------------- 96
WP_109967936.1   81 GKDMDSGGCGCGE-----E--SCCGE----------------- 99
WP_214046877.1   81 GKDIDSGECSCGG-----DSGSCCGVEQEQASCGCGK------ 112
WP_214418654.1   81 GKDMDSGECSCGG-----GAGSCCGGETEGSSCGCGEG----- 113
WP_011447904.1   81 GRDMDSGECSCGG-----DAGSCCGMK---------------- 102
WP_209674662.1   81 GKDMDSGECGCGG-----DDGACCGGGEEEEPSSCCEGN---- 114
WP_221056473.1   81 GKDMDSGTCGCGG-----DEGSCCGGDR--------------- 103
WP_004039396.1   81 GKDMDSGVCGCG-------DGSCCGGGQEEPSCCGER------ 110
WP_011991428.1   81 GKDMDGAECSCGGP----GASSCCGGEACEEGCRTDSQEKAV- 118
```
