## Supplemental Figure 5 for "Eight Unexpected Selenoprotein Families in ABC transport, in Organometallic Biochemistry in *Clostridium difficile* and other anaerobes, and in Methylmercury Biosynthesis"

ABC transporter substrate-binding protein SaoX

```
WP_248000900.1    1 ------MTIRKRLLCLMLAASMLMMGLSGCKDSSND----TNTVQIDYNAEEEVAKYKDYLGELSDADKDYVIELGYNNCDHMVASLIGQGSGIFDALDLN-VNVTKTG-
WP_248000901.1    1 ------MTMRKRLLCLMLAASMLIVGLTGCKDSSND----TNTVQVDYNAEEEVAKYKDYLGEIPEADKDYVVELGYRNCDHMVAALVGEGSGIFDALGLT-VNITKTG-
WP_216415483.1    1 --------MKKRVILTILALSM---MLAGCTDSSQQS---VVIEDIGYDPAAEVVRYE--LEELNDADKEYVVQLGYRDCDHMVPAIIGEKAGIYEALGLN-VVVTKTG-
WP_094900354.1    1 --------MTKRIIAMVLVLAMGITFLSGCTDSSKE----TAIQEVDYDPVAEVAKYK--LGELTEADKNFVVQMGYRDCDHMVPSIIGEKAGIYKALGLN-VVVTKTG-
WP_154442895.1    1 --------MKKRIIAMSLVLAMMVAVVSGCTDSSKN----TVIQEVDYDPAVEVAKYE--LGELSAGDKDFVIQLGYRDCDHMVPAIIGEKAGIYDALGLN-VVVTKTG-
WP_136713541.1    1 --------MKKRIIAMVLVLAMGAVFLVGCTDSSKE----TIIQQVDYDPVAEVAKYE--LGELSDFEKNFVVQMGYNNCDHMVPAIIGDLAGIYDALDLN-VVVTKTG-
WP_055078437.1    1 --------MKRKILSILLISSMLF-SLTACKDDSAG-----NVEEVAYDAKAEVAKYE--LPEIPEADKDLKVQMGYNDCDHMVGAIVGEKAGIYEALGLN-VTVTKTNN
WP_117606494.1    1 --------MKKRLVSVLLVMALLVPTVTACHDSSKDSQATKDVQATGYDPEEAAKKFD--FGELSDEDKNYTIEMGYNNCDHMVGAIIGEKAGIYKALGLN-VNVTKSG-
WP_173693615.1    1 --------MKKRFLAALLVVTLLVPVITACHDSSTQT-ATKDVKATGYDPKEAVKDFD--FGDLSEEEKNYTVEMGYFNCDHMVAGIIGEKAGIYKAMGLN-VNVTKSA-
WP_016219567.1    1 --------MKRRAIAALLTAAMSVSLFTGCHDSSQET-AAVDVENTDYDPEEAVKDFE--FGELSEEEKNYTIEMGYFNCDHMVGSIIGDKAGIYEALGLK-VNVTKSS-
WP_178037934.1    1 ----------KRLVAVLLVAAMSMSLLTACRDSGENK-ATVDIEQTDYDPEEEVKNFE--FGELSDEEKNYTVEMGYNNCDHMVAAIIGEKAGIYKALGLN-VNVTKSG-
WP_070088345.1    1 --------MKKRVVAALLVAAMSFSLVTGCHDSSSET-ATVDVEQTDYDPEEAVADFE--FGELNDTEKDYVVEMGYFNCDHMCCSIIADEAGIYEALGVN-VNITKSA-
WP_087378251.1    1 --------MKHMKAL----SLLLAGALAGCHDSGQEAEQ---TAEVKYDPQEAIAPYVDNLPEIPEADKDYVIEMGYNNCDHMVAAIIGQDTGLYEALGLN-VNITKTS-
WP_089610280.1    1 --------MKKRVIIGALVTTMLIAALTGCTDSSKQSN----VGQSDYDPNEEIAPYLDNLPEIPEADKEFVIQMGYNDCDHMVAAIIGETTGLYEALGLN-VVVTKTG-
WP_251861116.1    1 ------MEKRKILSLLCTL--FLGTTLIATG---------------------------CSKKTSAEKDKDYTVKLGYYNCDHMTSACVAKDAGIFDKLGLK-VSVSGNG-
WP_226098837.1    1 ------MLKLKKIALVVSLAAVMASSLIACGNKGETA------------------KTTSSNSTKVESKDDYTIKLGYYNCDHMTAACVAKDAGIFDQLGLK-VEVTGNG-
WP_073089020.1    1 ------MKRNYKWLTLLLLLVFAVS-ITGCGG-------------------------TKPAAEGGK--DDYVVKLGYYNCDHMIGAVVGEAAGIYEQMGLK-VKLFGNG-
WP_038602018.1    1 MKFIDLTKKLKKAAVLTLVATVLVTSVVGCSP------------------------KEKPATESAEKVDDYTVKLGYYNCDHMVSACVGEAAGIYKELGLN-VEILGNG-
WP_052635915.1    1 MKFFNQHRKMKKLACMLIMASLLATSAVGCSKGTE---------------------ESSVANQASQQLEAYTVNLGYYNCDHMVPACVGEAAGIFKELGLD-VKISGNG-
WP_027702358.1    1 ------MFNTRKFQVVMLTGIMGLSFLTGCSSKG------------------------DGKESSK-ANDDYAITLGYYNCDHMTAAPIGDKAGIYKELGLN-VKTLGNG-
WP_027702359.1    1 ------MSKSKKVTALMLATVIGTTVLTGCSSK---------------------------KDKAK-GDDDYAITVGYYNCDHMTAAPVGVKAGIYEELGLN-VKTLGNG-
WP_026901404.1    1 ------MKLSKKVQSLLLVSIIGGSLMSGCSNSK------------------------DAKVDSSKGSEDYTIKLGYYNCDHMTAAPVAVEAGIYKDLGLK-VETVGNG-
WP_021127843.1    1 ------MKISKKVKSLVALGAISTLVLTGCSGKD------------------------EKKETSKKADSDYTIKLGYYNCDHMTAAPVAVDAGIYKDLGLN-VEAVGNG-
WP_227829267.1    1 ------MKVSKKIQALILVSVMGASILTGCSSKP-------------------------KEEGKKAGNDDYAITVGYYNCDHMTAGPVAEAAGIYKDLGLN-VKTVGNG-
WP_006440686.1    1 ------MKLAKKLMTLGLVGVIGMTALTGCGKK---------------------------EASKEGGDTSKEITVGYYNCDHMTAAPVAEAAGIYKELGLN-VKTVGNG-
WP_099188973.1    1 -----MRRFNKKLLCILVSGMLAFS-LAGCGST----------------------KETAKNE------DDYKISLGYYNCDHMTAGPIAKDAGIYEDLGLD-VEVTGNG-
WP_038262026.1    1 ------MKLPKKLALIMSMCIVSASLLSGCGNT----------------------DDAPSEEQATKAVDDYVIKMGYYNCDHMTAAPIAKDAGIFEELGLN-VEVTGNG-
WP_200898541.1    1 -----MIKKIKNISLFSIMILLVATFISGCSNK----------------------DSKEKNITISKT-DNYKIKVGYYNCDHMVSAPIADAAGIYKELGID-AEVIGNA-
WP_114642958.1    1 -----MKK------FILGLSLLAL---IGCGE---------------------------KKE------EDYTVKLGYYNCDHMTASVIAKDAGIFEDMGLS-VDISGNG-
WP_090043125.1    1 -----MNKKLKNLLLILGVVTVSLGALSGCGKK----------------------DSAESED------NDYTIKVGYYNCDHMTAAPIAKAAGIYDKLGLK-VELTGNG-
WP_051623689.1    1 -----MNKKLKLMTTMAAISVMVLGIFSGCAK-----------------------GKEEKAS------SDYTIKIGYYNCDHMTAGPIAKSAGIYEKMGLK-VDMTGNG-
WP_040327744.1    1 -----MSKKVRRLLAVLCIVG-VTTGLVGCGSGNDKNNN--SKAVQSSSASETEVIQSIKDMKLPKLEDDYTVKLGYYNCDHMTAACIGKDAGIYEALGLN-VEVTGNG-
WP_072832005.1    1 -----MNKNLKKLTLALGILT-LSTGLVACGS--DKG------SAQSGKLDTTGIVQSIKDMKLPEIKDDYTVKLGYYNCDHMTAACIGKDAGIYEALGLK-VEVTGNG-
WP_072903777.1    1 -----MNKKLKKLLALTCCIGMMMTTLTGCTD--KKK------DNGGASGKESDIVKSIKGMNLPKIKDDYTVNLGYYNCDHMTAACIGKDSGIFEACGIK-VNVTGNG-
WP_143583134.1    1 ------MKMKKRKLLLIVLSMLIISTLVGCTKKDE---------------------GKKVGNVD----DDYEINLGYYNCDHMVGAAIGEAAGIYKDLGLN-VKLTGNG-
WP_005542131.1    1 ------MKKNYVMWGLLLMSVFTVSGCAKSQTSET---------------------GGKGESKDSVTASNYEINVGYYNCDHMVAGPVGEAAGIYKAHNLN-VKLTGNG-
WP_134213590.1    1 ------MTKIKRALPLSVLLIISLIMLAGCGGG-KKQT---------------AQNTTAG---IPEKDKDYIVNLGYYNCDHMTAACIAKDAGIFDQLGLK-VNVTGNG-
WP_012031743.1    1 ------MRRYLRFVPLFALLVILLAMLAGCGGG-TKPA---------------VQNRTAG-SGIPEKDKDYVVQLGYYDCDHMTGACIAKDAGIFEELGLK-VNVTGNG-
WP_134219405.1    1 ------MNNKIRTVMLLIVSISLLTLLTGCGGG-DKPA---------------DQNKAAANIAIPEKDKDYVVQLGYYDCDHMTGACIAKDAGIYDKLGLK-VNVLGNA-
WP_214081967.1    1 ------MKKFKKILPLLTALVVLLVLLAGCGGG-SKTS---------------NQNKAAN-TAIPEKDKDYVVNLGYYNCDHMTAACIAKDTGIFDELGLK-VNVTGNG-
WP_161822867.1    1 ------MQK-KSRLILLIVTILALSLLAGCGGGEDKTA---------------SSNKAAN-NAIPEKDKDYVVQLGYYDCDHMTGACIAKDAGIFDELGLK-VNVLGNG-
WP_031483265.1    1 ---------------MGTSYKISLSSFLSITIMMVLG------------------ALLFSANQAVADDDDYVVQLGYYNCDHMTAAPIAKEAGIFDELGLK-VNVTGNG-
WP_163340326.1    1 -----MIVANYKKLNIKEWRKMKLLRLTCVSVAMFAV------------------AFSTGLLRSAHAGDDYVVKVGYYNCDHMTAAPIAKETGIFDELGMK-VKVTGNG-
WP_041441130.1    1 ----------MKKANERLSVLILLMAFTGFTMGFPISN---------------GSAQAAGESVLPAADKDYVVKLGYYNCDHMTAAPVAKDAGIFDELGLK-VEVIGNA-
WP_028893770.1    1 ---------------MSRGRILLMGIFLA-VIVLGATS---------------SIAA-----------DDYVIQLGYYDCDHMCAAPIARDAGIFRDLGLK-VNITKGS-
WP_073613907.1    1 ---------------MKKGLQTDRRQFLKGAGVLGASA---------------ALGMGFPSLMLGKGDDDYVVRLGYYDCDHMTAAPIARDAGIFADLGLK-VEVMGNG-
WP_243440605.1    1 --MKLNKKSCFCALVLTLVCVLGTSCLSAASVEKDD--------------PVPEMVAQYDLEPLPEEDADYTVNLGYYNCDHMTAACVGKDSGIFKALGMK-VEITGNG-
WP_008522721.1    1 ----MKKSLVALALSIALGTVLTAS--SAMAVDNES--------------IVAAMVSKYSLEPLPKEDADYTVNLGYYNCDHMTAACIGKDSGIFKALGMK-VEITGNG-
WP_157150839.1    1 -MKKIVIILFMMIFILSCGGSDNASKTTEASIPKD----------------VQAIVAGYDLEELSDEDKKYKINLGYYNCDHMTAACVGEDTGIFKALGLN-VTVTGNG-
WP_147738704.1   14 VFAVLTIALFSIFAIVSCAKEEGGS---ENAIPQD----------------VQSLVSGYNLEPLSKEDSDYVINLGYYDCDHMTAACVGEDTGIFKALGLK-VSVTGNG-
WP_155812582.1   14 AALALGLAGCGSTEKGNSSASSSGSEKAEKDSEENDGE------------IVSELLSGYSLPELSEEDKNYTVNLGYYNCDHMAAASLGEATGIFKALGIT-VKTTGNG-
WP_158269839.1    1 --MYKYNIKNMTRREFLRASSFMGVALTGATVP----------------------LLTGCTKEEVAGKDNYSVQIGYYDCDHMVAAPISRDRGLLMNNGVNNFRMTGNG-
WP_093314493.1    1 --------MSMRKRKSLIMMICLSLLLAACGGG----------------------SGSGTDDGAASG---EVIEIGYFNCDMMVACPVAYHAGIYEKYGLN-TNLTMTS-

WP_248000900.1   99 -QIMEAMASGEMDVGYQGINGAIRSVNNGAPLFMAAANHLGGSYYLVVSN---DIQSP-ED-----LVGKKL-AIGSEAETGTEWCGWAEKLDIPVVTEGNYETVDMG-A
WP_248000901.1   99 -QIMEAMASGEMDVGYQGIQGAILSVNNGAPLFMAAANHLGGSEYLVVSN---DIQEP-ED-----LIGKKL-AIGNEPWKSPQWSNWAEDLGIPVMTEGNYEIVDMG-E
WP_216415483.1   93 -KIMEAMSSGELDVGYQGIEGAINSVNQGAPLFMAAANHLGGSRYLVVSN---DIKEP-KD-----LIGKKI-AVGSGAEAKPEWRKWAEELGIPVELE-NYEVVDMS-D
WP_094900354.1   95 -KIMEAMSSGEMHVGYQGIEGAIKSVNQGAPLFMAAANHIGGSRYLVVSN---DIKEP-KD-----LIGKTL-AIGSGAEVKPEWRKWSEELDIPMELK-NYEVVDMG-D
WP_154442895.1   95 -KIMEAMSSGEMHVGYQGIEGAIKSVNQGAPLFMAAANHLGGSRYLVVSN---DIKEP-KD-----LIGKTL-AIGSGAETKPEWRRWAEELDIPVELK-NYEVVDMG-D
WP_136713541.1   95 -KIMQAMASSEMQVGYQGISGAIRSVNEGAPLFMAAANHLGGSRYLVASN---EIKEP-KD-----LIGKRL-AISASSETSPEWIRWCEELGIPVELS-NYQVVEMA-Q
WP_055078437.1   94 TNIAQAMTTGEMHVGYMGIEGALNARNEGAPIIMAAANHLGGSRYLVVAN---DIKDP-KD-----LVGKKL-AITADPENNPEWIKWSAKLGLPTEKD-NYEIIEMA-Q
WP_117606494.1   99 -ETLKALTSGAMDVGYTGIEGAIRAVNQGAPFSMAAANHMGGSRYLVVSN---DITEP-SQ-----LVGKTI-SMTAEPEINPEWLTWSEKLGIPADAS-SYNIVSMG-G
WP_173693615.1   98 -ETLKALTSGAMDVGYTGIEGAILAVNKGAPFCMAAANHMGGSRYLVVSN---DIEKP-ED-----LVGKTIGSMTATPEIDPEFLTWSEKLGIPADAG-SYNIVDMG-Q
WP_016219567.1   98 -ETLKALTSGAMDVGYTGVTGAVRAVNQGAPFFIAAANHMGGSMYLVASN---EIKTA-ED-----LKGKTL-SITATPDIDPTILTWEQEIGMSSNPE-DYEIVDMG-Q
WP_178037934.1   96 -ETVKALVSGAMDVGYIGVGASALSQNP--PYFIAAANHTGGSRYLVVSN---DITEP-SQ-----IEGKTI-AMTSTPEGNAEWRAWARDYNVSFNAA-DYNIVSMG-Q
WP_070088345.1   98 -ETVNALLSGAMDVGYIFFSKNLMAEDS--PVFQASANHIGGSRYLVVNPDVIDVNDP-KS-----IEGKTI-GCLTEPETDSEWRKWSEEYGFSFQSA-DYDLVTMG-Q
WP_087378251.1   94 -QVASAMASGDMVVGYAGFGGLINNFNKGARVIMPAGSHLGGAMYLVVSP---DVESL-DD-----VVGKTL-DMGPDSQYSTFWLEICDKLGIPSDIGTYYQGSSMG-D
WP_089610280.1   97 -QVASAMGAGDMVAGYSGFSSMLKAANKGAPVFMPAGNHLGGSKYFVIRE---DINSA-KE-----IIGSRI-TITEDDMYSPYWLSYAEQVGIPKDY-TLYQGVDMD-D
WP_251861116.1   74 -KVPQAMAAGKMDVGYIGYPGLIRGLEKGSPITVAANNHLGGSMYLVVSN---KIKKP-KD-----LIGKKLGIGTKPQENE-TWLMCATKLGIPVEGK-NYNCVEFGSD
WP_226098837.1   85 -KIPQAMAAGQMDAGYIGFKGLNTAYAKGSPIIVAANNHDGGSYYLVVSN---DIKNPAKD-----LIGQPMGIGSKPEQTDYNWEQMANDAGIPVKGE-NYKGINFDSD
WP_073089020.1   75 -KVPQAMAAGQMDVGYIGTRGLMASRPKGAPIKVAANNHIGGSMYLVVAN---DIKSP-QD-----LIGQPLGIGNEPEKSEDWPV-YAKTLGIPVEGS-NYKCIEFDSD
WP_038602018.1   85 -KVPEAMAAGQMDVGYIGATGLSEASTKGAPIMIAATNHLGGSMYLVVAN---DIKKP-ED-----LYGQPVALSSDLSTDEAWLLAYGPQTGLPADSS-KFQCLEFGSQ
WP_052635915.1   88 -KVPQAMAAGQMDVGYIGTRGLMAARPKGSPIMVAATNHIGGSMYLVVSN---DIEKP-ED-----LLGQPVAIGSDPLKSESWVCGYSQDTGLPSDPS-NYECLSFGSD
WP_027702358.1   78 -KVPEAMAAGQMDAGYIGTGGLAGAIPKGAPIVVAANNHSGGSEYLIVSN---DIKDP-KD-----LIGK--KIAGDPDNDTLWHTDYGPEAGIPAKSS-EYDCIDMESD
WP_027702359.1   75 -KVPEAMAAGQMDAGYISTQGLAGAVTKGSPIVIAANNHSEGSNYLIVAN---DIKEP-KD-----LLGK--TIGANPDTNLLWKTDYGPETGLPADSS-KYDCVDMESD
WP_026901404.1   79 -KVPEAMAAGQMDAGYIGTKGLVGAIPKGSPIVVAANNHKGGSEYLVVSN---DIKEP-KD-----LIGQ--KIAADPDGDLLWTTDYGPQTGLPVDSS-KYECVNIDSS
WP_021127843.1   79 -KVPEAMAAGQMDAGYIGTKGLVGAIPKGSPIVVAANNHTGGSEYLVVAK---DVKDP-KE-----LIGK--KIASDPANDLLWTTDFGPETGLPVDAS-KYQVVNMDSS
WP_227829267.1   78 -KVPEAMAAGKMDAGYIGTKGLVGAIPKGSPITIAANNHTGGSEYLIVSN---DIKEP-KD-----LVGK--KIATDMS-DFLWTSDYGPETGLPTDSS-KYEVVNMDSD
WP_006440686.1   76 -KVPEAMAAGQMDAGYIGTRGLCGAIPKGSPIVVGANNHKGGSEFLIMSN---KYSDP-KE-----LIGK--KIATDMS-DFLWQSDYGPETGLPVDAS-KYELINMDSS
WP_099188973.1   75 -KVPQAMAAGQMDAGYIGINGLMSSSTKGAPILVGANNHKGGAYYLVVSD---DIKEP-KD-----LIGKKLGIGTKPEENSEAWIQFAKELNIPVEGS-NYECVDFGSD
WP_038262026.1   81 -KVPQAMAAGQMDAGYIGVNGLMSSAAKGAPIVIGANNHKGGAYYLVVSN---DINSP-QD-----LIGKKLGIGNEPEKNSEAWIQFAKEMDIPVEGS-NYECVGFGSD
WP_200898541.1   81 -KVPQAMLAGQMDVGYIGIDNVIKSVDNKVPMIIGADNHIGGSYYLVVSN---DIKDP-KD-----LIGQPLGIGSNPEKSS-GWVEFARELGIPVEGK-NYEAIDFASD
WP_114642958.1   62 -KVPQAMAAGQMDVGYIGVNNVFRSQLKGAPLVIGSNNHKGGSYYLVVAN---DVEKP-ED-----LIGEKLAIGTHPERST-GWTQIAKSLNLPVDGK-NYQGIDIGSS
WP_090043125.1   76 -KVPQAMAAGQMDAGYIGIGNVLNSPKNNVPLVIGANNHEGGSYYLVVSN---DINEP-KD-----LLGKTLGIGSKPEEDL-SWATMAKELNIPVEGS-NYDAINMNSI
WP_051623689.1   75 -KVPQAMSAGQMDAGYISISGALNSPKNNVPLVLGANNHEGGSYYLVVSN---DVKDP-KN-----LLGKTLGIGSKPQEDL-SWQTISKSLSISQEGK-DYDAINMNSD
WP_040327744.1  101 -KVPQAMSAGQMDAGYISFQNVTRGINKGTPITIAANNHDGGSYYLVVSN---DVKEP-KD-----LIGQKLGIGTEPEKSW-SWRQMAKEMNIPVEGK-NYEGINFDSD
WP_072832005.1   95 -KVPQAMSAGQMDAGYISFQNVTRGINKGTPITIGANNHEGGSYYLIVSN---DVNEP-KD-----LVGQKLGVGTEPEKSW-AWRQMAKDMNIPVEGK-NYEGINFDSD
WP_072903777.1   96 -KVPQAMSAGKMDAGYIGFQNIAKGFLVGTPITIAANNHTGGSYYLVVSN---DITDG-KQ-----LVGQTLGIGPEPEKGS-SWRVMSKTLDIPTEGK-NYNGISFDSD
WP_143583134.1   78 -KVPEAMAAGQMDAGYIGNRGLIAANGKGAPILIGANNHIGGSMYLVVAN---DIEKP-EE-----LYGQKVAIG--STQDEGWIAGYSKLLNLSTDEK-DYDIVNISSD
WP_005542131.1   82 -KVPQAMAAGQMDAGYIGARGLVAANGEGSPIVYAANNHIGGSMYLVVAN---DIEKP-ED-----LYSQKVAMPD-PSTSESWLAGYAQTLNLSADPA-KYELIQTGSD
WP_134213590.1   84 -KVPEAMAAGKMDVGYIGTEGLIRAFQKGSPIFVAANNHLGGSFYLVASN---EIK-DAKG-----LVGKKVAIG---DVTDSYWVRMANDIGIPFDEK-SYQNFDMA-D
WP_012031743.1   86 -KVPEAMAAGQMDAGYVGNTRMMRAFLQGSPIIVGANNHIGGSHYLVVSN---SIK-TPQE-----LVGKKLSIGTDPEKNNANWITWANQFGIPVEGK-NYEVFNMA-D
WP_134219405.1   87 -KVPEAMAAGQMDVGYVGNSRMMRAFLQGSPIVVGANNHIGGSHYLVASN---DIK-EPKD-----LLGKKLAIGTAPEKNNPSWIVMAERLGLPIEGA-NYEVLNMA-D
WP_214081967.1   86 -KVPEAMAAGQMDAGYVGNSGSMRAYIKGSPIIVAANNHVGGSYYLVASN---DIT-DPQQ-----LIGKQLAIGTNPEKTSISWINMAKELGLPVEGK-HYEVLNMS-D
WP_161822867.1   86 -KVPEAMAAGKMDAGYVGNSGTMRAFLKGAPILVAANNHIGGSYYLVASN---DIK-DPQE-----LIGKKIALGTDPEKNNISWIGMAKELGLPIEGK-NYEVFDMS-D
WP_031483265.1   76 -QVPQAMAAGRMDAGYIGIHGLMRARLKGAPIFVSANNHLGGSYYLVVSN---DIK-TPQD-----LIGKKLSLGSNPTKRSASWVGFTAANNIPVDPA-KYEVFAMA-D
WP_163340326.1   86 -SVPQAMAAGQMDAGYVGIHGLMRAHIKGAPIFVSANNHVGGSYYLVVSN---DIK-DPQQ-----LVGKKLALGTNPNKRSVSWVSFVKENNLPKDPS-NYKIFAMS-D
WP_041441130.1   84 -KVPEAMAAGQMDLGYIGFERTVRAYLKGCPIFIGAMNHLGGSYYLVLRN---DIY-EQYQKDRRVLLGKKLALGTEPEKNSPEWVTFTGKIGLPPDGK-NYETFNMS-D
WP_028893770.1   67 -NVLPAISAGKMDVGYGNYISLARAHLKGSPAFVAAHNHTGGAFYLVMAN---NFKGKPKD-----LIGKRLALGMAPEETLPWWHDYAKANSIPVKAD-GYQNFIMK-D
WP_073613907.1   79 -KVPQAMAAGQMDVGYIGFVGMTKAILKGSPMVAVAHNHKGGSMYIVARH---GIE-KPED-----LLGKKVALGAKPETNNGWWVTYAMQNNIPKEAE-HYQNFVMT-D
WP_243440605.1   93 -KVPEAMSAGRMDMAYAGWTTTLRAVQVGTPLFIAAENHTGGAEYLVCSF---NIKKP-ED-----LIGKKVAAGGDPANS-MNWMEWTDQLGIPNDIE-KYENYSMS-D
WP_008522721.1   89 -KVPEAMSAGRMDMAYAGWTTTLRAVQVGTPLFIAAENHTGGAEYLVCSF---KIKTP-HD-----LIGKKICAGGDTAND-LNWQEWTAQLGIPNDIK-NYENYSMS-D
WP_157150839.1   92 -NVPEAMSAGQMDMAYAGFTTTLNAVKNKVPLFIAAENHTGGAEYFVVSK---DIEKP-ED-----LLNKRISMGANAETVNLNWAEWTEQLNIPRDTK-YYENFSMS-D
WP_147738704.1  103 -NVPEAMSAGKMDVAYASFNTTLNAVKNKVPLFIAAENHTGGAEYFVVAK---NINKP-ED-----LLSKRISMGANAETENLNWAEWSADLNIPRDTK-YYENFSMS-D
WP_155812582.1  110 -NVPEAMSAGQMDMAYCGWTTTLSAVQKHVPLFIAAENHTGGSEYLVVSN---KIGKP-EE-----LIGKKISVGTEPEKNNLNWAEWTKQLNIPEDIT-KYDNQEMS-D
WP_158269839.1   86 -QVPQAMTAGHMDVGYVGTTTIERAFRNGAPIRVVANNHRGGSWYLVAAN---HIKEA-KD-----LIGKRVSLG---AHPNPFWCNCATQVDIPWQFE-HYDVYVMT-D
WP_093314493.1   76 -DIGVLMAAGQMDVGYMGSNAVNNALNQGTPIKMTAFNHLGGSYYLVGSN---EIETA-QD-----MVGKRINMGPDAHEVNPEWRIIAHDAGIPYDPA-EYEAYSMS-Q

WP_248000900.1  197 TDALIAMKAGQIDGFATUDPFASQAEFEGIGHVMATDWGYKGSKLGVDSDSNDLDNWGICCVYAMSTNFYEEHPELAKRLVLAHALSVKYVYTHPYNAAMIFADAYGTKP
WP_248000901.1  197 TDALLAMKAGQVDCFSTUDPVASQAEFEGIGHVMATDWGYKED----GTDGNDLDNWGVCCVYAMSNNFREEHPELAKRMVLAHALSIQYIYQHPYNAAMIFADAYGTKP
WP_216415483.1  190 KDKVFALKAGQLDAFSACDPYGSLVEFEGFGRIMATGWGAHVSED-------MEEGWGICCIYAMNNDFLEEHPELAKRLVLAHSLAIKYLYEHPYNAAMMFADGFGTVP
WP_094900354.1  192 KDKLFALKAGQLDGFSCCDPYASQAEFEGIGKIMATGWGAHISDD-------LETGWGMCCIYCMNNDFLEEHPELAKRLVLAHALAIKYLYEHPYNAGMMFAEGFGVQP
WP_154442895.1  192 KDKVFALKAGQLDAFSACDPYGSMVEFEGFGRIMATGWGAHVSDN-------MDEGWGVCCIYCMNNDFLNDHPELAKRLVLAHALSIKYLYEHPYNASMMFAEGFGTNP
WP_136713541.1  192 ADALFAIKAGQLDAFVCCDPYASQAEFEGIGHIMGTGWGAFYPEDE--TNTGAVEDWGICCIYAMSDDFYEQCPELSKRLVLAHALAIEYLYEHPYNAGMMFAEGFGVDP
WP_055078437.1  192 KDGPMALKSGQISAFACCDPFASQCEYQGLGKVVAVGWGGLIDETKEEND--HEHGWGMCCIYAMNENFKNEHPEVAQRLLVAHALSLKYLYDHPYNAAMMFADGFGTAP
WP_117606494.1  196 QDAMFALKAGQIDAFTCCDPYASIAEFEGFGHIMGIGWGGANVDSDAT-----ADTWGLCCIYAMSNDFKEKHPELARRLVYAHEMALEYMYTHPYNAAMMFADGFDVDP
```

```
WP_216415483.1  190 KDKVFALKAGQLDAFSACDPYGSLVEFEGFGRIMATGWGAHVSED-------MEEGWGICCIYAMNNDFLEEHPELAKRLVLAHSLAIKYLYEHPYNAAMMFADGFGTVP
WP_094900354.1  192 KDKLFALKAGQLDGFSCCDPYASQAEFEGIGKIMATGWGAHISDD-------LETGWGMCCIYCMNNDFLEEHPELAKRLVLAHALAIKYLYEHPYNAGMMFAEGFGVQP
WP_154442895.1  192 KDKVFALKAGQLDAFSACDPYGSMVEFEGFGRIMATGWGAHVSDN-------MDEGWGVCCIYCMNNDFLNDHPELAKRLVLAHALSIKYLYEHPYNASMMFAEGFGTNP
WP_136713541.1  192 ADALFAIKAGQLDAFVCCDPYASQAEFEGIGHIMGTGWGAFYPEDE--TNTGAVEDWGICCIYAMSDDFYEQCPELSKRLVLAHALAIEYLYEHPYNAGMMFAEGFGVDP
WP_055078437.1  192 KDGPMALKSGQISAFACCDPFASQCEYQGLGKVVAVGWGGLIDETKEEND--HEHGWGMCCIYAMNENFKNEHPEVAQRLLVAHALSLKYLYDHPYNAAMMFADGFGTAP
WP_117606494.1  196 QDAMFALKAGQIDAFTCCDPYASIAEFEGFGHIMGIGWGGANVDSDAT-----ADTWGLCCIYAMSNDFKEKHPELARRLVYAHEMALEYMYTHPYNAAMMFADGFDVDP
WP_173693615.1  196 QDACFALKAGQIDAFTCCDPYASIAEFEGFGKIMGIGWGAADVDAEST-----ADTWGLCCIYAMSNEFKDNHPELARRLVYAHQLAIKYMYEHPYNAAMMFADGFDVDP
WP_016219567.1  195 QDAMFALKAEQIDAFSCCDPYASIAEFEGFGHIMAIDWAAPDISSDSE-----YDDWGLCCVYAMNTDFSKNCPELARRLIFAHSLAIKYMYEHPYNAAMMFADGFDVDP
WP_178037934.1  191 QDAMFALKAGQIDGFVCCDPYASIAEFEGFGHIMFTSWGINMPEDGSLP---GSDDWARCCALAMNKDFAEQHPELARRLVYAHMLSIKYLYEHPYNASMMFADGFDCDP
WP_070088345.1  196 QDAMFALKAGQIDAFTCCDPYASIAEFEGFGKIMAISWGANL-EDGELL---TDENAGTCCSYAMSKEFREKCPELSRRMIYAHMLAVEYLYTHPYNAAMMFADGFDVDP
WP_087378251.1  192 EDAMMALRAGQLDIMTCCDPYASIAEQEGFGKILATAWHADVKEDK-------STGWGIHCGYYINADFAEAHPELTTRLVLAHCLAIKYMYQHPYSAAEMFAETFGTST
WP_089610280.1  194 NDAIVAMKAGQLDGFMCCDPFASMAEMEGIGKIIGTEWGGHISEDL-------ESGWGLHCGYFINSDFAKAHPELTTRLVLAHCLSIKYMYQHPYNAGMMFADDFGVDP
WP_251861116.1  172 KDAYLALKAGKIDGFVCCDPWGSMAEYEKTGNIIATADKTVGKDKW-----------GNCCVYSMNTKFEKEHPELAKKMIQAHVEAMKYCYEHPIKAAKIFAKNYKVPE
WP_226098837.1  185 QAKYLAMKTGKIKGYVCCDPWGSMAEFEKTGRILSTHLMTG--KEM-----------GNCCVFSLNKKFAKEHPELAKKMVLAHTKSIEYIYTHPLKAAKIFAKNYNVPE
WP_073089020.1  173 QAAYLALKTGQIKGFTCCDPWASMAEYEGTGYIMAT--YMEMMTGDE-------KQMGSCCVYSMNENFKKEHPELARKMVQAHVEALKYVYTHPIKSARIFAEYYHVPE
WP_038602018.1  184 ADKYLALKTGQIKAFTCCDPWGSMAEFEGTGKIMGT--YLEMDG-----------QLGVCCVYSMNKKFKEEHPELAKKMVLAHTKSMEYVYTHPLESAKIFAEYYKVPV
WP_052635915.1  187 ADKYLALKSGNIKAYTCCDPWGSMAEYEGTGKIMST--YMEMDG-----------AMGVCCAFAMNSNFEKEHPDIARKMVLAHTKSIEYIYTHPVKAAKIFADYYNVPE
WP_027702358.1  175 KDAYLAFKTGKIKAFTSCDPWGSVAEFEGTGKIIASTQYKEKSTG----------KQYNCCSFSLNKNFIKDHPELAKKLVLAHTKSVEYIYTNPVKAAKIFAEDYNVDE
WP_027702359.1  172 KDAYIAFKTGKIKAFSACDPWGSVAEFEGTGKIIASSKYHSESTG----------NEYNCCSFSLNKNFIKDHPELAKKLVLAHTKSIEYIYTNPIKSAKIFAEDYNVDE
WP_026901404.1  176 KDAYLAMKTGQIKAFSTCDPWGSVAEFEKTGKIIATTKYKEESTG----------KEYNCCSFALNTNFIKEKPELAEKLVLAHTKAIEYIYTNPVESAKIFAKAYSVDE
WP_021127843.1  176 KDAYLALKTGQIKAFSTCDPWGSLAEKEGVGKIISTTQFKEKNTG----------KDYNCCSFSLNTNFVKEHPDLAKKMVLAHTKAIEYIYTHPVKAAKIFAKNYNVDE
WP_227829267.1  174 KDKYLALKTGKIQAFTSCDPWGSVAENDGVGKIIASTQYKEKAND----------KEYNCCSFSLNKNFIKEHPDLAKQLVLAHTKAIEYIYTNPAEAAKIFAKYYNVEE
WP_006440686.1  172 KDAYLAMKTGKIDAFTACDPWGSVAEFEGTGKIVASTQYKEKDG-----------TEYNCCSFALNTNFVKENPELAEKLMLAHTKAIEYIYTHPAQAAKIFAEYYDVDE
WP_099188973.1  174 KDAYLALKTGQIDGYTCCDPWGSMAEYEKTGHIL-AVNSKL-EGEW-----------GVCCTFSLNKNFVKDHKELAEKLVLAHTKAMEYIYTNPVKSAKIFANNYDVPE
WP_038262026.1  180 SDKYLALKSGQIDGYTACDPWGSMAEYEGTGKIL-AVNSLL-EGEW-----------GVCCTFSLNKNFTREHPELSKKLVLAHTKAMEFIYTNPAESARIFAANYNVPE
WP_200898541.1  179 KDAYLAFKTGKIKGFIACDPWGSMAEYEKTGKIL-ITSSKL-EGQW-----------GDCCVLSINKDFAEKHPELAKKVALAHTKAMEFVYLNPLKTAKIFAKTYMVPE
WP_114642958.1  160 TDALTALRMGEIKGFTACDPWASMAEYQGIGKIMGTENKLL-NDKW-----------GVCCTYTLNKKFIEEHRDLAKKMILAHAKAIEYIYENPLKSAKIFADNYNVPL
WP_090043125.1  174 QDSYVALKSGQIKGYTACDPWGSMAEYEKVGKIM-KEYVGP-EGDM-----------GLCCAFTLNKNFVKEHRKLAEKLVQAHVEAIKFMYEHPVQSSKMFAEAYKVPE
WP_051623689.1  173 KDAYLALKSGQIKGYTACDPWASMAEYENTGKIM-YEYKGP-DNKP-----------GICCAFTLNKNFISEHKDLAEKLVKAHVEAVKYMYEHPVKSAKIFAETYMVPE
WP_040327744.1  199 QNAYMALKTGQIKGYTACDPWGSMAEYEKTGKIMKIFDSVD-GET------------TICCVFALNTNFMKEHEEVAKRLVLAHTKSIEYLYTHPIESAEIFAKNYSVPK
WP_072832005.1  193 KSAYIALKTGEIKGYTACDPWGSMAEYEKTGKIMNVFDSVD-GEK------------TICCVFALNTNFVKDHPEVAERLMLAHTKSIEYLYSNPYKAAKIFAENYSVPE
WP_072903777.1  194 QSKYLALKTGKIKGYTACDPWGSMAEYEKTGKIFAMDDRIK-GEQE-----------GECCVLSLNTNFAKAHPDIAKRLILAHSKSIEFIYTHPLEAANIFANNYGVPL
WP_143583134.1  175 ADAYMALATGQIKAFTSCDPWASMAVYEGKGKIMAT--YMEMDD-----------KMGICCAFSLNEKFVEEHPELAEKLLTAHQRSIEYVYEHPHEAAKIFAKYYNVPE
WP_005542131.1  180 SDSYLALSTGQIKAYTCCDPWGSMAEFNKVGKIMGT--YYEMDG-----------AMGICCGFALNKNFIKEHRDLAVELLKAHQDSIKYIYEHPKKAAEIFSKYYNVPY
WP_134213590.1  179 KDEFFAVKAGQLDGYLCCDPWGSMAEYDKSGHIVKAVGKLPT------------GDWGSCCVYSMNRNFAKDHPEIAAKMLLAHTKAIEYVYTHPVRSSEIFAANYYVPQ
WP_012031743.1  184 KDEYFAMCAGKLDGFAACDPWGSMAEYEKTGHVLAASGKLPN------------GEWGECCVFALNKKFASEHPELAKLMILAHTRAIEHMYLKPVRSAEIFAQNYKVPM
WP_134219405.1  185 KDEYFALKAGKLDGYAACDPWGSMAEFEKTGHVLAAMDKLPS------------GEWGECCAFNLNKNFATEHPELAKLMVLSHTKAIEHMYLKPVRSAEIFAENYKVPM
WP_214081967.1  184 MDEYFALKAGRLTAYSTCDPWASMAEYEKTGHVLAVDGKLPS------------GEWGECCAFHLSKQFASEHPELARLMVLAHSRAMEHIYTKPVRSAEIFAKNYQVPE
WP_161822867.1  184 KDEYFALKAGHLDAYTACDPWGSMAEYEKTGQILAIDGKLPS------------GEWGECCAYHLSKNFAAEHPELAKLMVLAHTRALQYIYTKPVHSAEIFAANYNVPL
WP_031483265.1  174 KDEFIALKTHSLDGYLTCDPWGSMAEYDGCGWIMETFSKLPS-T----------NDWGVCCVLAMSNEFGKEHPELAKKMIQAHVKALEFIYTQPKRAAEIFAKAYSVPE
WP_163340326.1  184 KDEFLALKTGKLDGYLTCDPWASMAEYDGSGHVMKLFDKLPS-T----------QDWGVCCVLAMSRKFAKEHPDLAVKMIQAHMKALQYIYTHPKRSAEIFAKVYSVPE
WP_041441130.1  187 KDEFLAMKTGKLDGYTTCDPWGSMAEHDRSGRIVPEFTITRMPS----------GKWGACCVLSVNRNFAEAHPELARKMILAHSKAIQFIYTHPLRTAEIFARNYSVPL
WP_028893770.1  166 QDEYLAIKTGKLDGALMCDPWGSMIEYEKIGHIMHTFDPLPG------------DKEGICCVYSMSKEFSVKHPDLAVKMLLAHSRALQLVYTHPLKSARIFAASYSVPL
WP_073613907.1  177 RDEYLAFKTGELDAFWCCDPWGSMAEYENTGKIIRTFDVFPN------------GTWGICCAQVMNRNFVNEHPEIAQKMVLAHVRAIQYIYTQPIKAAHIFSESYHVPY
WP_243440605.1  190 ADAYFSMVAGKLDAFTCCDPWGSMAEHEKTGWVMVRQNTDRK------------NGHGTCCKVCMNYNFAKKHPKLASRMLLAHTLCVQYMYQHPYKAARIFAENYNVPL
WP_008522721.1  186 ADAYFAMVAGKLDAFTCCDPWGSMAEYEKTGWVMVRQNTQRE------------SGHGTCCKVCMNYNFAKKHPKLASRMLLAHTLSLQFMYLHPYKAAQIFSANYNVPL
WP_157150839.1  190 SDAYFAFKLGQLDGYGACDPWGSMAEYENTGKILKRQNTDREI-----------DGHGTCCKVCMNYNFAKAHPKLAERILLAHAMSIEYMYLHPYKAAEIFAENYNVPL
WP_147738704.1  201 SDSYFAFKLGKLDGYGACDPWGSMAEYEDTGKILVRQNTDREI-----------DGHGTCCKICMNYNFAKAHPKLAERILLAHAISVEYMYLHPYKAAEIFAKNYNVPL
WP_155812582.1  208 SDEYLALVAGSLDGYICCDPWGSMAEYSKAGHVMVRQDTDRP------------SGHGTCCKVAMNYTFSKAHPELAKRMLLAHTLCIQYMYEHPYRAARIFSAYYNVPV
WP_158269839.1  181 RDEYVALMAGHLDGYTCCDPWGSMAEYSDVGWIVATNHLLQDGS------------WGACCDLVMRDAFIEQHRELAKLCLKTHIDALMFVYEQPYKAAETFASDFKVPL
WP_093314493.1  174 RDAFFALEIGELDGVLVCDPFMSLAEHRGIGHELAVSRMLPDGD------------WGFKTLLIMHDDFIQERPEEAEKMLLAHVEAIQYLYTNPIESGKIFAEVFDVPE

WP_248000900.1  307 EVALNTIYMKTVKEGRTLTWQFS-EKNLENLVANWNQYDT--IPEEDRIKINDL--QSFMSTDLIEECGVEDFHDFIENE-VEPLYPTGISFEDWLAKAKEIDGITTNEY
WP_248000901.1  303 EVALNTIYMKTVKEGRTLTWQFS-EKNLENYIANYTKFDH--IPEEDIPNINDV--EKFMSTDLIEECGVEDFSDFIANT-VEPMYPTGITFEDWLAKAKEIDGITTNEY
WP_216415483.1  293 EVALKTVYMKTVAEGRTITWQFS-EENLNNYIDYFYDYE---IAEDNIPNINDI--EKFMSTDLLETCGIEDFDTFIEETNIDKEFPVGMSYDDWLAKAKAIDGITE---
WP_094900354.1  295 EVGLKTVYMKTVSEGRTITWQFS-QENLDNYIQYYYDFD---IPEEEIPNIADI--EKFMSTNLLETCGIEDFNTFIEEK-INKEFPIGLSYEEWMTKAKTIDGITEQ--
WP_154442895.1  295 EVGLKTVYMKTVAEGRTITWKFS-EENLDNYIKYFYDFE---IPEEEIPNINDM--QQFMSTDLLETCGIEDFNTFIKEN-IDANFPIGDTYEEWLAKAKAIDGIGESTA
WP_136713541.1  300 EVGLKTVYMKTVAEGRTITWEFT-KQNLDNYIQYYYDYN---IPEKDIPNINDI--EKFMSVELLASSGVGDFSKFIEEE-IDELFPIGDTYEEWLAKAKTIDGIEETTA
WP_055078437.1  300 EVGLMTLYMKTIAEGRTLTWEFN-QDNIDNFVKEYDEYN---IDEKFRPKLDKL--DEFVNLNFVKESGIEAFGDFTKNDKIDEKFPVGISFEDWLKKAKEIDGIENEVG
WP_117606494.1  301 YVALRTIYMKTVAEGRTITWHFS-EKNIENFENYYTQYP--QIPEEEIPRVSDV--SKFMTTDISKDAGVDDFDEFIKKN-VDDKFPLGMTFEDWYNEAKKIDDISDDEA
WP_173693615.1  301 YVALRTIYMKTVAEGRTITWHFT-EQNIENYEHYFTQYP--QIPEEEIPKVNDV--SKFMSTEISEDAGVDDFESFIKES-IDPVMPIGMTFEDWYNKAKEIDNISEKEA
WP_016219567.1  300 YVALRTIYMKTVSEGRTLTWKFS-DPNVDSFLEYYTQYS--QIPEEEIPLINDR--NTLISSEVLEAAEIEDFQTYIKDV-VDPIMPLGTTFEDWYEISKEIDGISDEDA
WP_178037934.1  298 YVGLRTVYMKTVAEGRTITWHWS-QANLQNDEDFDTQFTNPSITETDISYVSIFEDSLKRATELADSAKLDDFDEYIKEK-VDPISPLGITFEDWYEHAKTIDGISDEDA
WP_070088345.1  302 YVGLRTVYMKTVAEGRTITWQWS-EQNMINDEQFERQWTNPSVPEEDIINASDAQSSLELSGQIFEDAGIKDFNEFIAEE-VDDIFPLGMTFEDWYNAAKEVDGISDEDA
WP_087378251.1  295 DVGLRTMYLKTNAEGRTLCWTID-ESNMENYENWYDQLG---VPEEERTLVTDV--DGFMDLSWLDKCGIEDFQTFIDEEKIDEVQPIGMSYADWLYMAEEIDGIDHEST
WP_089610280.1  297 AVGLRTMYLKLCAEGRTLSWDVD-ETNIKNYTDYWYSFG---IPEEEIPYVKDI--PSFLDLSYMEACGIESFESFMEDNKINEKFPKDDPYLTWLEKAESIDGIDHSST
WP_251861116.1  271 EVGLMTIYKKTVGEGRTLTWKMD-DAYFKTEIDTLMKYKL-------IEEKPDY--DKLISKKIYEEAKVEDFDKFIKEK-VDSVFPVGMKYEQWKAKSMKIDK------
WP_226098837.1  282 EAALLTVYKKTVGEGRTLSWKVD-PKKFKHEMEFEISKGL-------YSDSNKY--EDIVITKYLDESGADDFDKFIKEK-VNAVFPLGMSFENWKAKSNEIDK------
WP_073089020.1  274 EVALRTIYKKCVAEGRTLTWKID-EGQMRFAMKVYQNEKL-------MDEVPAY--EDVIDIEFYKSCNLEDFDKFIKEK-VDPVFPVGMSYEDWKKKAKEIDA------
WP_038602018.1  281 EVAEMTIYKKTVGEGRTLSWKMD-EQAHKHALEVYKKFNI-------MENVPNY--DDIAMKDILKESGAQDFDQFIKEK-VDPVFPLGMSFEDWKARALEIDPQ-----
WP_052635915.1  284 EVAMMTIYKKTVAEGRTLTWKID-DTQYAHAMEGYKKYDL-------VDELPEY--DDVVKKNILAESGAKDFDEFIKND-VDEIYPVGMSYDDWKAKAMEIDA------
WP_027702358.1  275 EVALMTIYKKTVGEGRTLTWKIT-GEEYKNNLKMYKDYNK-------MDKVPSY--DEVVDKSLLDSCGADDFDKFIEEK-VDPVFPKGMSYADWKAKAIELDKQ-----
WP_027702359.1  272 EVALMTIYKKTVGEGRTITWNIT-GNEYKNNLETYKEYKS-------LDEVPTY--DQIVDKTLLESSGADDFDKFIKEK-VEPVFPEGMSYTDWKVKATELDKQ-----
WP_026901404.1  276 EVALMTIYKKTVGEGRTLTWKIT-GDEYKNNLALYKEHKA-------LEEIPTY--EQVVNDKLLKSCGADDFDKFIKEK-VNPVFPEKMNYEDWKAKAMEIDGKN----
WP_021127843.1  276 EVALMTIYKKTVGEGRTLTWKID-GDEYKNNLQMYKDNKS-------FEEIPDY--NKVVNDELLKSCGADDFDKFIKEK-VNPVFPEGMSYDQWREKAMEIDGQK----
WP_227829267.1  274 EVALRTIYKKTVGEGRTLTWKVT-GEEYKNNLQMYKDLKA-------LDDIPKY--EDTIDTSLLDSCGADDFDKFIKEK-VNPEFPEGMSYEDWKAKVL----------
WP_006440686.1  271 EVGMMTIYKKTVGEGRTLTWDIGEGEGYKNNLKMYKDLKA-------LEEIPSY--EEVVDDSILKSCGADDFDTFIKEK-VDPVFPQGMDYETWKEKALSIKE------
WP_099188973.1  271 EVALMTIYKKTVEEGRTLTWKIN-PEEINREMKWKKEHGF-------ADESLKY--EDIVDASYLENSGADDFDQFIKEK-VDPVFPVGMSYEDWKEKAFEVDK------
WP_038262026.1  277 EVALMTLYKKTVKEGRTLTWVVN-PDEINREIDWKKEHGF-------IDESVTY--DTVVDTSFLEASGADDFDTFIKEK-VDPVFPVGMSYEDWKAKVQELDKK-----
WP_200898541.1  276 EVGLLTVYKKTVKEGRTLSWKIV-PEQIKHELSWKQENGY-------VPNTVTY--EELVDSKFIDESGADDFDKFIKEK-VDPVFPVGMSYDEWKTKAKKLGK------
WP_114642958.1  258 EVAVSTIYKKTVGEGRTLTWKLD-PQEIKNELEWAQESGF-------TPKEVKY--ENIVDESIYNEADVPDFDKFIAEK-VEKNFPLGMNYDSWYAKAQELDK------
WP_090043125.1  271 EVGLMTIYKKTATIGRTITWKMD-EEDAKREIKARQDIGS-------VDKNVEY--EDVVDNQLLDELKVEDFDQYIKEK-VDPIFPLGMSYEDWKTKANEIDK------
WP_051623689.1  270 EVALMTIYKKTAEIGRTITWEVK-PDNIRREIKARQEVGS-------IDGNVKY--EDIIDKDFLESVKVDNFDDFIKDK-VEKNFPLGMKYEDWKVKATEIDK------
WP_040327744.1  296 EVALLTVYKKTVEEGRTITWKVN-ESAVDKRLKENVEHG--------DYENVDK--KALIDNSLLKECGADEFDTFIKEK-VDKIFPVGMSYDDWMAKAKEIEGKN----
WP_072832005.1  290 EVALMTVYKKTVEEGRTITWDIN-DAAIDKRLTENISHG--------DYEEVDK--NQLVQTDLLDKCGKDDFNTFIKEK-IDPVFPTGMSYDEWLKKAKEIAGE-----
WP_072903777.1  292 EVGLRTVYKKTVGEGRTLTWKLN-NEAIQRMMDENISHG--------DYGKVTV--EQIVNHDYINDSKVDDFEKFISEK-VDSVFPEGITYEEWLEKAKTREPEIKTVS
WP_143583134.1  272 EVAIMTIYKKCVEEGRTLTWEIE-EESFQHAYDVYKRYDL-------IETLPEF--KDLIAYDIYNAAKLGDFDEFIKEK-VDSVFPIGMSYEDFKKKAMEIDE------
WP_005542131.1  277 EVSLMTIYKKCIEEGRTLTWQLN-EEEFTHAYDVYKKFNL-------IKNLPDM--EDVIERDIYNEANLDNFEEFIKNN-VDKNFPVGMTYEEFEKKANEVDP------
WP_134213590.1  277 EVALMTIYKKTVDEGRTLTWKVN-PKNFQDEVDYILGLKT-------LETAPTL--DQFIDTKLLEGCGADNFDTFIKEK-VDPVFPVGMPYEEWKKKAYELEGKKA---
WP_012031743.1  282 EVALMTIWKKTVAEGRTITWKMD-RNYWKNQIDLELGYKT-------IDAAPNI--DEFLQPQYLEECGADDFDIFIKEK-VDPVFPVGMSYEEWKKKAYELEGKKA---
WP_134219405.1  283 EVALMTMWKKTVAEGRTITWKID-LNNWQRQIDMELGFGT-------LDKAPNV--NEFIQPQYLNESGADDFDTFIKEK-VDPVFPVGMSYEEWKTKAYKLEGKTA---
WP_214081967.1  282 EVALMTIWKKTVAEGRTLTWEID-LNNWQRQIDMELAAGT-------LDAAPKL--DEFIQPQYLRDSGADNFDTFIKEK-VDPVFPVGMPYEEWKKKAFELEGKTA---
WP_161822867.1  282 EVAFMTIYKKTVAEGRTLTWTID-YDYWQRQIDLELAEGT-------LEAAPNV--DEFIQPQYLNECGADDFDTFIKEK-VDPVFPIGMSYADWKAKAYQLEGKKA---
WP_031483265.1  273 EVALMTIFKKTVAEERTLNWEVD-KGKIQDEIDYNVSIGT-------LDAAPPL--EEFVNVEFLNAAKVDDFKTFIKEK-VDPIFPMGMTYEQWKAKALEIH--AQRG-
WP_163340326.1  283 EVALMTIYKKTVAEERTLNWEID-TKKIQAEIDYNLDMGT-------LDKAPPL--EKFVNTQFLDTANTDDFETFIKEK-VDPIFPMGMSYSDWKAKAIEID--KPQA-
WP_041441130.1  287 EVALMTIFRKTVAEDRTMRWDIH-EENVREAVAWNLGTKT-------LDQAAAT--NDYTDTRYLREAGTEDFERFIREK-VDPVFPLGMSYPDWKKKAYAVEGKAFQAA
WP_028893770.1  264 EVALMTIYKKTNSEGRTMTWDIH-RKGFENQTRHYMDIGY-------LTAMPDM--KRLVNPEIYDKAKPLDFARFIKEK-VDPVFPLGMSYADWKKKVMARGL------
WP_073613907.1  275 EVGLMTIYRKTVGESRTLRWNLE-EEAYNETVSHNLSIGI-------FPKAPKY--EEIVNTTFLENKQIADFDDFISSE-VDTVFPQGMPYADWKKKATELNV------
WP_243440605.1  288 EVGLMTLYKKLNEEGRTISWKLN-RQDMQNQLDTMRHYHV-------RDDINTVNLDDYIDLSYFEASGAIDFDRFIQEK-VDPVFPTGMSYKDWRAKAVEVDG------
WP_008522721.1  284 EVGLMTLYKKLNEEGRTLSWKLN-RQYMQNQLDTMKKYHV-------RDDINSVNLDDYIDLTYFNASGAIDFDKFIAEQ-VDPVFPLGMSYKDWRAKAVAVDG------
WP_157150839.1  289 EVGLMTLWKKLNEEGRTITWKLN-REYVHNQMKTMRRLGV-------RDDINSLDIDDYIDLSYFDNCGAKDFDVFIKEE-VDPIFPLGMSYEDWRAKAVEIDN------
WP_147738704.1  300 EVGLMTLWKKLNEEGRTITWKLN-REYVYNQMKTMRRLGV-------RNDINQLNIDDYIDLSYFNNSGAKDFDKFIEEE-VNPIFPLGMSYEEWRAKAVEVDN------
WP_155812582.1  306 EVALMTYWRKLVDEGRTIRWDLN-LDYMRNQLATMREYGI-------RDDINTVNVEDYVDLSYFDQCGAKNFEDFAKEK-VDPVFPEGMSYEDFKKKALAIDHAEGADL
WP_158269839.1  279 PVAIKTIWRKTNAEGTSMTWRHDVSEKERQFAYSKS-LGL-------PEFQETAPPAEWMDDTLLKEIGAPDFYEFIEKK-VKPYFPDRMSLEDWTDVATKRHTI-----
WP_093314493.1  272 EVGFRTIFQKTVDEGRTMTWVIDDERVERYYDMTQNYFDM-------LEWQDFTPQEEWVYHDLQNQLDLPDFDAFITEK-VDPYYPVGMSYDDWLEKAMELYES-----

WP_248000900.1  411 DDIAEITMND------------- 420
WP_248000901.1  407 DDAAEITMND------------- 416
WP_154442895.1  398 SK--------------------- 399
WP_136713541.1  403 SAK-------------------- 405
WP_055078437.1  404 NDIKIPEVYKEQK---------- 416
WP_117606494.1  405 VDISKTATSYLNKDLKDRQAYE- 426
WP_173693615.1  405 VDIADTATSYLNKDLKDRETYE- 426
WP_016219567.1  404 VDISDTATPYLNENLDERTSN-- 424
WP_178037934.1  406 VDISKTATPYLNENVEGTDKK-- 426
WP_070088345.1  410 VDISDTATPYLNENVEE------ 426
WP_087378251.1  399 VGKTVDKWMEGGNTTESPIEPYT 421
WP_089610280.1  401 LGKTVEKWNN--NQVVTKLDQM- 420
WP_072903777.1  390 EKK-------------------- 392
WP_041441130.1  386 L---------------------- 386
WP_243440605.1  383 -IVE------------------- 385
WP_008522721.1  379 -IVE------------------- 381
WP_157150839.1  384 -IIEEPKA--------------- 390
WP_147738704.1  395 -IIEKTGV--------------- 401
WP_155812582.1  407 SVYEEEKI--------------- 414
```

```
WP_041441130.1  386 L---------------------- 386
WP_243440605.1  383 -IVE-------------------- 385
WP_008522721.1  379 -IVE-------------------- 381
WP_157150839.1  384 -IIEEPKA---------------- 390
WP_147738704.1  395 -IIEKTGV---------------- 401
WP_155812582.1  407 SVYEERKI---------------- 414
```
