## Supplemental Figure 1 for "Eight Unexpected Selenoprotein Families in ABC transport, in Organometallic Biochemistry in *Clostridium difficile* and other anaerobes, and in Methylmercury Biosynthesis"

MerB-like protein SaoL (including sequences truncated at selenocysteine-encoding UGA codon)

```
WP_087258257.1   26 LTAEENALRNHIINAIVDTHRPY----APTAEEQ-----------AMVETLQSKNALAIAED-GKIYSIYPVSAKETPHRVTLADGRTTYVMCAIDALGCTYTFHQDIVI
WP_025645259.1    9 FTERQNELRLYIIQFIVDHQRPYHLDQDMTQTLKALGMD-QKEYEEIIECLLKKDGMVIDEE-RNVNFIYPVSALQTSHHVTLQDGREFTAMCAIDAIGAAFTFHQDTEI
WP_242868267.1   10 FTERQNELRLYIIGFILDSHRPFRLDDDMGQALESLNMT-REEYGDITACLLEKDGMVIDEE-RNVNFIYPVSALATSHEVTLADGRSFTAMCAIDAIGAAFTFHQDTEV
WP_242994277.1   29 FTARQNELRLYIINFTIDNKRPYNLESDKEVTLQVLQMD-EQEYEEIIQCLIDKDGMVVDEEAKNVNFIYPVSSLETNHRVTLADGREFTAMCAIDAMGAAFTFHQDTEV
WP_249536531.1   12 FTERENALRLAIINETVNNGRAFDLEEDRVAVLGKLQMS-GKEYKEICEKLLEEDGMVLDGT--KVNFIYPVSALPTNHRVTLADGRTFTAMCAIDAIGAAFTFHQDTEV
WP_242872719.1    9 FTERENQLRLYVIQFTIDRQRAFHLERDEEEAVCALQMS-HEEYEKIIERLIERDGMVIDEE-KNVNFIYPISALETSHRVVLEDGRQFFAMCAIDAIGSAFTLHQNTEI
WP_138307466.1   11 FNEKQNEVRIAIMNFIIDNKRPFHMKQDGYEALKGISLSGKDEFEDILDVLHSRDGFSADEE-GNINFIYPVSALPTAHRVTLADGREFSAMCAIDAIGATFTFHEDTQI
WP_089610284.1    9 FNAAENEIRLAIMEFILNEGRPFNITNDGFLALSDIKLSTAGEFTGIMETLASRDGLVADEE-QNVNFIYPVSALITNHKVTLADGRSFCAMCAIDAIGAAFTFHQDTEV
WP_238493469.1   12 FNAPQNEIRLAIMNFIVDNHRPFSLKEDGYTALKGITLSSAEDFSSITEVLCEKDGMVTDEE-GNVNFIYPVSALPTNHHVTLADGREFCAMCAIDAIGSAFTFHQDTEI
WP_040653003.1   20 FNERENALRFFLMNFIIDHKRAFDLSQDALCAMDACHMS-RKEYQDAISVLRERDGIVSNED-QKVCFVYPISALETEHQVLLGDGRSCFAMCAIDALGTAFSFQQDVEI
WP_163337276.1   21 LSQDAKKARIAILNFSIDQKKPFNIIEDKTAEID-----------VVLQELIDRQAMIVSED-GNVDFIFPVSTKPTQHKVSLQDGREFFAMCAIDALGSAFTFDCATKI
WP_238493463.1   20 LSPEGKNARLEIMNFTIDNQRPYNINSDASPQIL-----------EVIEELREKNAAVISSD-GEIQFIYPVSAIPTGHKVMLEDGRKFSAMCAIDAMGAAFTYRQNVTI
WP_242942357.1   16 YSEEENIIRKAIMDFTIENKRAFSLKKDIEKLFTKIRID---NLEKKLTLLKEHNGFVADED-GNVNFVYPVSALGTNHHVTLADGRRFSAMCAIDAMGTTFTFHQDVVV
WP_238493465.1   17 YSNIENKIRKEIMNFIINNKRAFNIEDDLEALKEIQDLD---LLNH-INNLIKENGIVVDEN-KNVIFAYPVSAVATNHHVKLNDGREFYAMCAIDAMGCAFTFHQDVLV
WP_242944338.1   17 YTKEDNILRKAIMNFVIDNKKPFHIEKDLSLLEVDLNIE---SINESLENLKQNNGFVLNEN-NEAIFVYPVSALPTNHLVTLEDGRKFNAMCAIDAMGTAFTFNQNIEV
WP_238493466.1   25 LLPEEQRLRLMLIMNDIINHQKPVSFESLVSMKEMSPELLK----KSWDGLIRKNAMVMNDQ-QEATFVYPVSALPTPHLVTLADKRQFSAMCGIDAIGAAFTFQQDTSI
WP_216415487.1   24 LNQTEKKTRRFLMNYTIDKDKPFNLVRITEDIVRNIGTSE-DKFNNLINSLINKRAIVVDKD-KNVNFIYPVSALPTNHKVTLSDGRSFYAMCAIDAIGTAFTFKQDINI
WP_238493462.1   24 LNQYERKVRRYLMNYTIDNGKAFNLGETTDEMSKDIGFEK-EELKVLFDNLDNKSSIVVDEN-KNVNFIYPVSAIPTNHKVTLADGRSFNAMCAIDAMGTAFTFNQDIEI
WP_246565988.1   23 LIKEEKILRRHLMNCVINKGQSLDINDVY-EVTSDIKIMDKEVVDKIIGNLLKKNAIVIDEE-RNVNFIYPVSAIPTNHKVILADGRQFSAMCAIDAMGTAFTFKQDVNI
WP_244969178.1   23 LLDKEKKVRRYLMNYVIDNKEAFKINNSF-GIAEDIGMTN-EELNSILKSLIEKNIIVVDED-GKVNFIYPVSAIPTNHKVKLQDGREFTAMCAIDAIGAAFTFKQNVEI
WP_246565854.1   23 LSIGEKRIRRYLMNYVIDNGQAFNINNSS-QVAKNLGMTD-EEVNIILNSLIQKNGIVADEE-GNINFIYPVSAIPTNHRIRLADGREFTAMCAIDAMGAAFTFKQDIDV
WP_244971114.1   29 LSKDEKKIRRYLMNYTIDKGQAFKMFNTS-EIARDIKITY-EEANKILDSLTQKNGLVVDDE-GNVNFIYPVSAIPTNHRVRLQDGREFTAMCAIDAIGSAFTFKQDVDI
WP_240696022.1   23 FSKSQNRIRRYLMNYTINNGQAFNLKNLF-TICNDIKDMTEEEIKDILDVLISKNGIVVDEE-KNVNFIYPVSALATNHKVKLADGREFYAMCAIDAIGTSFTFKQDVDI
WP_242967880.1   20 LTKEENDIRINIMKYMIDNNKAYNLEESYEEIQSRINIKK-YDLNKVIKNLIQKKALVVDEN-DNVNFIYPVSALETNHKVKLEDESVFNAMCAVDAMGSSFTFKQDIEI
WP_238493468.1   36 LSLIENEVRRLIMDYIIYNAIPFNTKETSSIILEKLNISK-EKFKQILDSLVSKNAMVMDED-NNVNFIYPVSAFNTNHKVHLSDGRVFSAMCAVDAMGSSFTFNQNIKI
WP_242842512.1   20 LEADERKVRMHIINSIIDSAAPYNYSVITADVQKKLGMSE-VQVKTAIGRIIEKNAAVADEE-ENINFIYPVSGFPTNHQITLEDGRSFCAMCAVDGMGCAFTFKQNVKV
WP_238493470.1   20 LDSDEKKVRFYIMDRIIENGMPFNFAVIPEAMQKSLNMTE-GMIRETIDSLAAKNAAVSDEN-QNINFIYPVSGFPTNHHIELADGREFSAMCAVDGMGCAFTFRQDIKL
WP_238493467.1   26 LTPDENACRLWLMEYTIDRGIPYNIDN-------GAGAEP-PGSRSSVRSLAEKRAIVVGPD-GNVNFIYPVSALPTQHRVRLSDGRSFCAMCAVDALGAAFTFKQDIRV
WP_240004101.1   36 LSYLEKKVRIYIMDYIIEEKRPYNLKNLDYNFICNIGVAK-DEFINIVLSLKDKKVLVIDEK-ENINFIYPVSAFDTNYKVKLEDGREINAMCAVDSIGTAFTFKQNIKI
WP_251861115.1   36 LSLLEKKVRTYIMDYIISKNKPYNLKNPNNDFIIGTGITK-NQLEDIVLSLMHKKALVMDEK-ENINFIYPVSAINTNYRVELEDGRKINAMCAIDSIGTAFTFKQNIKI
WP_161822871.1   34 LTANERLVRNYIMESIINYAALVNVADLEQVFNTRL-----DDVLAIVDSLVDKRTVVRDEL-GNIVFAYPVSALPTAHQVNLQDGRHLYAMCAVDALGVAFTFKQDIKI
WP_094207760.1   17 LTPEEQKDRIGILNLMVD----------EGGVLPISKCS----KFKILDSLIKKRKLILDEGGKEIQFAYPVSGLPTSHHITLKDGREFYSMCAIDSLGSYFTFWQDLEI
WP_005542126.1   17 LDDRESMFRKKMLDRMIS----------TQDLLHQDDFAG-E-EKEILKSLVDKDVLVLEEG--QIVFAYPVSKTPTNHHVELADGREFFSMCAIDALGSTFVFEQDTKI
WP_136713543.1   23 LSEIEQRVRKSLINYVVNNSIPFNINLDSVDLLNIDNISE-FSIKEIIKGLVAKRILSMDQEG-DINFLYPVSALPTNHIVTLGDGRSFNSMCGIDALGSTMTFNQDVHI
WP_050355226.1   45 LTEIEKKLRRSLMNYTLQYKKPFNIKAPSIEVIKDIGLNI-EKLNEIINRFVEKNIVVVNDKG-DINFIYPVSALPTHHLVTLSDKRKFNAMCAIDSLGSTFTFGQDIEI
WP_238476750.1   19 LDSDMQNARLRLIDKVIETGCTIS--KEEAIKICGD--------EKLYNSLIEKEIVTMSGD--SVAFLYPVSAMETNHRVTLSDGREFCSMCAIDALGSYSLFHQDTEI
WP_242003574.1   13 ISKEIQEARLRLIDFVIETGLPVN--KNEAIKICGN--------ESLYKELLEKEIITANGD--DIAFLYPVSALKTNHIVKLSDGREFFSMCAIDSLGAYSLFHKDVEI
WP_114642964.1   29 LKKKYQDVRLFIMESVIDSKTSLKDIKNELKNKYPD--------IDDILIEMQKKGVFVADE--DVKFIYPVSALPTPHRVKRKDGKEFYSMCGIDSMGTHFTFHQDIEI
WP_051623687.1   28 MNREEERLRRFFMDYIIDNKCSFKLDNINNYLGALDIKES--NVGEIIKNLRDKNGIYVEDN--EVAFVYPVSSKPTNHRVYLEDGRELYAMCAIDSIGCSFTFNKDVYI
WP_238493471.1   29 MSLKEEKLRRFFMDYIIEEKQPFELDSIDKYLDKLQLEKK--EVKKLVKSIEEKEGLWQEDG--FIKFVYPVSSIETNHRVRIEDGREFFAMCAIDAMGSHFTFDKNIKI
WP_026901398.1   10 LENEQKKIRVEIMNMIIEKNSVVSLNEISNNLLNTLELEE-SYIKQTLDYFVKNNIMVVDEEN-YVNFIYPVSAHETMHKVTLQDGRKLNAMCAIDALGVGLTFNQNIAI
WP_216467282.1   10 LDETERHIRLSIMNMIIDEKRSVSVDEVVEKLSAELNLDE-YRLVKTLEHFKNENVLVTDEE--NINFIYPVSALATNHNVTLDDGRSFYAMCAIDAIGTSCTFHQNVTI
WP_242825830.1   10 LDNTQRAIRVSIMDMIIEEKRSITLNEVVNVLSLKLSIDR-GYLQNTLDSFIKENILVAEGD--NINFIYPVSALPTSHKVTLDDGREFFAMCAIDAIGTSCTFHQNVTI
WP_243207816.1    9 LDKIQKEIRIAIMDIIIDKKRPVSLNEVIKELKLRFNLEG-DSINKTLKLFIDENIMVVDEDN-YINYIYPVSAHPTMHKVTLADKREFNAMCAIDALGWRCTFNQDIDI
WP_247011940.1    9 LNQIQKNIRVSIMDMIIEKERPVSLDESLVFLEDKLNLKK-EYIKETLKLFIDKNIMVVDDDG-YINFIYPVSAHPTMHRVTLEDKREFNAMCAIDAMGWRCTFNQDIDI
WP_240006030.1   10 LDNTQQKIRISIMDMIIDKGSSVTLQEVTEYISKKLNMEK-EYIESTLQYFIDKNIMVVDGD--SINFIYPVSALPTNHKVTLRDNRSFSAMCAIDAIGTSCTFNQDVRI
WP_238476748.1   10 LTDEQKAIRISIMDCIIDNGGSVNIELAKKWVKEHMESSE-ETIDATFKQFIDTNVMVVIDD--EITFIFPVSGKPTNHRITLEDGRSFCAMCAIDSIGTSYTFKQPVTI

WP_087258257.1  120 DSVCSNSGVPIHIEVKDGKIASASSDEIRILHADLKASDNWASCCCCQMLYFNSQADYDAY-AKAHDLCPCCSFCLDLNEGLTVGRMLFSDDE------------- 211
WP_025645259.1  117 HSACASCNEPVHIRIVDGKVAEYSPKNLHALTFPLGELSNWAGSC------------------------------------------------------------- 161
WP_242868267.1  118 HSVCASCGSPVYVKVRDGKVAEYAPETLHALTFPLGELTNWAGSCUNVMNFFCSGAEFDQY-VTDMELDAASVIKADIYKAVEEAKATFEV--------------- 207
WP_242994277.1  138 HSVCAMCEEPVYVKIVDGKVADYAPKTLHALTFPLGELANWAGSCUNVMNFFCSSAEFDAY-VTEMELDPNDVIKADIDRALIEAKATFEV--------------- 227
WP_249536531.1  119 HSVSAVSGEPVFVKITDGKVAEYSPKELHALTFPLGEMSNWAGSCUNVMNFFCSGAEFDQY-VADMELDPELVIKADIDRAVKEAYEIFHTEASADDLKRLLAKAI 223
WP_242872719.1  117 FSECAGCGKPVYVKIKSGEVESYEPKTLQALTFPLGEISNWAGSCUNVMNFFCSSAEFDQY-VTDMELDPDKVIKADIKRALKEAKATFEV--------------- 206
WP_138307466.1  120 HSSCCVCGNDVYVEMKNGECAKHRPEGLYALSFCLEEITNWAGSC------------------------------------------------------------- 164
WP_089610284.1  118 RSACSVCGEEVYVKMQGEKVVEHYPEKLHALTFKLEEIANWASSC------------------------------------------------------------- 162
WP_238493469.1  121 HSKCSSCGADVFVRVKDGKIVEHSPADLHALTFTLGEIANWAGSCUNAMNFFCKKNEMEQY-IADMELDLSTVIQADMDIALKEAIATFNV--------------- 210
WP_040653003.1  128 RSRCAVSGREIRISVRDASIASASPENFYALTFKLEELTNWSGFC------------------------------------------------------------- 172
WP_163337276.1  119 NSVCAHCNEPVAIEVRNGSVAYVSCNELRALHVDLNKSDDWSGSC------------------------------------------------------------- 163
WP_238493463.1  118 ESACAHCGESVRVTVKDGGLDLISPSTVHALYFDLNSDNEWAANCUNIINFFCTKEHYSDW-TNARKLDENQIFCLPAKEAIFVSEMIFKLEDR------------ 210
WP_242942357.1  122 ESVCSNTGKSVRVEIRDGKLYDFNPPTLHALHVDLNRHTDWASSCUNIMNFFCTERDFDEW-VSKMQVNKDEIYKLSVEEAIEVSYELFDFR-------------- 212
WP_238493465.1  122 KSKCSNTSEDVYVKITNGEISEYYPKTLHALHVDLNKHTDWGTTCUNIMNFFCTEKDFDIW-TSNMNINIDNIFKLHVTDAIEVSRELFDFR-------------- 212
WP_242944338.1  123 KSKCSNTNTEISVEIVDGEIKNYSPSDLHVLHVDLNKYTDWGKNCUNIMNFFCTEKDFDAW-VDKMKISDIDIFKLHVSDAVYVSKELFEFR-------------- 213
WP_238493466.1  129 SSQCSQCGEPISLEVNDKQLINVHPANAYVLHVDLNQIDDWAASCUNIMNFFCTKNHYNEW-IHTMKLDQKTIFGLSLSDGLLVAQMLFSIDNT------------ 221
WP_216415487.1  132 DSKCSQCGESIYIQIKDGRLLSYIPEDACILHVDLNKHKNWSGDC------------------------------------------------------------- 176
WP_238493462.1  132 ESICSNCGDSIYVEIKDGQLMEYTPEDMHILHVDLNKNKNWSGECUNIMNFFCRKNEYDEW-VAKMNLSEDDIFCLNAIEALQVAKMLFTASDL------------ 224
WP_246565988.1  131 ESKCSNCGADIKVSIRNGKLDSYEPTDLHILHVDLNKNLNWSGNCUNIMNFFCTKKEYDTW-VAKMQLNEFDIFCLNVEEALRVSEMLFSVTDI------------ 223
WP_244969178.1  130 VSTCSNCGSAIKVTVKDGKLYDYEPDNLHILHVDLNKNANWSGNCUNIMNFFCREKEYNLW-VDNMGLNRDEIFCLNAEEAIRVSKMLFSVTDI------------ 222
WP_246565854.1  130 ESKCSNCGFDIKVSIRDGRLHSYEPSDLHILHVDLNKNSNWSGSCUNVMNFFCKEKEYNVW-IEEMELDKSEIFCLNANEAIKVSKMLFSVTDI------------ 222
WP_244971114.1  136 ESKCSQCGSDIKVSIKNGELYSYEPSDLHILHVDLNKNLNWSGNCUNIMNFFCTEKEYNTW-VEIMELDKSQIFCLNAEEAIKVSKMLFSVTDI------------ 228
WP_240696022.1  131 DSQCSHCGTNVKISIRNGKIVKYEPLDLHILHVDLNKNSNWSGSCUNVMNFFCKKNEYDLY-VAEMELNEEDIFCLDVNEAFYVAKMIFSVTDI------------ 223
WP_242967880.1  128 QSKCSLCGEEIFVEIKNREIVNLVPYSTHVLHVDLNKNQNWSGNCUNIMNFFCTKNHYDEW-VNNMNLSEDNIFCLSAREALNVAYMIFNVE-------------- 218
WP_238493468.1  144 NSKCSECGEDIFVEIKNGQLANYSPELTHVLHVDLNKNQNWSGNCUNIMNFFCTKKHYDTW-VNKMQIPEEDIFCLNIQEAILVAKMIFNSNFSKDFPYANV---- 244
WP_242842512.1  128 HSKCSECGADIAVEIRDGQIVSLSPEAAHVLHVDLNGNQNWSGSCUNIMNFFCTKEHYDSW-VKSMELPDDEIFCLEAKEAIWVSKMLFSLED------------- 219
WP_238493470.1  128 TSSCSQCGEQVRVEISNGEIVNLLPEDAHVLHVDLNKNQNWSGSCUNIMNFFCKKEHYDQW-IAEMGVPNDQIFCLEAQEAIWVSRMLFGLEGDI----------- 221
WP_238493467.1  127 ESSCIECGETIHVDIETRRIARLCPESAHVLHVDLNKVANWAASCUNIMNFFCMKKHYDQW-TSNRGVNEEDVFCLDVREAMIVAGMLFTMEE------------- 218
WP_240004101.1  144 ESSCIMCNDSIEIIIENGEIKKCIPENLRVLHVDLNKNDSWASSCUNIMYFFCSEEHYRNW-VKESNINEDNIFCLDTQEAVCVAKMLFSVTDLK----------- 237
WP_251861115.1  144 ESNCITCNESIKIIIENGEIKECMPENLRVLHVDLNKNNSWASSCUNIMYFFCSEDHYRNW-IKETKTDEDNIFCLDIYEAMCVAKMLFSVTD------------- 235
WP_161822871.1  138 ESVCRHCNAPVALQIINGEIFDLHPTTAHVMHVDLKKLDNWAGSC------------------------------------------------------------- 182
WP_094207760.1  113 DSVCSTSGEPIKVVVKNGEIVYKSSPDIHCIHVDLNEFEDWSACC------------------------------------------------------------- 157
WP_005542126.1  113 SSICPVSGEEIHVTIKNKEIVEYSPENIHVIHVDLAAVENWAANC------------------------------------------------------------- 157
WP_136713543.1  131 ESVCSHCGERIVIDVKNRNLEYLSSEEIHVIHMEASENEDWAVNC------------------------------------------------------------- 175
WP_050355226.1  153 LSECSECGEKIKIEIKNGKIKRVSSKNIRVLHVDLNKHKNWSGSC------------------------------------------------------------- 197
WP_238476750.1  117 NSICSQTGEKIYVRIKDRQIVEHSPKDIHVIHVDLNKNKNWASTCUNIMNFFGSKNDMNEWLKNQEVDNNTTYLLDLQTAFKVATAIFQI---------------- 206
WP_242003574.1  111 NSICSNTGNKIYIKIKDRKIFEHCPNDIHVIHVDLNKNKNWASTCUNIMNFFGSKNDMNEWIKNQNINNNTIFLLDLNTAFEVATAIFKL---------------- 200
WP_114642964.1  129 NSKCCETGESVNLKLSDGKLVSYSPEDLHILHVDLNNHTNWAGDC------------------------------------------------------------- 173
WP_051623687.1  134 KSSCSHCGEKITISVKAGEIVDHSPEDIHILHVNTNKFKDWGTSC------------------------------------------------------------- 178
WP_238493471.1  135 NSKCSECGQEIFVEINNGEVVECQPSSLNILHVNTEKFTEMAKTCUNIMNFFCMENELDTW-VNKMNISKELYFKLDILEALDAAKMIFRVDK------------- 226
WP_026901398.1  118 DSVCNITSKEINIKLKDNKLHSINNDNLRILHINLDKHENWAASC------------------------------------------------------------- 162
WP_216467282.1  117 NSMCCVRGTEINIRVKDEQIEYVNNQDLRVLHVNLDKHSEWAASC------------------------------------------------------------- 161
WP_242825830.1  117 DSMCCVTGQEIKIRVEDEQIKYVSNPNLRVLHINLDKHSEWAASCUNIMNFFWTKDEYESWININEMNDDEDIYCLDILGALDVAYKIFKL--------------- 207
WP_243207816.1  117 SSRCYVTGKEINISIKDGKIVSLNNPNLRVLHLNLAKHANWAASCUNIMNFFWTKNEFDKWIKENGLENDEDIYCLDIDEAMEASYDIFYVG-------------- 208
WP_247011940.1  117 KSRCYVTGNEINISIKDGKVTSLNNPDLRVLHLNLDKHENWAANCUNIMNFFWTKNEFDEWVKENGFENDEDIYCLDINEAMDASYDIFYVG-------------- 208
WP_240006030.1  117 NSICSVTGREIEIVIKNEKIEYVNNPSLRVLHINLDKYSNWAANCUNIMNFFWTKSDFDAWTIEAGLSDDEDIYCLDINEAIVESYKIFKLKQKIIL--------- 213
WP_238476748.1  117 NSMCAATGKEIELKVDNGKVLYTNNPGMYIIHVDLLKNTDWANACUNIMNFFWTKNDLEAWLKAAGKENDPDVYAYNLDEAIEESYYTFEV--------------- 207
```
