## Supplemental Figure 6 for "Eight Unexpected Selenoprotein Families in ABC transport, in Organometallic Biochemistry in *Clostridium difficile* and other anaerobes, and in Methylmercury Biosynthesis"

Cys-Cys-COOH protein SaoC

```
WP_252346668.1    1 -------MKKTVSFFKSLKGKIVLIVCILLAGIIIRQSLENLYEEKQVQN--GTVTTYEDGSRVGRSLSYAETVPEDNKILQAFYEQFPTATMLVACEEDLTNDGKKDLV
WP_252346670.1    1 -------MKAVKSFMKGLGGKALIVALIVFAGFGVRQKLERTYEDNQARTEDGEVNVYEDGSRVGRDLDYAENVPEDNKILQEFKKLYPQAKVLVACEEDLTNDGCSDLV
WP_252346671.1    1 -------MEQREKFYHKLWFKAVVVVVVVLIGFSVRTYLEKVYKDSQREE--GTVQFFDDGTRVGRELDYAPKVEDDNKILQAFKAIYPDATVLVACEEDLTDDGLDDLV
WP_044937124.1    1 --MNDKNVKEKTQILKSTKVKIALVMLALVAGFIIRGKLEETRKTQLEEE--------AEEDVMSKRADYAEPVGKDNKILQAFLAYMPGADVQIACEEDVTDDGCKDLV
WP_009249861.1    1 -------MDILKKFWRSVKGKCLIAVLVLLAAFEVHNALEGKEGEKQITQ--------TEARKTKREVEYAKPVDDSNQILGKFKEQFPEASVVLACEEDVTDDGCKDLI
WP_227186491.1    1 -------MDMLKKIAGSTAGRCIIVAVILIAAFIIHNELEGQRDRHQAAQ--------EKQHKPQRDVEYASQVESGNKILNAFEDQFPEAEVILACEEDVTDDGNKDLV
WP_005334080.1    1 -------MSIIKKLEDSIWAKVILAVIVVVIAFAARSVLENKHEESQIDK--------QTAGKTIRETSYAEAVPEDDSILQVFKNTYPTAEVLLACREDVTDDGLDDLV
WP_105308941.1    1 -------MSIIKKLEDSIWTKVILAVVVVVIAFAARSILENKHEESKIDK--------QTAGKTIRETSYAETVPEDDPILNVFKNAYPTAEVLLACREDVTDDGLDDLV
WP_089610267.1    1 ----MNNKFLKN---------ILSIIIVVAVAFAAKQYLEAREERKYQDQ---------TAWMGGEGKDFAVMVPEDNRALQYFKEEFPAREVILACAEDITDDQLQDLL
WP_022132693.1    1 ----MEEKTKKSNKKVSLLIKAVIVLVVVVIAFFLSQYVAEKNEKSTGSR---------RERI------YAPMVEDDNAALVAFQQENPQRKVVLACEEDVTNDTLKDLL
WP_120388140.1    1 MRENGREKTAASPKKT--MIGILAVVLICFTAFGASRFLKERDETINAAR---------RELT------FAPMVADDNEALIAFREQFPERNVVLACKEDVTNDSLPDLL
WP_119205923.1    1 ----MTKKPTRLAPYKRIALYTVLLLSVCLLAFSASRYLQKQAEEKNSGR---------RRLA------FAPAVEKDNEALLAFKKEFPDREIYLACQEDVTDDGISDLL
WP_207746804.1    1 ---MMSKKTKKLTPYKRISLYTILLLSVCFLAFSASKYLRKQTEETNTGR---------RRLA------FAPAVEEDNQALLIFKKEFPDREIYLACEEDVTDDGIPDLL
WP_087258577.1    1 ----MTNNSKAKRSFP-LWARLLLILAAVVAVFFLRQYIYSLEEEKQNSS---------SRLQ------YAESVPEDHEALLYFQQEVPGREIFLACEEDLTDDGRKDLV
WP_009214287.1    1 ---------MMRKRLIRFFALLFFLLLLITALSALRSFLSERERHSAEKAG------------------FAPMVSERDPLLLEFQKRHPERSIILACGEDITGDGRRDLV
WP_093371958.1    1 -----------MKTVKTWGASILIIALVLMAGWNYSQRADGSMEYL-----------------------------ATTPAIDHWRIYYAENELILWDQEDLTDNGKLDTV
WP_093314495.1    1 -----------MKIVKTWFAGFLIIVLVVMAGWNYSQRKEVSIEYLDL---------------------------ATTPAIDHWRKHYEENELILWDQEDLTDNGKLDTV
WP_025436347.1    1 -------MIRK-----ISYVMAMIILIAGLSGCSSNPKSENKKADLGVPA--------------------------DNELLAYFKAQYPDREPVQCAYEDITSDGSKDLI
WP_075727681.1    1 -------MRKK------AGIAIITIIIFGLGVFAFSQDKIKRENNLGVKE--------------------------DNKLLQYFISLYPDKEVLKCGYGDVDGDDRKDLV
WP_212381469.1    1 -------MNKK------FKNLLIIFVIFALGLYSFSLKKSNTIEALGVEP--------------------------DNELLIYFKENFPNNEVLKCGYEDLNDDGRKDLL
WP_154442893.1    1 -------MKKN------IKIILVVFILFSLGFYIYYLNNGNTLQALGVEP--------------------------DNELLLYFKEIHPDNKVILCGYEDLNNDGRKDLL
WP_216515823.1    1 -------MKKN------TKMILIGLVIFALGFYAFFLEKSSSVESLGVEP--------------------------DNELLLYFIETHPDNKVILCGYEDLNDDGRKDLL
WP_094900355.1    1 -------MKKN------IKSILIFLVIFGLGLYSFSIKKNITVNYLGVES--------------------------DNELLLYFQETHPDNDAILCGYEDLNDDGRKDLL
WP_136714224.1    1 -------MNKNV-----VKKILIFTSIFAIGLYGYYMKTRPSLATLGVEP--------------------------DNELLLYFMETHPDNKVILCGYEDLNDDGRKDLL
WP_216542254.1    1 -------MKKK------IKVLLIALIIFSLGLFIYIEDNRQKLIDFGVDY--------------------------DNELLLHFKENFPDNEVIKCGYEDLNGDGIKDLL
WP_099188975.1    1 -------MTKK----ITLNIVFMVLILIIILGGCSEVKEND----IGIEK--------------------------DNPLLNYFEEWNPDNMVIKCAYEDVDNNNTKDLV
WP_216276261.1    1 -------MRKKE---IILYVLVLIIITFMFSACVSAPSKDKELKNIGVTN--------------------------DNALLQYFTKLHPDNEVILCGQEDVNNDNTKDLV
WP_038262032.1    1 -------MTKRK---SILYMILIFMCVTLMG-CSQEERKN---VDMGVER--------------------------DNELFVHFQKKYPENAVIKCGYEDVTNDGAKDLV
WP_099840253.1    1 -------MKN-------IKIFFVFAIVFLLFGCSSNINELKESKNMGVKS--------------------------DNERLLYFQHKYKDKEVLKCAEKDLNNDNKLDLI
WP_163221228.1    1 -------MKN-------IKIFFVFAIVFLLFGCSRNVNEPKQSKNIGVKS--------------------------DNERLLYFQRKYKDKEVLKCGEKDLNNDNKLDLI
WP_076042448.1    1 -------MRN-------IKIFFVFAIVFLLFGCSRNISEPKQSKDIGVKN--------------------------RNERLLYFQHEYKDKEVLKCAEKDLNNDNKLDLI
WP_251861117.1    1 -------MRN-------IKIFFVIVIGFFLSGCFSNVNGYNEKKTIGVKN--------------------------NNEKLLFFKEKYNKKQVLKCGEKDLNNDNKLDLI
WP_050355225.1    1 -------MK--------IKKIFILIISLFLIGCS-NISANEKKKDVGVKE--------------------------NNKALNYFRKHFPEKEIIKCSEKDINGDKINDLV
WP_027634554.1    1 -------MKKFK----GIYFFIFIICL---SFLGCGDK-EEVTE-KVAN-------------------------IKEKELLEFFNKNVEGSKIIVAKEDDINGDELKDLV
WP_090043126.1    1 -----MRFKMRQ----LAVAFILITSI---CVLSACDKSNNKGD-KKIQ-------------------------VSNSKLLEFFENEIPDAEILLAKEEDLNGDGLKDIV
WP_046822596.1    1 -------MKRKK----LLIILSIIVVL---IGISVYSKEKNKQE-SNLP-------------------------DVNEEYLSYFKENVKYEKILTYAQHDINNDKVDDLL
WP_195947720.1    1 -------MKRKK----FLIILSIIVLL---IGIAVYSKEKKKQE-SSLP-------------------------NVNEEYLNYFKENAKYEKILTYTQHDINNDKVDDLL
WP_142729833.1    1 -------MKRKK----ILIVLSIIVLL---IGIAVYSKDKKKQE-NSLP-------------------------NVNEEYLNYFKENAKYEKILTYTQHDINNDKVDDLL
WP_148549829.1    1 -------MKRKK----ILIILSIIVLL---IGIAVYSKEKKKQE-SSLP-------------------------NVNEDYLNYFKENAKYEKILTYTQQDINNDKVDDLL
WP_101509180.1    1 -------MKRKK----LLIIVGVVIVL---VGIGVYSAEKKKEI-KGLP-------------------------KVNEKYLDYFKKNAKYEKIITYTEEDINNDNEKDLL
WP_055336502.1    1 -------MKRKK----LLIIVGVVILL---VGIGAYSAEKKKQI-KDLP-------------------------KVNEKYLDYFRKNAKYEKIITYTEEDINNDNEKDLL
WP_187527164.1    1 -------MKRKK----ILILLGVIVVF---IGIGIYSAEKNKQS-KDLP-------------------------QVNESYLNYFKKNAKYENIITYTQKDINNDNVEDLL
WP_026901405.1    1 -------MKHKK----LFIIIGVVIFI---VGISIYSKDKNTHQ-NNLP-------------------------KVDEDCLKYFQENCEYEEIITYAQHDINNDKEDDLV
WP_227829268.1    1 --------MLKK----VLIISCVVIAL---IVLSRVVDDNLKQN-DSIP-------------------------NVNQETLKYFKKNYK-EDIITCAEEDLNNDGKKDLV
WP_227855571.1    1 --------MLKK----VVIISCVIIAL---IVLSRVVDDNLKQN-DSIP-------------------------NVNQETLTYFKKNHK-EDIITCAEEDLNNDGKKDLV
WP_169467649.1    1 --------MLKK----IVIISCIVVAL---IILSKIVDDNLEED-ASIP-------------------------NVNKETLEYFRKNYK-EDIITCAEEDLNNDGKKDLV
WP_054270966.1    1 --------MLKK----IVIISCIVVAL---IVLSKIVDDNLEED-ASIP-------------------------NVNKETLEYFRKNHK-ENIITCAEEDLNNDGKKDLV
WP_227449898.1    1 --------MLKK----IVIISCIVAVL---IVLSKMVDDNLEED-ASIP-------------------------NVNKETLEYFRKNYK-EDIITYAEEDLNNDGKKDLV
WP_070113298.1    1 --------MLKK----IVIISCIVVVL---IILSKIVDDNIKED-ASIP-------------------------NVNKETLEYFRKNYK-EDIITCAEEDLNNDGKKDLV
WP_107595028.1    1 --------MLKK----IVIISCIVVAL---IVLSKILDDNLKED-ASIP-------------------------NVSKETLEYFRKNYK-EDIITCAEEDLNDDGKKDLV
WP_216467277.1    1 --------MRKK----IIVAGIVIVCL---ILGSKILEENLKED-DGLP-------------------------NVNKQTLKYFKSNCEYKDIITYAEEDVNGDNKKDLV
WP_024620396.1    1 --------MRKK----IIITVVVLVCL---ILGSKLVDENLKED-DGLP-------------------------DVSKQTLKYFKKNCKYKEIITYAEEDVNGDDKKDLA
WP_201416748.1    1 -------MRNKK----IIIGIAIAILAGFALVLSNNYEEKQKQE-SEIDIH----------------GGAVAEKKADKELMDIFKKNHD-EEIITYAQQDMNNDGKDDLL
WP_005542132.1    1 --------MKKT----AIWATAVIILL--EIGFIIYSKDKNIQAGRGIG------------------------VDVNHPFLKKYQKEFENQTIIRAAQEDVNNDGKKDLV
WP_051540177.1    1 -------MIFMK----KFLGIFIIIFTIFLLGCEGQAEKENE---VLSKY-------------------------NENKLLKVFKNEFPERKVILCEAADLTNDNIEDLI
WP_072832006.1    1 -------MKRLS----KISIIFIVIVITMLSGCGKTEKAGNEQ--AAHNY-------------------------EDNKLYIDFKEKFPNREAIICEHADVTNDGLEDLI
WP_084672179.1    1 ---MVKNNLFKK----KCIIAMLVFVCIFSMGCDSKNKVDKVHNEALKVA-------------------------EGHKLYKSFKEMFPSCEPITCKAGDVTNDGKEDLI
WP_114642957.1    1 ---MKKKIKFTK----SNAAFLFIFFIGFIFYMYSSKEVSQEN--YENID-------------------------KKNPLLKIFQSEYSN-KVILMKEGDITNDGIEDLI
WP_073613906.1    1 -------------MIQVHSTILLLLICLFLWGCGGDDRSGTEVGDQRNFS---------------------EEAAGKLPLLSHFLKLHPDRVILKYAEADLDDDGRKDII
WP_049766440.1    1 ----------MKWWKGCGTCMCFVLACLVAAACG--ERPKGEKGAEKAQT---------------------PVSG---VLLEHFKRWNPQREVIKWAEADLDNDGRRDCI
WP_161822866.1    1 --------MLSRT--QLLVILTSLLIFLGCGGCG---NQPAENDAVRATE---------------------ALMRSD-ALLQHFKSVHPDKEVIVFARADVNNTGNDDRV
WP_190239397.1    1 -------MMPRRK--LLQRFFACVVLVLLAGGC---AAGNGKMPPPAAGD---------------------EAFRGG-PMLEYFKAAHPGKEVIILDKADLNSDNTEDMV
WP_214081968.1    1 --------MFKRLPTLTCSLLMCLLVCIAACGCGKVSIDKGEQSDISRES---------------------AAGEPS-PCLGFFISANPGKEVIKCATADLDGDNTEDLV
WP_153189197.1    1 --------MSRRW--LICVLLMCLLAGAAAAGCGKMSVNKGEQSDTSRVS---------------------ATGEPS-PCLSFFALTHEGKEVIKCASADLNGDNIEDLV
WP_012031742.1    1 ---------MKRT---VCYGFCIFLLLIIAPGC---RPGGETKTGTSAVN---------------------GLDKDN-QLLQFFVASHPENVVVKCAAADLNADSLEDLV
WP_028893771.1    1 ---------------MKTRLFPLLAILLLLLACQ----DQPRNSLAEQNK---------------------SALEAS-PLHQYFVRSYPDKPALVWAFHDVNNDGRDDLI
WP_107740074.1    1 -----------MKCLFFIKMLAAAALLFSASGCT--ELATPPLGDQKDFC---------------------PDVLNHNTLLRNFAESYPGNPILKYAQADLNDNGQEDLI
WP_073089019.1    1 -----------MKGKNAIKVALFLIALILAGSLLSKYHASNEGRQVG----------------------------DWPEALREWQEANPGKEVVVWAEGDLDGDGAEDLV
WP_252346672.1    1 -------MKAEKNKSIILRKVLIIIIALLLGYFGYRNNIKYKKVDFGVEK--------------------------DHYLLQAFYDEFPGKEVLKCKLGDVSGNGKDDLI
WP_252346673.1    1 -------MKAEKNKSIILRKVLIIIIALLLGYFGYRNNVKYKKVDFGVEK--------------------------DHYLLQAFYDEYPGKEVVKCKLGDVTGNGTEDLI
WP_252346674.1    1 -------MKAEKNKSIILRKVLIIILALLLGYFGYRNNIKYKKVDFGVEK--------------------------DHYLLQAFYDEFPGKEVVKCKLADVSGNGKNDLI
WP_252346675.1    1 -------MESKKNKSIILIKVSIIILALLLGYFGYRNNIKYKKVDFGVEK--------------------------DHYLLQAFYDKFPGKEVVKCKLGDVSGNGKDDLI
WP_163340325.1    1 -------------MKGSLLISQTILLFIAILSCGCDNSPKKVVEIQQELN-------------------------LSSDLYLKFYKEHSSYQIIVAEEGDINADGLCDLV
WP_051677063.1    1 -------------MR-NAYILFAELLVLSVLCCGCENIQKTPKDVAEGLT-------------------------QCTDIYDVFFQKNPKCTPILATEGDVNDDGDADLV
WP_008520778.1    1 ----------MRRPRIWLQAARLSVLVGLFLN-----GLAYRVWAA----------------------------PVSS-LIERFQECVPDREILLTLAGDCNADGIEDLV
WP_015556509.1    1 ----------MTRLVPWGAALLCALFLLAFPGPGAAPGGAEDVFLP----------------------------VTSHPLMAVFRKSLPDREVLLTTPGDCNADGIEDLV
WP_028329263.1    1 -----------------MLKVLYAVCVFILISCS-NNKEDISNVDG----------------------------IKNNELYEAYIEKYSDKELLIFLEGDFSGDNIDDLI
WP_096737795.1    1 -----------------MLRLFFVLCIFIFVSCS-DNTQDIKSAEG----------------------------INNNELYKAYMKEYSEKELLTFLEGDFSGDNKDDLI
WP_013114956.1    1 -----------------MLRLFFVLCVFIFVSCG-NNTQDIKSAEG----------------------------INNDELYKSYMKEYSEKELLTFLEGDFSGDDKDDLI
WP_219708866.1    1 -----------------MLRMLFALCFFVFISCN-DNTEYIKNAEG----------------------------INNKELYKEYIKKYSDKELLTFLEGDLNGDNTDDLI
WP_157150838.1    1 -----------------MLKIVWALCFFIFISCN-DNTEYIKSAEG----------------------------INNTELYKEYIKKYSDKELLTFLEGDFNGDNIDDLI
WP_147736609.1    1 -----------------MKNLLFIILGIFIVCCKYNNKNNKENLDG----------------------------IKNIELYKEYINKYNDKEILTFLEGDFSGDNISDLI

WP_252346668.1  102 VIYN----TPEE---DEASKNTALVNGGHIRLTVMIDLGNGE-TYEHTEPIPAPVENQKIQFQNIDKADEMEFVLQGQKGAKVGYGIFRVIEG-KAVNLFGEGMEECU- 200
WP_252346670.1  104 VIFN----QPEEEADHEHDASTQLVDGGHIRLVVMVDSGDGE-NYVGSDPIPAPVENQRIKFQDIDQKDEIEFILQGQKGSNVGYGIFRVVDG-SPVNLFGEGMEDCU- 205
WP_252346671.1  102 VIYN----TQEA---DEYSETTTLINGGYVRMVVGIDSGDSE-NYTFSDPVPAPIENQKIKFQNIDKEAEMEFILQGQKGSKVGYGIYRVIDG-ALVNLFAEGLQECU- 200
WP_044937124.1  101 VIYK----KDGL-----------------TRTVVAVDSGDGT-NYEYTEPIPGPIENQKIQFKNIDKEGEIEFIITGEKNGAVGYAIYRMIDG-APTDLFGEGMEDCC- 185
WP_009249861.1   96 VIYT----EKEL-----------------TRTVAVIDSGDGV-NYNFSTPIPGPIENQKIQFKNIDKEGEIEIIVTGEKKGAVGYAIYRMIDG-EMVDLFGEGMEDCC- 180
WP_227186491.1   96 VIYK----EEGL-----------------TRTVTVIDSGDGV-NYTFTAPIPGPIENQKIQFKNIDKEGEIEIIVTGEKNGAVGYAIYRIIDG-EMVDLFGEGMEDCC- 180
WP_005334080.1   96 VICK----MEEG-----------------NRTIVVTDKGDST-NYDFSDPIAAPVENQKIQFKNIDKEGEIEIIITGEKKGAVGYAIYRMIDG-QPVDLFGEGMEDCC- 180
WP_105308941.1   96 VICK----MEEG-----------------NRTIVVTDKGDST-NYDFSDPIPAPVENQKIQFKNIDKEGEIEIIITGEKKGAVGYAIYRMIDG-QPVDLFGEGMEDCC- 180
WP_089610267.1   89 VIY-----RENE----------------HTRLAALIDKGEGH--WYISPEIPAPIENQTIRFKNIDKEAEMEFIITGEKEGAVGYAVYRMIDG-EIKDLFGEGMADCC- 172
WP_022132693.1   92 IIYKEDDPTEGE----------------ITRLVVSIAQQDGS--YTYTEPIPAPIENQGIQFKNIDEEAEMEFIVSGEKDGAAGYAIYRMIDG-QPMDLFGDGMEDCC- 180
WP_120388140.1   94 VIY-----TEGD----------------LTRFVTAIAGQDG---YTYTEPIPAPIENQGIQFKNIDKKDEMEFIISGEKKGAAGYAIYRIIDG-QPKDLFGDGMDDCC- 176
WP_119205923.1   92 VIY-----KEGS----------------LVRFVTAMGQDGGG--YVYTDPIPAPIENQGIQFKNIDHEGEMEFIISGEKNGQAGYAIYRIIDG-QPKDLFGDGMDNCC- 175
WP_207746804.1   93 VIY-----KEGS----------------LVRFVTAIGQDSGG--YVYTDPIPAPIENQGIQFKNIDNEGEMEFIISGEKNGQAGYAIYRIIDG-QPKDLFGDGMDNCC- 176
WP_087258577.1   91 VLYHN--PEEGV----------------INWMVALINRGDGT--YDITEPTRAPIENQAIRFFNMDKEGEMEFVVTGEKDGEVGYAIYRIIDG-ELINLFGEDMEDCC- 177
WP_009214287.1   84 VISG----EEE-------------------RIESIVLYWEGE-ELRETDAVPAPRENQKIKFFDMDRIAPLETLITGEKNGKLGYAVYRIIDG-SFKDLFGEGMKDCC- 166
WP_093371958.1   71 IIFSVG----------------------HRKNNVLVVMDMGD-ELVMTEPIPAPVENQVIEFLDFDNEPPNELYISGSKGPHVGHAIYRIVDG-ELVDLFSMDMSLCC- 154
WP_093314495.1   73 IIFNVG----------------------HRKNNVLVVMDMGD-EFVMTEPIPAPVENQVIKFLDFDNEPPNELYISGSKGPHVGHAIYRIVDD-ELVDLFSMDMSLCC- 156
WP_025436347.1   73 VIF---------------------ENEKDKKQMCAVIAEENE-KHQLTEPIPAPAENQKIQFKDIDEKDELEFIVSGSKNGSVGYAIYR-IIDNRMINLFGEGMEDCCD 158
WP_075727681.1   72 VIF---------------------NNSKRSNGMVVVLDKEG--KYEITGELPAPIDNQKIEFKDIDETKPLEFIVSGSKDGQFGYAIFR-VEGLEVKNLFGEGMRDCC- 155
WP_212381469.1   72 VIY---------------------NESKRKNAMLVVIDKED--GFQLSDHIPAPIENLKIEFKDIDKTGPIEIIISGSKDENFGYSIFRLIDDTELKNLFGEGMEDCC- 156
WP_154442893.1   72 VIY---------------------NEGPKKNAMVVVIDTGE--GFKLSNHTRAPIDNQTIEFKDIDKTGPIEFIVSGSKDGNFGYSIFRLIDDTEIRDLFGEGMEDCC- 156
WP_216515823.1   72 VIY---------------------NEAPRKNAMVVVIDTGN--GFKLSNHTSAPIDNQKIEFKNIDKTGPIEFIVSGSKDGNFGYSIFRLIDDIEIRDLFGEGMEECC- 156
WP_094900355.1   72 VIY---------------------AEEYRKNAMVVVIDTGE--GFKLSDHTHAPIQNQKIEFKDIDKTGPIEFIVSGSKDGKYGYAIFRLIDDKEIKDLFGEGMEDCC- 156
WP_136714224.1   73 VIY---------------------EEGRRKNAMLVVLDNEE--GIKLTDHIQAPVENQEIKFKDIDKTGPIEFIVSGSKDGNYGYAIFRLIDDWELKNLFGDGMEDCC- 157
WP_216542254.1   72 IIY---------------------NVSRRKNEMLVVIDYGD--DYKFSESTPAPLDNQKIEFKDIDKTPPMEVIVSGSKDGNFGYAIFRLIDDTIFKDLFGEGMEDCC- 156
WP_099188975.1   69 VVY---------------------SVSKDKNKMKIVIDNNG--KYEFTEEVDAPRENQNIKFKNIDAKDQIEIIVSGSKNGAVGYAIFR-VVDMKIINLFGQDMEDCC- 152
WP_216276261.1   75 IIY---------------------RVSKTKNAMKIVIAKEH--NYKCSNEVPAPMENQIIKFKNIDDKDQIEVIVSGSKNGQVGYAIFR-LQNMKIVNLFGNDMEDCC- 158
WP_038262032.1   71 VIY---------------------NIEKGKNGMKVVVGGD---EYSISNEVPAPAEDQIIKFKNIDDKDEIEFIVSGSKHGNVGYAIFR-FQQMEIINLFGQDMEDCC- 153
WP_099840253.1   71 VIY---------------------KENNDKNSMVVVLSDKE--KYKITNEVSAPIENQKIEFKDIDKKPPIEFVVSGSKNGSFGYAIYR-IEKDKIINLFGEDMKDCC- 154
WP_163221228.1   71 VIY---------------------KEDNDKNSMVVVLSDKE--KYKITNEVSAPIENQKIEFKDIDKKPPIEFVVSGSKNGSFGYAIYR-IEKGKIINLFGEDMKDCC- 154
WP_076042448.1   71 VIY---------------------KEDNDKNSMVVVLSDKE--KYKITNEVSAPIENQKIEFKDIDKKPPIEFVVSGSKNGSFGYAIYR-IEKSKIINLFGEDMKDCC- 154
WP_251861117.1   71 VIY---------------------KKNSDKNSMVVVLSDKE--RYKITNEVSAPIENQKIEFKDIDKKAPLEFIISGSKNGNLGYAIYR-IEKGKIVDLFGEDMKDCC- 154
WP_050355225.1   69 VIF---------------------KDGKKENSMVVVFSENK--SYEVTNKVHAPISNQNIQFKDIDNKIPIEIIVSGSKNNKIGYAVFR-LEDGKIIDIFGDGMDDCC- 152
```

```
WP_038262032.1   71 VIY---------------------NIEKGKNGMKVVVGGD---EYSISNEVPAPAEDQIIKFKNIDDKDEIEFIVSGSKHGNVGYAIFR-FQQMEIINLFGQDMEDCC- 153
WP_099840253.1   71 VIY---------------------KENNDKNSMVVVLSDKE--KYKITNEVSAPIENQKIEFKDIDKKPPIEFVVSGSKNGSFGYAIYR-IEKDKIINLFGEDMKDCC- 154
WP_163221228.1   71 VIY---------------------KEDNDKNSMVVVLSDKE--KYKITNEVSAPIENQKIEFKDIDKKPPIEFVVSGSKNGSFGYAIYR-IEKGKIINLFGEDMKDCC- 154
WP_076042448.1   71 VIY---------------------KEDNDKNSMVVVLSDKE--KYKITNEVSAPIENQKIEFKDIDKKPPIEFVVSGSKNGSFGYAIYR-IEKSKIINLFGEDMKDCC- 154
WP_251861117.1   71 VIY---------------------KKNSDKNSMVVVLSDKE--RYKITNEVSAPIENQKIEFKDIDKKAPLEFIISGSKNGNLGYAIYR-IEKGKIVDLFGEDMKDCC- 154
WP_050355225.1   69 VIF---------------------KDGKKENSMVVVFSENK--SYEVTNKVHAPISNQNIQFKDIDNKIPIEIIVSGSKNNKIGYAVFR-LEDGKIIDIFGDGMDDCC- 152
WP_027634554.1   70 VIY---------------------NKDNESQMVVVINKSNR--EYKLSDVHKAPKENQDIQFKNIDDKDPMEFIVSGSKNGAVGYAIFRLEGD-KVIDLFGDGMEDCCS 154
WP_090043126.1   73 VIY---------------------TRYNNNEMVVAINNEKD--EFKLSNSHKAPIENQSIQFKDIDDKDPIEFIISGSKNGAVGYGIYRLEND-KVIDLFGEDMNDCC- 156
WP_046822596.1   71 VVY---------------------KKGNNHNEMVGIISGDE--NVYITEPIRAPIDDILIEFKNIDDKDEMELILSGSKNGNVGYAIYRLENK-KLIDLFGEGMNACC- 154
WP_195947720.1   71 VVY---------------------KKGKNHNEMVGVISDNE--NVYMTEPIRAPIDDILIEFKDIDDKDEMELILSGSKNGNVGYAIYRLENK-KLIDLFGEGMNACC- 154
WP_142729833.1   71 VVY---------------------KKGKNHNEMVGVISDDE--NVYMTEPIRAPIDDISIEFKDIDDKNEMELILSGSKNGNVGYAIYRLENK-KLVDLFGEGMNACC- 154
WP_148549829.1   71 VVY---------------------KKDKNHNEMVGVISDDE--NVYITEPIRAPIDDILIEFKDIDDKNEMELILSGSKNGNVGYAIYRLENK-KLVDLFGEGMNACC- 154
WP_101509180.1   71 VVY---------------------KKNNRYNEMVGIISDGD--KVYMTKPILAPQEDVVIEFKNIDNKGQMELILSGSKNGNVGYAIYRLEKD-KLIDLFGEGMNSCC- 154
WP_055336502.1   71 VVY---------------------KKNDRYNEMVGIISDGD--KVYMTKPILAPQEDVVIEFKNIDNKGPMELILSGSKNGNVGYAIYRLEKD-KLIDLFGEGMNLCC- 154
WP_187527164.1   71 VVY---------------------KKNDKYNEMVGIISDES--QMYMTKPILAPQEDVTIEFKNIDNKDEMELILSGSKNGNVGYAIYRLEGD-KFTDLFGEGMNSCC- 154
WP_026901405.1   71 VIY---------------------KKDAKHNQMVAVVKNKD--KMYLTKPILAPREDAVITFKDIDKKDEMEVMISGSKNGNLGYAIYRLQDD-KFIDLFGEDMEACC- 154
WP_227829268.1   69 VIY---------------------KKGNNSNEMVVVVSDKD--GHYITKPIPAPIENQTITFKNIDEKPPIEVIVSGSKNGNVGYAIYRVEGK-KLVDLFGEDMDKCC- 152
WP_227855571.1   69 VIY---------------------KKGNNSNEMAVVVSDAE--SHYITKPIPAPIENQTITFKNIDGKPPIEVIVSGSKNGNVGYAIYRVEGK-KLVDLFGEDMDKCC- 152
WP_169467649.1   69 VIY---------------------KKGSNSNEMVVIVSDKG--SHYITKPIPAPIENQTITFKNIDDKPPIEVIVSGSKNGNVGYAIYRVEGN-KFVDLFGEDMDKCC- 152
WP_054270966.1   69 VIY---------------------KKGNNSNEMVVVVSDKN--SHYITKPIPAPIENQTINFKNIDDEPPIEVIVSGSKNGNVGYAIYRVEGN-KFVDLFGEDMDKCC- 152
WP_227449898.1   69 VIY---------------------KKGNNSNEMVVVVSDKN--SHYITKPIPAPIENQTITFKNIDDKLPVEVIVSGSKNGNVGYAIYRVEGN-KFVDLFGEDMDKCC- 152
WP_070113298.1   69 VIY---------------------KKSNNSNEMVVVVSDKN--SHYITKPIPAPIENQTITFKNIDDKAPIEVIVSGSKNGNVGYAIYRVEGK-KFVNLFGEDMDKCC- 152
WP_107595028.1   69 VIY---------------------KKSNSSNEMVVVVSNKN--SHYITKPIPAPIENQTITFKNIDDKQPIEVIVSGSKNGNVGYAIYRVEDK-KFVDLFGEDMDKCC- 152
WP_216467277.1   70 VIY---------------------KNIEGHNRLIVVASKGD--DYYMTDPITAPVEDQAITFKDIDSKGVMEIMISGSKNKKFGYAIYRLEGN-KLEDLFGENMESCC- 153
WP_024620396.1   70 VIY---------------------KNKENHNRLVVVATKGN--EYYVTEPVAAPVDDQVITFKDIDSKGVMEIMVSGSKNKKIGYAIYRLEGN-KLEDLFGENMESCC- 153
WP_201416748.1   82 VIY---------------------YVNKNSNQMTVVASKGD--EYIFTEPIPAPIDDQQISFKNIDDKDEMEVIVSGSKNGNVGYAIYRLEGN-EMIDLFGQDMESCC- 165
WP_005542132.1   73 VVYNP-------------------KGDEKNYSIVLIYKDEN--DYILSKTAVAPRENVEIKFKNIDDKDNIEFIISGSKNGNYGYAIYRLEEDLEIRDLFSEDMDNCC- 159
WP_051540177.1   72 VIF----------------------KEEKETRMVVLVDSGD--GVKYTNIVPAPIEDQSIKLKNIDNKDEMEFIVSGSKRGNIGYAIFR-IENMKIVDLFGEGMDSCC- 154
WP_072832006.1   73 IIY----------------------KEDKNTRLIVATDSSE--GVKYTNEVPAPIENQSIKLKNIDDEKEMEFIVSGSKRGNLGYAIFR-VENMVLTDLFGDGMEDCC- 155
WP_084672179.1   79 MIY----------------------RSSEKVRMRVLID-GP--SVAVTDEVPAPIENQTIELKDIDNKDEIEFIVSGSKRGNIGYAIYR-IVNNNIKDLFGQDMDQCC- 160
WP_114642957.1   76 VIY----------------------DISKSEKGMVVVLGGE--NK-LTNSVKAPIERQSIRFKNIDEKNEMEFVVSGYRGSKLGYGIYR-IENNRIIDLFGEGMDQCC- 157
WP_073613906.1   77 VIYR---------------------ADGEKNQMSVIRNVVA--APVETNPLPAPISDQMIQFRNIDDTPPLEFILQGRKGAKVGYAIFRIEEG-VLVDLFGDGMEDCC- 160
WP_049766440.1   75 VIYR---------------------VAREKNMMRVVLDRGG--NFVDTNEVPAPYSDQVIQFRDIDGKPPMEFIVQGAKGAKMGFAVFRVEEG-WLTDLFGEGMEDCC- 158
WP_161822866.1   76 VIYR---------------------DTRDQNMMVVVFDAAG--KLQCSNKVPAPATNQMIQFKDIDERPPLEFIASGVKGANYGYAIFRLEAG-VLSDLFGEGMEDCC- 159
WP_190239397.1   77 VIFR---------------------ESKETNSMLVVLDLSG--SYQCTNKVPAPVSNQVIQFKDIDQKPPLEFIVQGMKGANVGYAIYRVEGT-KLEDLFGEGMKDCC- 160
WP_214081968.1   81 VIYR---------------------ESREKNKMLVILDKKG--KYQCTNDVPAPVSNQVITFRDIDDKPPLEFIVQGMKGANIGYVIYRVEGV-ELEDLFGEGMKDCC- 164
WP_153189197.1   79 VIYR---------------------ESEEKNSMLVVLGGTG--ECQCTNEVPAPVSNQMITFKDIDNKPPLEFIVQGMKGANVGYAIYRVEGT-SLEDLFGEGMKDCC- 162
WP_012031742.1   74 VIYR---------------------QGADKCGMLVVLSQPA--GYVFTNEVAAPVSGQVITFKDIDDKPPLEFIVQGMKGANAGYAIFRVEGT-RLEDLFGEGMKDCC- 157
WP_028893771.1   70 LIYR---------------------LDRERNAMRVILSTGG--TYTITNDVPAPISNQTIAFKDIDDKPPMEFIVQGMKGTNMGYAVYRIENS-KLVDLFSEGMAGCCG 154
WP_107740074.1   77 VIYQ---------------------VVKGQNEMRVLLNHGT--DYFVSDPVPAPVSDQMIQFKDIDNKPPLEFLVQGRKGPFIGYAIFRVEDG-EIVDLFSEGMADCC- 160
WP_073089019.1   72 IIYRQ-----------------------HKKCFTRVLIRKGN-DYRLLRDMPAPVENQQIQFRDIDNKPPVELIISGTKGSEVGYAIYRIEEE-GLIDLFAENMDNCC- 154
WP_252346672.1   78 IIF---------------------RKTKEANHLVVAIDSGK--EIYFSEESRAPYENQKIELKDIDNSPPNEFVVSGSKRGQYGYGIFR-IEDKHLHNLFSEGMGVCU- 161
WP_252346673.1   78 IIF---------------------RKTKEANHLVVAIDTGK--EIYFSEEGRAPYENQKIELKDIDNSPPNEFIVSGSKRGQYGYGIFR-IEEKHLHNLFSEGMGVCU- 161
WP_252346674.1   78 IIF---------------------RRTKEANHLVVAIDSGK--GIYFSDESRAPYENQKIEIKNVDNKPPNEFVVSGSKRGQYGYGIFR-IEDKHLHNLFSEGMGICU- 161
WP_252346675.1   78 IIF---------------------RKTKELNHLVVAINSGK--EIYFSEESKAPYENQKIEIKDIDNSPPNEFVVSGSKRGQYGYGVFR-IEGKHLHNLFSEGMGICU- 161
WP_163340325.1   73 LIYHN---------------------PRVGNRMCALMRKTDN-TYEISNEFKAPVSNQNVYMRDIDDTPPVEFVVQGMKGAKTGFAIYRFVDG-RIEDVFGEGMDDCC- 157
WP_051677063.1   72 LLYVG---------------------VDCNNWMCVLLRDKDG-GYTASKTHKAPVSNQSVIFRNIDGKPPLEFIVQGMKGAKSGFAIYRMENG-ELMDVFGEGMEDCC- 156
WP_008520778.1   67 VMYR---------------------ENENENRMVAVYSSNG--NFLVSEWTRAPLENYRMQWRDIDERPPVELLVSGQKGIHIGFAVFRFVEG-KWISLFGEGMEDCC- 150
WP_015556509.1   73 VVYR---------------------EDRDKNHMVAVYSDGG--AFRLSAPVRAPLENCRLQWRDIDEVPPVELLVSGQKGIYIGYAVFRFVDG-EWISLFGDGMDGCC- 156
WP_028329263.1   65 IIYR---------------------ESAYSNKLVGIYKDSN-GRILITAPIPAPIENYTIRFYNIDEKVPNEILVSGQKGANIGFAVYRLENG-QLISLFENGMEKCC- 149
WP_096737795.1   65 IIYR---------------------ESKYSNKLIGIYKDYNRGKILMTSPIPAPIESYTIKFYNIDEKVPNEILVSGKKGAHIGFAVYRLENG-EFISLFENGMEDCC- 150
WP_013114956.1   65 IIYR---------------------ESKYSNKLIGIYKDYNSGKILMTSPIPAPIESYTMKFYNIDEKVPNEILVSGKKGAHIGFAVYRLENG-EFISLFENGMEDCC- 150
WP_219708866.1   65 IIYK---------------------ESEYSNKLIGIYKDSN--KILMTTPIPAPIENYTIKFYNIDEKVPNEILVSGKKGANIGFAVYRLENG-ELISLFENGMEKCC- 148
WP_157150838.1   65 IIYK---------------------ESEYSNKLIGIYKDSN--KILMTTPIPAPIENYIIKFYNIDEKVPNEILVSGKKGANIGFAVYRLENG-EFISLFENGMEKCC- 148
WP_147736609.1   66 IIYR---------------------ESNNSNKLIGIYTDEK-GKILITNPIPAPIENYIIKFYNIDEIEPCEFLVSGQKGANIGFAVYRLENG-EFISLFESGMERCC- 150
```
