## Supplemental Figure 3 for "Eight Unexpected Selenoprotein Families in ABC transport, in Organometallic Biochemistry in *Clostridium difficile* and other anaerobes, and in Methylmercury Biosynthesis"

ABC transporter permease subunit SaoP

```
WP_239591061.1     1 ------MEIKISKDKSDRLTKKKTDGS-----MDILYKIILLVIIFLVWHFAAKDIGSSLLLPMPVDVIKGFFFCVTDAETVTNLFITLQRVLKGFMYALLFGL
WP_242825831.1     1 ------MESKISADKRIAIRKKKNSKSL----IENLYRIILLVVIVTVWQIAAIEIGSSLLLPMPLDVFKGFITCVTDPEIVKNLMITLQRVLKGFLCALGIGL
WP_242834147.1     1 ------MEVKVSANTRLKKK--KINSKL----LDILYKILALVIIIGIWQISAEKIGSSLLLPMPMDVFKGFITCITDPEIVKNVIITLQRVLKGFMYALIFGL
WP_006440689.1     1 ------MQTGVSKNK--LKKTFVSKKSM----LEIFYRIALLVGIYLLWHFTAKDLGSDLLLPMPSKVIEGFLYCLTDPETIKNVFITMQRVMKGFGYALVIGL
WP_239591048.1     1 ------MESKASEYRGRLKYSNITKRNI----RERIYRLLVFIGIIGLWHYAAVRIDSPLLLPKPMEIGKALISSATDPKILLNLSITMKRVLKGFLYALLVGL
WP_239591055.1     1 ---MEKVECKTRAYNSIIRNKDLSKQNI----KDFIYKFIMIIILLFIWQISAMHFDNGLLMPYPIDTLKAFIYCITDKETVINILITLQRVLKGFVYAMLFGL
WP_242943825.1     1 ---MEKAENRASIYSGVLKYKNVTKQDA----VENLYKVLVAAIMIGAWQAAAMKIDNQLLLPYPLVTAKAFVHCVTDAETVKNILITLQRVLKGFLYALAFGL
WP_025436070.1     1 ---ME--NNRAIVYNGSFKYAELKKQNM----LDTLYKLIMAGLLFGIWQIAAMKIDSQLLLPYPAVTMKALLECVTDAETVKNILITLGRVMKGFVFAMLFGL
WP_239591056.1     1 ---MEKVESGVSKNS-----MIIDKKNL----KELVYKIALFIAIIIAWQVAANYYDSALLLPTPKATFDRFVENIQNKEVITNISITLSRVLRGFVWALIIGL
WP_242845427.1     1 ---MAEIESKVLSNTSTGILKETSKNRI----KELMYKLIMMVILIIVWYLLAEHYGNELILPTPKRTGKAFIKCLTSSEIMTNLLITLNRVLKGFMYAMIIGV
WP_239591059.1     1 ---MAKMEYKSCTYNSTNKLSKLKNKNT----KYIFYKILVGLALIGIWQLTAMHYDKEIILPYPKNTAIAFFHCITSIEIVTNILITLGRVLKGFLWSLALGL
WP_239591049.1     1 ---MEESQSKASIYNKSGTLVKSSLLNNP---KIFIYKLIFFSLIIFIWYFLAKYYNNDLILPSPYNTGKSLIQSITSKEVITNILITLKRVLTGFLYAFLIGV
WP_242944339.1     1 -------MADVKVTNYTPRGVKFSGSGK----KEIIYKIILLVIIAIAWQSYAVTKNNDLIMPKFTDTIMQFFNCVVDSKVLLNISITLVRVLKGFLWALIIGV
WP_242838408.1     1 -------MEDAKYENVKAKRIKFSGNSK----KEIIYKVTILAFLIIIWEIYATSKNNALIMPKASDTIKQFFLCVTDMEVILNITITLGRVLKGFLWAMVIGV
WP_242942358.1     1 -------MAEAKYQNTSPKKVKLSGNSR----KEIIYKIIVTIILVTLWQMYAVKQNNALIMPKASDVFTQFISCITDQEVIYNISLTMGRVLKGFGWAMIIGI
WP_239591053.1     1 ---MEEIEREIRLS-AESRGMKKEDARRHARRKQILCNIITFATVVLIWQLAATAYKSDLLLPPPWKTAKAFFFAVHDPDTLKNLLLTLQRVTIGFGIALCAGM
WP_239591051.1     1 ---MEPVGDKARFYNSRNTRLMREKVMGAFGQKQLLYRLITIAFVFGVWQLAANHYHSEFMMPSPWKTMVTFTSVVQDPEVVKNLMITLKRVLTGFLYALMIGV
WP_239591052.1     1 ---MEPVENKARIYNSRNSRLMRETFMGSMGYKKLFYRAITIASFLAVWQLAANYYNSEFMMPSPWTTMVTFAAILHDPEVIANLLLTLKRVLTGFCYALLIGI
WP_073089022.1     1 ---MEKGGSEIFIPSVRSKVVKREKAVS------FFYMLLTLVLFILVWQMAALKYNSELLLPTPLKTAVALYDAVRDPWILKNLLITLKRVLLGFLYALAIGV
WP_084218871.1     1 ---MEKGN---------------NVTKKKIDWTKILYPLIAIAAFIGLWQMAAWSYKSNLLLPPPLKTIQALWRVLQDVSVLKNLLLTLRRVLTGFSIACMIGL
WP_239591050.1     1 ---------------MFPSIIGRFDKAGVKVNGTKFHWLGSVAFLFVIWQAASMYYQSNLLLPSPMVTMIALGDAITDQAVLLNMAITLRRVLLGFFYATLIGL
WP_020003907.1     1 MEN-------NGRQVQQNKSRVINLGNAGNSIFMKFYSVSVLIVIMIVWYFLAKKIDNNLLLPDFLLTCKEFFLSWVDPYTVKNLLITLERVFRGFIIACIIGL
WP_243440603.1     1 MAQN-ERSIVNASGINFSTRRGLNLGAIMGRRKGFVCMLAGTLVVCLLWYFLARAIDRSLILPDFLVTMKTFFLGWFDKKVMSNLGITLMRVFMGCVYAVAIGL
WP_040653001.1     1 MEG-------YGAKLRESERRER------GKRR--LISISVLLLLAVLWYFAAKWIDRPMILPDFSETMGEFLRDWTEPRVLLNLGITLCRVLTGTFYAALIGT
WP_239591047.1     1 MGDN----KQVDRMIHATQNKDRRIWKKFGGKNYNIGAIISLIIIALIWFFAAHAVNSPFIFPYLEDVLYQIVYSLTDMYVLENLGITMRRVITGSVYAFVIGF
WP_242647730.1     1 MEEN--HDQQVDRMRAATNAGDRGRWRKFGGRNRNVAALIGLAVICIIWFAAAHIVDKPFLFPYLEDVFKEVGFSLTDLYVLRNLGITMRRVLTGSLYAFIIGF
WP_239591054.1     1 MNSG---EKEVNRMAQAAKMSDRRNWKKLNGKEFNIYTPIAVGVICVVWFFAAHMSENNFLFPTLESVLFKTVDSLRDLYVLRNIGITLRRVLTGTLYAFLVGL
WP_243129913.1     1 -------------MITGVRRQDRRNWKKLSGEGFNYYTPAAVAVIAVIWFFGAHITNNFFIFPTLESVVYSVFDSLRDLYVLRNIGITMRRVLTGVMYACLIGF
WP_249168726.1     1 MNAQ---QKQVNRMVHATKMKDRRAWKNSSGKDFNYHTPIAIIILGIIWMIAARIVNKPFIFPSLESVIEAFFKAITDLYVLRNIGITIRRVMTGVFYAFIIGL
WP_242965469.1     1 -------------------MTDRRNFRNMVGESYYFSFVIALLVLAAMWYFAAHRVDKPFTFPYLESVIYELVTALADLHVIRSFGITMRRVLIGVCYATLFGF

WP_239591061.1    94 PIGFIMGFSKTFERVLSPVVDSVRQVPIMAWVPLTIVWFGIGDGPTIFLIAFSGVFPIILNTIQGVRAISKDYYNAARSMGASPIVIFTNVIVPASLPDILTGS
WP_242825831.1    95 PIGLIMGFSMTSEKVLSPMIDSVRQVPIMAWVPLTIVWFGIGDGPTIFLIAFAGVFPIILNTIQGVRAISKDYYNAARSMGASPVSIFFNVIIPAALPDILTGS
WP_242834147.1    93 PLGLIMGFSSSCEKLFSPLVDSARQVPIMAWVPLTIVWFGIGDGPTIFLIAFAGVFPIILNTIQGVRSISKDYYNAAKSMGASPFTIFTNVMLPAALPDILTGA
WP_006440689.1    93 PIGFIMGFSNTCEKVLSPVIDSVRQVPIMSWVPLTIVWFGIGDGPTIFLIAFSGIFSIVLNTIHGVRTISKDYYNAAKSMGASPFIIFKDVIFPAALPDILTGA
WP_239591048.1    95 PLGYLMGLSELAEKLLSGIIDSVRQVPIMAWVPLTIIWLGIGDGPTLFMIALSGVFPVILNTIQGVRGISKDYYNAAKSMGAGPLSIFINVVIPASIPDVLVGA
WP_239591055.1    98 PIGFLMGFSKLAQKLLGSFIDSIRQVPIMAWVPLTIVWFGIGDGPTIFLIAFSGVFPIILNTIQGVRNISKDYYNAAESMGASKLSIFIHIIVPASIPYILTGA
WP_242943825.1    98 PLGFLMGFSKLADKLMGGFIDSIRQVPIMAWVPLTIVWFGIGDGPTIFLIAFSGIFPVILNTIQGVRGISKDYFNAARSMGAGRLSIFFNVILPGALPDILTSA
WP_025436070.1    96 PLGFLMGFSKPADKLLGGFIDSIRQVPIMAWVPLTIVWFGIGDGPTIFLIAFSGIFPVILNTIQGVKSISKDYYNAARSMGAGRISVFLNIILPASLPDILTAA
WP_239591056.1    93 PLGFLMGLSKLANKLIGGIVDSIRQVPIMAWVPLTIIWFGIGDGPTIFLIAFSGIFPIILNTIQGVRNISPDYYHAARSMGASYFSIFKDVILPASIPDILTGS
WP_242845427.1    98 PLGLFMGAFKMANNLIGGFIDSIRQVPIMAWVPLTIVWFGIGDGPTIFLITFSGIFPVILNTIAGVKNISKDYYNAARSMGAGPWSIFSNIILPASIPDILTGA
WP_239591059.1    98 PLGFLMGSFPLANELISWIVDSVRQVPIMAWVPLTIVWFGIGDGPTIFLIAFSGIFPIILNTIHGVKSISKDYYNAARSMGAGHLSIFKNIVIPASIPDILTGA
WP_239591049.1    99 PIGFLMGYSKIANGVVEGLIHSLRQVPLMAWVPLTIVWFGLGDGPTIFLIAFTGVFPIILNTIDGIQGISKDYYNAAKSMGAGPVSIFKDIIVPATIPNILTGC
WP_242944339.1    94 PLGFMMGLSKVANGLIGGLVDSVRQVPVMAWVPLTIVWFGIGDGPTIFLIAFTGVFPIILNTIQGVKNISKDYYYAAKSMGAGKVSIFKDIIIPATLPDILTGA
WP_242838408.1    94 PLGFVMGISEFANKLIGGLVDSIRQIPVMAWVPLTIVWFGIGDGPTIFLIAFTGVFPVILNTIQGVRGISKDYYNAAKSMGASKLSILKDIIVPATLPDILTGA
WP_242942358.1    94 FLGFIMGLSRFMNSMLAGFMDSVRQIPIMAWVPLTIVWFGIGDGPTIFLIAFSGVFPVILNTISGVRSISKDYYNAARSMGAGPVSIFKDVVVPGTIPYIMTGA
WP_239591053.1   101 SLGFLMGYSKTAMQLIDPVLGTVRQVPIMAWVPLTIVWFGLGDGPTVFLIAMVGVFPILLNTVAGVQSISADYYNAARTMGAGPWSIFKDVMVPAAMPDILTGM
WP_239591051.1   102 PLGFLMGYSKTALQLFDPLIDSLRQIPIMAWVPLTIVWFGLGDGPTIFLIAFAGVFPIVLNTVAGVQNISRDYYNAARSMGAGPWSIFAHIIVPASLPDILIGG
WP_239591052.1   102 PLGFLMGYSKIAMQLFDPLIDSLRQVPIMAWVPLTIVWFGLGDGPTVFLIAFAGVFPIIMNTIAGVQNISKDYYHAARSMGAKPWSVFAHVIVPASLPDILIGS
WP_073089022.1    96 PLGYLMGYSKTVLQIMDPLIGSLRQVPIMAWVPLTIVWFGLGDGPTVFLIAFSGVFPIILNTIAGVQGIDRDYYHAARSMGAKPWHIFSDVIFPGSLPDIMTGM
WP_084218871.1    87 PVGFAMGTSRALFRFFDPLISSIRQVPMMAWVPLTIIWFGLGDGPTLFLIAMVGIFPIILNTISGVRSISPTYYHAARSMGSGPWSIFARVILPASMPDILTGM
WP_239591050.1    90 PLGFLMGYSGTILKIFDPLISSMRQVPIMAWIPLAIVWLGLGDGPTIFLIAMAGIFPLVLNTINGVQNISPDYYNAAKSMGAGKLSIFTSVIVPSSLPDILNGM
WP_020003907.1    98 PLGLIMGYSKPILLAISPIINSIRQVPIMAWIPLAIVWFGLGDGPTIFLITMSAIFPLIINTIDGVRGIDPNYINAARSMGARNIDIMFDIIFPGALPSFLTGC
WP_243440603.1   104 PLGLLMGSSRTALVALSPYVNSIRQVPIMAWVPLSIIWFGLGDGPTIFMIAMSAVFPILINTISGVMNIDPNYKNAARCMGAGTWQVLRDVVVPGALPSFLTGC
WP_040653001.1    90 VLGLSMGYSETALQLISPIVNSIRQVPIMAWVPLSIVWFGLGEGPTVFLIFMSAVFPLILNTVSGVSSIDRNYINAAKSMGAGTAAIYRDIIVPGAMPGFLTGL
WP_239591047.1   101 PLGMIMGYSPRLLQWMSPFINSLRQVPIMAWVPLAIVWFGIGDGPTLFLIAFSGVFTIILNTIAGVQDISKDFYNAARSMGAGTLSIIKDIVIPGSLPGVLTGL
WP_242647730.1   103 PLGMLMGYSPQILKALSPFMNSLRQVPIMAWVPLAIVWFGIGDGPTIFLIAFSGIFTIILNTISGVQDISKDYYHAARSMGASTFGVIKDVVLPGSLPGVLTGL
WP_239591054.1   102 PLGLIMGYSPKMLKALAPFINSLRQIPIMAWVPLSIVWFGLGDGPTIFMIAFSGVFTIILNTIAGVQDISKDFYNAARSMGASTLDIVKDIVVPGALPGVITGI
WP_243129913.1    92 PIGLLMGYSPKLMKAIAPFINSLRQVPIMAWVPLSIVWFGLGDGPTIFMIAFSGVFTLILNTIAGVQDISRDYYHAVRSMGASTWDVIRDVIMPGALPGVITGI
WP_249168726.1   102 PIGMIMGYSQNMLRAFAPFINSLRQVPIMAWVPLSIIWFGLGDGPTIFMIAFTGIFTVILNTIAGVQDISKEYYHAARSMGARTIDIIKDIVLPGSLPGIITGI
WP_242965469.1    86 PLGMLMGYSRLVMQTIAPFINSLRQIPTTSWVPLAIIWFGLGDGPTIFVITFNAIFTIILNTVAGVQSIQSDFYNAARSMGANTLDLVKDVVLPGSLSGLITGI

WP_239591061.1   198 RIAISTGWMSVIUAEFIATSAGLGYSMVQAQTLMHTDVLIGLMIFAALIGFIIDRVLKFINKILCRWRFAD----- 268
WP_242825831.1   199 RLAISTGWMSVIUAEFIATSAGLGYSMVQAQTLMRTELLVALMCMAAIIGFLIDLILKFVNKTLCKWRFAD----- 269
WP_242834147.1   197 RLAISTGWMSVIUAEFIATSAGLGYSMVQAQTMMNTELLIGLMIFAALIGFIIDRILKFINKSLCKWRFDN----- 267
WP_006440689.1   197 RLAISNGWMSVI------------------------------------------------------------- 208
WP_239591048.1   199 RLAVGIGWMSVVUAEFIATSAGLGYSMVEAQARMQTDVLISLMFMSAIVGFGVDKILQTINKRLTKWRQVQ----- 269
WP_239591055.1   202 RIAISSGWMSVIUAEFIATSAGLGYSMVQAQTRMQTEMLVALMFFAALVGFGIDKIIQSINKSLTRWRIAD----- 272
WP_242943825.1   202 RIAISAGWMSVIUAEFIATSAGLGYSMVEAQTRMQTETLVALMFFAAFVGFGIDRMIQYMNKALTKWRYVE----- 272
WP_025436070.1   200 RIAIGAGWMSVI------------------------------------------------------------- 211
WP_239591056.1   197 RIAISSGWMSVIUAEFIATSAGLGYSMVEAQVRMQSDSLISLMFLAAIVGFLIDRTLQLLNKSLTKWRYVK----- 267
WP_242845427.1   202 RIAISGGWMSVIUAEFIATSAGLGFSMVEAQTKMQTDMLVALMILAALIGFFIDRGLLQLNKLLTKWRYVQQ---- 273
WP_239591059.1   202 RLAISTGWMSVIUAEYIAAGAGLGFSMVEAQTRMQTDMLIALMVFAALVGFGIDRILQLLNKRLTKWRYAQ----- 272
WP_239591049.1   203 RIAISNGWMSVIUAEFIATSAGFGYSLVEAQTMMQTDKLVALMIMAALVGFGIDILLQLLNKTLTKWRVN------ 272
WP_242944339.1   198 RIAISSGWMSVIUAEFIATSQGLGFLMVDAQTMMQTDRLIALMVLAALVGFAIDRALNLLNNTLSKWRVVE----- 268
WP_242838408.1   198 RIAISSGWMSVIUAEFIATSQGLGFLMVNSQTMMQTDRLIALMIFAAIVGFVIDRLLNLLNKYLSRWRTNEQ---- 269
WP_242942358.1   198 RVALSSGWMSVIUAEFIATSQGLGFLMVRSQTMMMTDRLIALMIFAALIGFIIDRGLYALNKYLVRWRREE----- 268
WP_239591053.1   205 RLAVSAGWMSVIUAEFIATSAGFGYSMVEAQTRMETPLLIALMFMAALVGYGIDRLLSFVNGMLTRWKCA------ 274
WP_239591051.1   206 RVAVGLGWMSVIUAEFIATSAGFGYSMVEAQTRMQTGKLVALMIIAALIGYGIDRILQFFNRSLTKWRYIQ----- 276
WP_239591052.1   206 RVAIGLGWMSVIUAEFIATSAGFGYSMVEAQTRMETDKLVALMIMAALVGYGIDRVIQLFNSKATKWRFIQ----- 276
WP_073089022.1   200 RIAVGAGWMSVICAEFIATSAGFGYAMVEAQTRMYTDKLIALMIMAGIVGYLIDRLLLFLNIRLIKWRYVA----- 270
WP_084218871.1   191 RLSMGMGWMSVICAEFIATSAGFGFAMVEAQTRMETDSLLALMIMGASVGFAIDKSIRFVESRLTHWRYAK----- 261
WP_239591050.1   194 RLAISAGWMSVIUAEFIATSAGFGYSMVQAQTRMQTDRLIALMIMAGLVGYIIDRLIMLLQRALVKWKYV------ 263
WP_020003907.1   202 RLALGLGWMSVICAEFIATSKGFGFILVEAQQRLETPKLYALMIVSAIIGFSMDRVLVILEKILTSWRFKNGNS-- 275
WP_243440603.1   208 RLALGGAWMSVICAEFIATSKGFGYIMVEAQVRMNTPLLYALMIMSALVGFAMDKSIVLLEHSLTGWRYKDGAPDR 283
WP_040653001.1   194 RLAVGAGWMSVICAEFIATNKGFGYLLIEAQQRLQTSKLYSLMIMSALVGLFIDQLIRLLERKLTGWRFRNGKT-- 267
WP_239591047.1   205 RLAIGLGWMSVIUAEFIATSAGFGYIMVEAQTHLETATVIAYMIIAAVIGFLIDTLLVLFERVLLKWRAQI----- 275
WP_242647730.1   207 RLAVGLGWMSVIUAEFIATSAGFGYCMVEAQAKMQTEKVIAYMVIAGLIGFLIDTLMVLAEKVLLKWRAQ------ 276
WP_239591054.1   206 RLAIGMGWMSVIUAEFIATSAGFGYLMVEAQGRMETATIMAYMLIAGLVGFAIDRIMLLVETVLLRWKAD------ 275
WP_243129913.1   196 RLAISAGWMSVIUAEFIATSAGFGYCMVEAQGRMRTDVMMAYMIIAGLIGFVIDKGMLLIESILLKWKVD------ 265
WP_249168726.1   206 RLAIGLGWMSVIUAEFIATSAGFGYCIVEAQSRMETHTIIAYMIIAGLIGFSIDKVMLLIESMLLRWKAD------ 275
WP_242965469.1   190 RIALGHAWMSVVUAEFIATNSGFGYLIKEAQLEMLTEKVIAYMILAGVFGFLQDRVLLVVESVLLRWKQ------- 258
```
