## Supplemental Figure 2 for "Eight Unexpected Selenoprotein Families in ABC transport, in Organometallic Biochemistry in *Clostridium difficile* and other anaerobes, and in Methylmercury Biosynthesis"

ABC transporter substrate-binding protein SaoB

```
WP_245398079.1    1 -----------MENIKNIIILFIIVLGSIFIIFFNFEDKEEIEGKLRIGVSDDTSGFVINYMVNRDY-FENIKVSDVMDTFTINDCUASTMQWALSSEEIDIAIICKDAA
WP_216467279.1    1 ---------------------------------------------------------------------------------------------------MDMAIICKDAA
WP_027702356.1    1 ----------------------------------------------------------------------------------------------MSSEEMDMAIICKDAA
WP_239591062.1    1 -----------MDKSKNILIGLAILIGSLFILFFNPNS-EKIDGNIRVEVSDDTSGFVINYMKNQNY-FSDVDVENLLEPYSINDCUASTMQWALSSDQIDMAIICKEAA
WP_239591063.1    1 -----------MDKTKNIILGLIIIIGSLGILFFNPKT-EKIDGNIRIGVSDDTSGFIIDYMINKDY-FTDIEIENLIETYSINDCUASTMQWALSSEQIDMAIICKEAA
WP_026901402.1    1 ----------------------------------------------------------------------------------------------MSSEEVDMAIICKEAA
WP_239591072.1    1 ------------MKIKDIVKILSIIFIFAFIVLFNFKT-EKITGNIRLGVSDDTSGLILNYMQDKNY-FENVESENIIDSYSIADCUSASMQWALSSDEMDMAIICKDAA
WP_187296238.1    1 ----------------------------------------------------------------------------------------------MSSNQMDLAIICEDAA
WP_164952401.1    1 ----------------------------------------------------------------------------------------------MSSDVLDLAIVCVNAA
WP_251861114.1    1 ------------MKYFKKVIMTLIILLIFFIILNKLPDESLNS-KIKIGVSDDSSGLIINYMINNDR-LKNVELDKNFEPYFIKDCUGNTSQWALSSDVLDMAIMCPEAA
WP_239591069.1    1 ------------MKYLKKIIITLTILIIFILVLNKLPDESLNS-KIKIGASDDSSGLVINYMINNNR-LKNADLDENFEPYFIKDCUDNTSQWALSSDVLDIAIMCPEAA
WP_050355227.1    1 ------------MRLINKRTVLLLIVMIIIVLIINLPKESSSE-KIKVGVSDDSSGIVLNYMIKNG--FKAIHMQKDFETYSIKDCCSNTSQWALTSDLLDISILCPDAA
WP_239591067.1    1 ---------------MNRIFLIIFIIGIAVAITFFLPSEVLVGGKIRVGVSDDISGFVVDYMIKEE----KLKIGDELEAYFIKDCUAKTSQWALSSDMLDMAIVCKEAA
WP_090043123.1    1 ----------------------------------------------------------------------------------------------MSSNSLQVALLCSDAA
WP_239591070.1    1 ---MRTGKLKQQKLTNNLRKTLTFIILIGALVLGFYLPKEDKKKEITVAAGDDTSGIVINYMLKNKK-FNGKVLK-DIEGLTMNDCUSNNTQWALSSDNIEIAILCSNAA
WP_239591071.1    1 ------------MESVRLLKNTIILAAVAVAVFLFMPKES-LSESVRVGASTESSGLLLDYISNSTE-AAGKVESLELEAYIIGDCUSASAQWALSSDVMDIAFLCPDAA
WP_239591068.1    1 ------------MEHIKKILISTAAVFFVFAVFFLLPKEQRLSANVRIAASNDSSGMLLDYMIRNKD-KSGNIKNLDVESYILGDCUSPSAQWALSSDSVDMAIICPDSA
WP_242838414.1    1 --------------------------------------------------------------------------------------------------------MCSSKA
WP_239591065.1    1 -----MKRKLMKSIRKSFAFMPVLCIAAFIVIFCFHYYENKKSQGIGIGAADDMSGFVIDFIIREES----LEPNKKIKPFFIRDCUAKTSQWALSTEVYSMAILCKDAA
WP_143156824.1   11 LMLKHFHCMQYIMRGRRSKKHLLLPFLLIIFMPFLGGCQSKAHDFLVIGVPDDAGGLIVEYMIANK--LPGLKVEERFDMYAIKDCCSSTSAWALSSSEVNMAVICPDAA
WP_093314491.1    1 MTSQKNESVKNIIKKRIGILIPLLISGMMLWMHFHYEAD-MPDGSLILGVPDDAGGLVVDYLINEK--NFPVKQEDVYGIHTLRDCCSSTAEWALSGNRMHMAIVCPDAA
WP_073613908.1    1 ------------MQRRQLSYAFLFLVLGLISIGLVYGNDADKREIRRIGIPDGAAGILFQYVLKEKH-REPAIQTVHFEPYTLYDCCASAGQYALSSGKLDLAVMCPDAA
WP_214081966.1    1 ------------MS-KVFAVTVVA-ALLLLAACTASPQKEGNNGKVRIGAPVDTGGMIVHYLVNEKG-YQEAEVLGDFELYQVKDCCASTSQWALSTNQYEIAVMCPDAA
WP_134213589.1    1 ------------MK-KVFAVLVAS-VLLFLVAFSAFPQREENNGKVKIGAPDDTGGMIIHYLINEKG-YGDAETQGDFEIYPVKDCCASTSQWALSSDLLDLAVMCPDAA
WP_134219406.1    1 ------------MK-KLFVIAAVA-VLL-LTAF-LYPQRGGDNGKVKIGAPDDTGGMIIHYLVHEKG-YGDAEVQHDFELYPVKDCCSSTSQWALSSNQYDLAVMCPDAA
WP_012031744.1    1 ------------MK-KMLAATAAA-VLLLFTFFTVFPQREESGGKVKIGTPDDTGGMIIHYLINEKG-FQGAEARDDFEIYPVKDCCSSTSQWALSTGQYNLAVMCPDAA
WP_011700728.1    1 ------------MKGGVFAFALLSFVMAAVVGLFSMESEERTPSGIRIGAPDDSAGLVIHCIVGGSG-FEAGKIASDYDPYVMKDCCAGTSEWAFGTNRLDVALMCPDAA
WP_163337278.1    1 -----------------------MIVVTALHLGDEKGLP----ILVNIGAPGDTAGLLLAVAVEESE-NIDLQLRRGIETLLVNDCCAVVSQWAMSSESLDLALMCPDSA
WP_031483264.1    1 ---------------MKGSLLILLGLVVALSVAAAFGSPTESTTDIRIGTPSDTAGLLADLAVEEAD-IEGLTTVSGIKTFQINDCCTPVAQWAMSSQSVDLALMCPDSA
WP_156919681.1    1 ------MEKEGDGPRTVIPMKRFGSRCILLAAAWLLFSCTEKHEGIRIAAPDDFCGVLAYYLVHNK---ELVNAESAEEVFPLVDCCSTTTEWAMSADRIDAAWLCPDAA
WP_146164898.1    1 --------MGYRALTANYARAAVATALCLTVLFCGCNRANQDASSIRIGVPSGAGGLLVHYLVHHVD-ADNLIRSFNYEIFSLEDCCAAISQFALLSLAVDAAILCPTAA
WP_028329265.1    1 ------MKHIIKRNKYIIIFSVIFILCFILFQNYKENINSK-KNKITIACGNDASGMLFDYIINSK---NSNLEFSNFDYIQFLDCCGSQAEFAFISGKVDMAVLCPDAA
WP_014932783.1    1 ------MRHIIKRNKYIIVFSIIFVICFILFQNYKNSLDKK--DTITIACGNDASGMLFDYIINTKK--NSNIEFSNIDYIQFLDCCGSQAEFAFISGRVDMAVLCPDAA
WP_147736607.1    1 ------MRYLKNIIIFTMFTVILIIVGCNKNANSDNSASYIMTNKIIIACGTDASGMLMSYILKNK---NSNIELSNFDYTELMDCCGAQAEFAFMSGKVDMAILCPDAS
WP_208873081.1    1 ------MTGSLNR-RPSPLVLILCAALAATAVFLAAPARRPHGERLMIACGEDLAGQIVAYVARTGSARVSSVGMKQLSFLQLSDCCGTQAEFALAGGDFDMAVLCPDAA
WP_039879528.1    1 ------MTVSLNR-RFIPLFAILGAALAVTLLQEITPRRAPR-EYLKIACGNDLAGHVIAHAAQDYCAKAKAVDMNRLSFLQLSDCCGTQAEFALAGGDFDMAVLCPDAA
WP_009214282.1    1 -----------MRRGRYRIFLLLLLPAVLLAFSLLWPYLRRREEKFRIACGEDLASRLFMELIEEQR-----DVDAVADPTVFGDCCGSYAMMSLVTGELDAALLCEDAA
WP_195838058.1    1 ----------------------------------------------------------------------------------------------MSSDEIDMAFYCNHMA
WP_216560655.1    1 ----------------------------------------------------------------------------------------------MSSDEIDMAFYCNHMA
WP_246565990.1    1 ------------------------------------------------------------------------------------------------------------MA
WP_216542239.1    1 -----------------------------------------------------------------------------------------------------MAFYCNHMA
WP_169300993.1    1 MRLNHKIMLVALVIFIIGFCSYYFLPREKHALNSEENLKIGAGDDITGILLKRIIELNKAEQGGKLLSVEENLDEEDLEFYTFKDCCSNTAQWALTAKEIDMAFYCNHMA
WP_212381464.1    1 ----------------------------------------------------------------------------------------------MSSEEIDMAFYCNHIA
WP_198306502.1    1 ----------------------------------------------------------------------------------------------MSSDEIDMGFYCNHKA
WP_120388142.1    1 ----------------------------------------------------------------------------------------------MRSDEIDLAFYCSHMA
WP_239591073.1    1 -MEIQKKILLAGVGLVVVLSAMQFILPRENEKNAQAQLRIGAGDDISGYLVQEILSQQDSPVQLSEQSMEG---------YEFQDCUSNTAQWALRSEEIDMALSCSHQA
WP_138307482.1    1 ------------------------------------------------------------------------------------------------------------MA
WP_242996699.1    1 ---------------------------------------------------------------------------------------------------MNLAFYCQHVA
WP_242872717.1    1 -----------------------------------------------------------------------------------------------------MAFYCSHVA
WP_016219572.1    1 ----------------------------------------------------------------------------------------------MSSEEINVAFYCSHIA
WP_005334056.1    1 MKIDRVIGIIGGVFILAGIGAALFLPAE-KKWNKDADIRVGSGADISG----VLMEETLQEINKKYKVSTSTESSS------FQDCCSNTAQWALNAKEINIGFYCPHIA
WP_004608618.1    1 MKIQRAIVITLIVFLAAGLGAALFLPKE-KRWNKEAEIKIGAGDDISG----VLMDETVEAL-KDYKVSRTHQSSS------FQDCCSNTAQWAMNAKEINVGFYCTHIA
WP_066572758.1    1 MRINKIAAVVFGIFLTVGVAAYILLPEE-ERRNKEAEIKIGAGDDISG----VLMDETVRNLSKEYEVTQSLESSS------FQDCCSNTAQWALNAKEINVGFYCTHIA
WP_087258574.1    1 MNAMKRGAIFLSAALVLGLLLAFVLPREGKRPADDETLVVGVGDDITG----RLMEQILQRYEADGQEGLTHAGGEDLDSYLFIDCCSNAGQWSLLTQDIDLGFYCSHIS

WP_245398079.1   99 KQYVNATSGFEIVGTVVQNSDVFLLSN-------SNPKK--IGIIQNRNYQSELVKKYY-ENVEISPLLGTALAYGLESNLVDGVVIDCIKSIGLEGTRK-STTELGDYD
WP_216467279.1   12 KEYVKNDANFEIVGPLIKNSDIFLLGP-------KIPKS--IGVTQKRDYQYDLVKKYY-KDAEIVPLIGTALGYGLESNLVDAIVIDAIKSLTIKGDKR-STATEGEYE
WP_027702356.1   17 EEYIKNDTNFEIIAPVVKNSDIFLIGT-------DKPKR--IGVTQKRDYQYNLVSEYY-KDAETVPLIGTALGYGLESNLVDGIVVDIMKSLSLKGEKL-STATNREYE
WP_239591062.1   98 DKFVKYDKDFEIVNALVQNSDVFLLKD-------KNPKK--IGYTQNRNYQLDLIKQYY-KKAKSIPFISGGLAYALESNKVDGVIVDAIKSLSLEGTKL-STSINNSYD
WP_239591063.1   98 KKFVEYDENFEIVDTVVQNSDIFLVKN-------ENIKK--VGVTQNRNYQIDLVKEYY-ESAKEVPFISGGLAYALESDKVDAIVIDSIKALSLNGKKF-STAINKDYD
WP_026901402.1   17 KEFIKADNNFEIVNTLVQNSDIFLTKK-------DDIKK--IGTSQNRNYQNDLVKEYY-KDSKPVSFINSSLPYALESNKVDAIVIDAIKALSIQGKKQ-STSLNKDYD
WP_239591072.1   97 EKYVQTEKNFEIVGTVVKNSDVFLLSD-------ENPNV--MGTVQNRKYQDELVKEYY-PNAKVKHLLANALPYSIESNKVDGGVMDVIKSLPIEGNKK-SCATNKDFD
WP_187296238.1   17 KNFVKNNNEFKIVGSCVKNSDVFILKDN------KIPKK--IGIAQNRKYQKELVKSKFGSSCEVYPMTATALGYALENNTLDAILIDALKSWKLNGKKE-ISNYDKDYI
WP_164952401.1   17 EKFIEKDPRFEIVGVCVENSDIFVTKSG------VTPKK--IGVTQNKKYQEQIVKDRFGNKCETVPMLGRALPYALENNLVDGVILDVTKGWMLNGVKD-KISVNRDYV
WP_251861114.1   97 KNLMKNDTRFEIVCPLVNNSDMFVLKDL------NNVRN--IAMTQNRFYQEKIIKDQYGKGVQIYSMIYSALPYALEKKNVDGIIIDSIKALKLHGYKK-GISVRTNYT
WP_239591069.1   97 KNLIENDDRFEIICPLVSNSDIFVLKDE------NKIKK--IAMTQNRFYQEKIIKDEYGEKVQIYPMIYSSLPYTLEKGRVDGIIIDSIKALKLNGEKK-ALSINKNYT
WP_050355227.1   96 KALVEKDSRFEINSIVMLNSDIFVLRK-------NNPKK--IGITQKRYYQKDLVKERFGNDVEIVPMISTALPYALEKGMVDGIVIDILKGLSIDGEKL-SAHTNKDIP
WP_239591067.1   92 KEYLENVEGFQIVSNSMMNSDIFITKT-------ANPKK--IGMTQDRDYQVDLIKKRFGEDIEIAPMMVSALPYALEKGEVDGIIIDFIKGVHVNGIKE-SSVIDGDYT
WP_090043123.1   17 EEFIKNNQEFEVVGPIVKNSDVLILNK-------NNPKK--IAVSQKRIKDSEIIKKKFKDQVEIVPTLHNILPYVLEKGEVDGVIIDAKNATFLNGYDEGAYEIIGEYI
WP_239591070.1  106 KDFTENNKDFVIVGTVVKNADVIVSRT-------DVPKT--IGVSQNRFNKLNLVNKRYKDNIKPVFLLNNILPYALEKGEVDAIILDSKNAILLQGDKEGTYNINGDYD
WP_239591071.1   97 REFVKTNKEFEIIGDVMQNTSMFVFKDR------GNPGK--IGVSQNRPHQDAIVSENFGANAQMKKLMGTALPYALENGQVDSVLADFTSGFFLKGQKE-SAATKSDHT
WP_239591068.1   98 VEFLKSTDEYVLVGSCIQNSSIIVSKTG------MPARR--VGYAQNRPHIQSMVDKLYPEAVEKKALIMHALPYSLENGMVDTVLLDITTGLSLSGKKN-NAKLENPIV
WP_242838414.1    7 NEFVKSDNRFEIVSPVIQNSNLVVMKNE---------KIKKIGVSEGRSYEKDIAKKMFKETFEIVELPSKSLPYAMEKGDVGGIVIDYLKSLEVIGERK-IIDIGEDYN
WP_239591065.1  102 KTLVENNKDLEIVCPLFQNTDVIVKKNE---------HIKKIAIQQSKFFHEEYLLSIYPESLIIQTN-RTALAYAYESKQVDAVFIDIAKSGGLEGIRE----YPDLGD
WP_143156824.1  119 EDLIAKDERYVIVGPVLANSDLFVLGRE------QNIKT--IGYTQNRWYQKDLIHKKW-KGVRPVPLLSTALPYALEKGVVDGVVIDVSKCQGIKGTLMPVYEEGQDVI
WP_093314491.1  108 QRLIEKDPRYQIVGPVLANSDMLVKHPS------KSVKK--IGITQNRWYQKEIIEKQFGEEVEVAAMLPAALPYAYEMGEVQGIVVDITQGIYLPGEHLPGYKKS-ERI
WP_073613908.1   98 EALVARDDRYRIVGPIMENSDVIFLHPD------ADLQAPTIAISQKRKYQQQMVASRFAGRGKCMPMLHSAIPFAYARGVVQGAVLDIVKTRDLSGVAYPAAPTEPGVV
WP_214081966.1   96 VSLLEKDSRFEIVTPVLINSDIVVARPG------TEPQQ--IGVSQNRSHQEQLATAVFGPGVISAPMLPAAIPYAYEKKAVDAVVVDALRGFTMDGEKIPASTAAGGHV
WP_134213589.1   96 ASLLEKDGRFEIVSPCLVNSDIVVVRPG------VVPQK--IGVAQNRSHQEQIAAGVFGPDVATAPMLPAAIPYAYEKNAVDGVVVDALRGFTMAGEKIPAPAVPDGHI
WP_134219406.1   94 ESLLEKDGRFEIVSPCLMNSDIVVVKPG------LTPQK--IGIAQNRTHQTQIVADIFGQGCATAPMLPAAIPYAYEKNAVDGVVVDALRGFTMTGDKLPATPVEAGHI
WP_012031744.1   96 ASLIEKDGRFEIAGPCLANSDIVVVKPG------LTPRK--IGVARNRSHQAQIAADIFGKGCAAVPMLPSALPCAYEKSAVDGVVVDALRGISMEGEKLSAGG-ATGHV
WP_011700728.1   98 RRLVEKSSMFEIAGPAVLNSDVLVIRPG------AVPKR--IGIAHRRFYQERLVTEWFTSGCSVVPMLPASLPFAFERGVVDAVVVDVLKAFALSGERISVRRSGADVV
WP_163337278.1   83 LALVERDNRFEVVGPILMNAEVLVTKKD------RPILS--IGFAQGRIRHAEHVASTFKDSVAVSAMLPASLPYAYEKNVVDGVVVDFLTALTLNGNKTSLAGAEKHNI
WP_031483264.1   95 GALVAKDARFNIIGPILYNSEVLVSRLD------TKILS--VGVAQGRTRLEPILKEALGEEISVSPMLPTSLPYAYKKKVVDAVVVDFFTALNMPGQITRLAQQKNENL
WP_156919681.1  102 QNLLKKDDRFIIAGPALVNSQVLVIRKD------GNPKR--IAYSHKREYQRAFIQKRFGMDCEAIPVMPTVLPHVYEQGNVDGIVIDILKGITLKGKLLRLAGDDTDIV
WP_146164898.1  102 AELVAKDSRFWEVGPVLANSVVIVTRDP--------EKVRKVGIAQHREYQVSIARKAFGEGVEILPISPMALGVAYESGQVDAAIVDIEDSTNLTGHKIQVGGFLEYVE
WP_028329265.1  101 FQLLSIDTNYYIVDSIVNGSNVLVYYDEIYDH-NIEPKN--IGYMNKR-LLQEELLKSTYSNSNYYPMLATALPYALERKTVDAIFIDIMLALKTKNTRF--KKINTEKP
WP_014932783.1  101 FQLVSIDTNYYIVDSIVKGSNILVYYNESYDDDNVKPKN--IGYMNKR-LLQEILLKEKYTNSNYYPMLSTALPYALEKKTVDAVLIDIILALKIKEARF--KKIDTEEI
WP_147736607.1  102 AKFLGVDTNYFIVDSIVRGSNILVYFDESYKN--VDPKN--IGYMNKR-LLQESILKNVYPNASFYPILSTALPYALEKKEVDAIFTDIILALKIKNAKF--KMINTDKV
WP_208873081.1  104 QKFLASGVPFRVFGGLVKNANVLVARDV------AHPRT--VGYMNGRDLQKQAVFANLGQEVELHPITAPALPYALECGAVDAVVLDAADAIRLPQYRN--RALSCDAP
WP_039879528.1  103 QKFLGTGAPFRIFGPLVKNANVLVSRGN------NTPRS--VGYMSGRVLQYEVLVRTLGRTAELHPIAAPALPYALECGAVDAVVLDAADAIRLPQYVL--RPLPSDAP
WP_009214282.1   95 EKLRSLREDYVELGVIARDTDVLALAEAG-------REIRRIGYMNQRDTQRRSLEAAY-PSAELFPMSPSALPTALFGGEIDAAVLDLGTALSLSLPLR---KLGEGTV
WP_195838058.1   17 LHLVRANADFEIYGPVIMNGEVIAYKG-------DVSNIHRLGIGQKREHLHRLIKKNYNHIDEIVEIQPASLAYSLEGEQIDGAVLDISKAPLLHGFKF-ASLSEDDYI
WP_216560655.1   17 LHLVQANDDFEIYGPVIMNAEVIGYKE-------DVESIHRLGIGHKREHLHKLATESYKQVKEVIEMSPISLPYSLEGGQVDGVVLDITKAALLKDINF-VPLSKDDYI
WP_246565990.1    3 LHLVRANDDFEIYGPVIMNAEIIVYKG-------DMENIYRLGIGQKREHLHKLAKESYRQVEEVVEMTPVSLPYSLEREQIDGAVLDLTKAALLQEFNF-VPLSENDYI
WP_216542239.1   10 LHLVRANDNFQIYGPVIMNAEVIAYKG-------NINNIYKLGIGQAREHLYTLARDSHKQVREIVEISPISLPYSLEIGQIDGAVLDISKVSLLPGFTF-TPLSKEDYI
WP_169300993.1  111 LHLVRANEDFVIYGPVIMNAEVIAYKD-------ELSEVERLGIGQKKEHLHRLAEKSHNQIKEIIEMSPSSLPYSLEGGLIDGAVLDTTKAALLPAFNF-VPFSEDDYI
WP_212381464.1   17 LHLVRSSDDFEIYAPVIMNGEIIAYNK-------ELDNMYKLGMGQKREHLHKVSKENYPQIQEILEMQPISLPYSLEEGQIDGAVMDVTKAALLPKFNF-ASLSENDYV
WP_198306502.1   17 IHMVKANDRFVIYGPVIMNSEVLAYKG-------NIVDQQVLGIGQDRKHIWEQVRKSYPNIKETQEMVVAALGYTLENKQIDIAAIDVSKAALLSKYNF-APISQEDYI
WP_120388142.1   17 AHLVDQDDRLMIYGPVIMNAEVLAAKK-------ESEKITRLGIGQQRKHLQGLIREAHPEIGEIVEMSPRILPYSLSDGQIDGAVLDVTKASLLSGFQF-APIAKHDYI
WP_239591073.1  101 AHLVEEDDRFVIYAPAIMNAEVVAYQG-------DIQQIKTLGVGQQRLYLQNLAKEAYPQIETVEEIKPNMLPYTLSTGEIDAAVMDVTKATLVPEFSF-APLSDHDYI
WP_138307482.1    3 YHLIE-DKRFMIYAPVIMNSDVMIYSGT------DMEHVSHIGISQGRRHLSKMVSEAYPQITEISEISPRVLGYSLETGQIDAAVMDVTKAVLVPGVTI-APISDEDYI
WP_242996699.1   12 VHTVEQNEHVEIYGPAIMNAEVISYKN-------EWEKVKTVGISQGREQEKQVAQETYPQIESFQDVTQKGILYTLEDGQVDAVIQDVTKAAKVPQYRN-KPLSAHDYI
WP_242872717.1   10 RHTVEENENVMIYGPVVMNAEVISYKD-------SWENVKTVGISQGRQTEKTLAEQSYPQIQEFQEVTQKGLLYTLENQQIDAVIQDLTKAADIPEYPY-KPLTDGDYI
WP_016219572.1   17 KHTAQANENVMIYGPAVMNAEVICYKD-------DWEDVKTVGVGQGREQEKRLAEENYPQIEAFREVTQKGILYALEDRQVDAVIQDVTKAAQTSHYDS-LPLSENDHI
WP_005334056.1  100 KHTIENNEDVEIYGPVLMNGETIVHKK-------DWADVKNVGITQGRQQEKALAKAAYPQIEEFDEITQKGILYAMEDEQVDAAVLDITKAAEVSDIPT-MPLSDNDYI
WP_004608618.1   99 RHTVMQNQDVEIYGPVIMNGETIVYKD-------HWENVENLAVTQGREKSKDLAQETYPQIQGFNEITQKGILYALEDEQVDAAILDITKAAKVPDYRN-MPLSDTDYI
WP_066572758.1  100 KHTVEQNQDVELYGPVIMNGETIVYKK-------DWEKVGNLAVTQGREQSKALAQETYPQIQGFNEITQKGILYAVEDEQVDAAILDVTKAAQVQEYKD-KPLSGTDYI
WP_087258574.1  107 MAIVNQSEGFSIYSPVVMNGEVLGYWT-------DPEDIRTMAIPMKREHLAELVREEYPWVESIEEVSSTSILYAFGDQQVDGAVMDIARAFQMPEYQY-TRVCDQDYV

WP_245398079.1  198 TYVLVVSKKFKKSKAYSDFIKLYNKSVDALNNE-STFKN-TLKEYKN----IRN--IDMEEIRNW--KLKFLKIKEID------------------- 265
WP_216467279.1  111 SYVLVVNKDFKDTDSYAKFIRFYNEAVKDLENG-KVFKK-AIENYKG----IDISNKEMEEIINW--KVKFLQIEE--------------------- 178
WP_027702356.1  116 SYVLVVSEEFKQNDLYKDFIRLYNEAVYDLEDR-EVFKK-AIENYKN----VDISNKEMEEIVNW--KVKFLPIKE--------------------- 183
WP_239591062.1  197 TYVLVVNKTFKQSKSYKKFLKLYNKSVNDLQNK-AIFKN-ELENYTD----IKISDKDLGEFKKW--KLKFLQIQN--------------------- 264
WP_239591063.1  197 TYVMVVNKSFKQNKLYDQFIKLYNQSVKELNDE-NLFKK-ELEKYTD----TEISKEVWGEFKKW--KLKFLQIQK--------------------- 264
WP_026901402.1  116 TYVLVVSKAFKNSDSYRVFKNVYNKSVLDLKDK-NNFKK-QLQSYTD----KNISKEDMGEFEKW--KLKFLPIEN--------------------- 183
```

```
WP_245398079.1  198 TYVLVVSKKFKKSKAYSDFIKLYNKSVDALNNE-STFKN-TLKEYKN----IRN--IDMEEIRNW--KLKFLKIKEID-------------------- 265
WP_216467279.1  111 SYVLVVNKDFKDTDSYAKFIRFYNEAVKDLENG-KVFKK-AIENYKG----IDISNKEMEEIINW--KVKFLQIEE---------------------- 178
WP_027702356.1  116 SYVLVVSEEFKQNDLYKDFIRLYNEAVYDLEDR-EVFKK-AIENYKN----VDISNKEMEEIVNW--KVKFLPIKE---------------------- 183
WP_239591062.1  197 TYVLVVNKTFKQSKSYKKFLKLYNKSVNDLQNK-AIFKN-ELENYTD----IKISDKDLGEFKKW--KLKFLQIQN---------------------- 264
WP_239591063.1  197 TYVMVVNKSFKQNKLYDQFIKLYNQSVKELNDE-NLFKK-ELEKYTD----TEISKEVWGEFKKW--KLKFLQIQK---------------------- 264
WP_026901402.1  116 TYVLVVSKAFKNSDSYRVFKNVYNKSVLDLKDK-NNFKK-QLQSYTD----KNISKEDMGEFEKW--KLKFLPIEN---------------------- 183
WP_239591072.1  196 TYVLVVNKEFKKTDNYEIFKKLYNKSVDEISDK-NKLKK-AIENYKN----IKLEKKDMGVIENC--KLEFLKIN----------------------- 262
WP_187296238.1  118 TYVMVVNKDFEKKDLYKKFIKSYNESIIDLKSE-KLLKK-HLSSYID----TTISDEEVREWKKL--NVKLEPIIR---------------------- 185
WP_164952401.1  118 TYVLLVRNDFKKNPVYKKFITAYKKSVKELESE-DVLLE-ELKGYVN----ISITERWLSEWRRW--NINLVPIIPQTNLVN---------------- 191
WP_251861114.1  198 SYVLVVKKGFKKNKRYKIFTKNYNKSVMELKNE-DKIKT-SIEDYKK----IKLSKDEVEQWLKL--RVKFFLIP----------------------- 264
WP_239591069.1  198 SYVLVIKKKFKKDKKFKEFITNYNKSVTELKRK-DRIKN-SIEDYKK----INLSNEEVEQWLKL--KVKFFLIPQQEY------------------- 268
WP_050355227.1  196 TYVLVVRKDFKKDARYKEFIENFKIATTKLNNE-EILLN-ELKTDKE----NNFGRKEVENWKNL--KVKLVYIISQEH------------------- 266
WP_239591067.1  192 TYVLISNVEFMKTKEFKKFVRTYNESLEELMTD-DKLLEKHFYNYTK----TNLEKGGLEKWK-----VKLLNIGED--------------------- 258
WP_090043123.1  118 SYVMVVNKKFRESKEFGNFIEIYNDGVDEINNDYNKLFHAIKLIYNT----GEFSEKEANLWKKW--KVEFLKIQ----------------------- 186
WP_239591070.1  207 SYDMVVNKKFLESNDYREFIETYNESIEDLKYD-EKLYEALRLMYKE----GEFTERSFDLWKEW--NIKLQKLH----------------------- 274
WP_239591071.1  198 TCVLVVNKEYKKSEDFREFIKLYNKCARELSNT-NVLAK-QLEKYKD----MELSPADRHTLGEL--HIKFLKLSD---------------------- 265
WP_239591068.1  199 THVIVASKSFIEREDFKGFVELYNESVNELAKP-KTFKR-AFEDYKG----AALSDKDYEFIKQA--NIEFVQIEP---------------------- 266
WP_242838414.1  107 TFSIIVKKEIKDKEEFKRFISAYDSTIDFLQKEDNFLQSISENRKKD-----YVNERRRKEWKML--NMKMLKLKE---------------------- 175
WP_239591065.1  198 TFVLVAVKGYKDSSDFKAFAKVYNEAVDILNEDVAKRNSHIARYTDL-----------KEVDGETW-KVKLLKIKK---------------------- 261
WP_143156824.1  220 TYVLVANKEVLNLKEFKEIMQAWDKAAHELNEV-EKLSAFLQQNVSN---GDETKGVEIEIWKKA--GVKFLSPLSDQK------------------- 292
WP_093314491.1  209 TYVLIARDGIMETQEAQFFLELWQEAMDELKDM-ETLQTAIDHYMKKEKTGKETTGKEAEQWMNL--GMKLMNPMEK--------------------- 282
WP_073613908.1  202 TYVLVVRKSIAASSGFTALMADYDRAVQEAAHD-ERLLYLLKRYGSE-----EYTEGEVAQWKKL--HVRFVSPLGNRQSG----------------- 274
WP_214081966.1  198 TYVLVVNKSFKSEPRYQQFIDLFKAAVEELNDP-AVLIKEKCHYP-----------------------LLYVHI------------------------ 247
WP_134213589.1  198 TYVLVVNKSFKKDSRYHEFISLFRVSVEELNKP-DALLEEIMKYKNI-----EFTREEVEQWNRL--GIRHVFTTPGIRG------------------ 269
WP_134219406.1  196 TYVLVVNNSFKDDPRYQEFLDLFQQSVEELNNP-DILIEEIGRYKNI-----DFSREEVNQWNRL--GIKHVFTIPEMRG------------------ 267
WP_012031744.1  197 TYVLVVSKRFKEDPRYREFISLFRESVDELNNP-DILRKEIGRYKNI-----ELSQEEIETWNRL--GIRYIFTTPEM-------------------- 266
WP_011700728.1  200 TYVLVARKDFTASPVHRQFMERYERVVRELGDA-ENLARALEGYKAV-----RWTDREVEEWKRL--NVRFVFPPKAGA------------------- 270
WP_163337278.1  185 SYVLVVRKSFRKSLAYTQLLTAIEKATYKLAQP-DELSRVVTQYKNK-----KMGTKEIELWQKM--QVRFVIPTSMLIASLKSRELNLQNTENVVTN 274
WP_031483264.1  197 SYVLVIRKDLQITPLYSRIIKAFENAATKMSNS-TELQAAVLKYKKT-----TLNDEEVKLWQTL--RIRFVSPISMPISLPKL-------------- 272
WP_156919681.1  204 TYVLVVSKRFSRSANFSRFSVALQDSIANLNDIGTLTKVLDEQKKMT--------AAEVKQIATL--SIRFVSPQLHGKR------------------ 273
WP_146164898.1  204 TYVLVVRQNIP---DLGWLLSAFTAAAIELNNTKTLQSAIDGYSRVT------TTGNEADIWKQQS-GIVFLPIVASCNCDPR--------------- 276
WP_028329265.1  205 DYYLVVSKKLKDSAVLNDFIKIYNESVDDIEND--TILSNVLISYLD----ISNSEGEMNIWKTM--GVKFHKIKADL-------------------- 274
WP_014932783.1  206 DYYLVASKKLKDTLMLNNFIKAYNEAVNDIENE--TILSNMLTHYLD----ISNSQGEIKTWKTM--GVKFLKIKAES-------------------- 275
WP_147736607.1  205 DYYLVASKKLKGSDTLNNFIKAYNNAVDDIQND--NILSNMIGSYLG----ISDSKGETDTWKKM--NIRFNKIETKL-------------------- 274
WP_208873081.1  204 SAVLVVHEALIGSTDFKEFVGHYNDTVGRLEA---SLCEELLPQVLG----VECGEETSRAWHKM--NVQLLTLPASTFPPEEG-------------- 278
WP_039879528.1  203 TAVLLVHNDALQGQNFKNFIRFYNDTVRRLEG---SLCNGLLTDVLH----VERREETSKAWQKL--NVQLLTLP------EEGSI------------ 273
WP_009214282.1  194 SAVLVAREDCLKKKAFQKLLSDYEKKLPELSNEREITGLLSHYMRED-----EITKEKREQWKAMGPSFESLREGREGRGG----------------- 269
WP_195838058.1  119 SYSLIVRKDVINTKVFQEFLMAYNKTIEELNKVKTLKEYIG---------------MPKEFWDNT--NIKFLSLE----------------------- 176
WP_216560655.1  119 SYSLIVKKDIIGTKVFEDFITAYNNTVQELNQTEILKEYIS---------------MREEFWQNK--NIKFLSLE----------------------- 176
WP_246565990.1  105 SYSLVVRKDIIGTKVFEDFLTVYNKTIEELNQVGILKEYIV---------------MTEELLNNT--KIKFLSLE----------------------- 162
WP_216542239.1  112 SYSLVVRKDIIDTKRFKNFLITYNKTIEELNQIEKFKEHIK---------------IPDNVYNDI--KIKFLSLE----------------------- 169
WP_169300993.1  213 SYCLVVRKDIIETEEFKGFLDSYNKAVDEFNQEEVLQNLIN---------------GNKALWDSI--NIKFLNLE----------------------- 270
WP_212381464.1  119 SFVLVVRKDIIDTRAFRDFLDVYNKTIEELSQTETLITHMG---------------MTKEFWDKA--KLKFLSLE----------------------- 176
WP_198306502.1  119 SYSLIVRKDLIDTEAFDNFLVSYNKAVEELNKKETMISVMG---------------MTKEFWDMV--NLKFLEL------------------------ 175
WP_120388142.1  119 SYVLVIRKDLIGSAPFERFLEAYRNAVDGL---QEVYFDPA---------------LPKEY-------TRFLYLD----------------------- 168
WP_239591073.1  203 SYVLIVREDLIGTEKFDQFIKSYNRSVEQLKDGQKLQQDLG---------------MDAQFWK----QVKFLTMDSYE-------------------- 261
WP_138307482.1  105 SYVLVVRRDVADTEPFLKFVDAYNKTVRALNRTGTFSEVIN---------------ADGILPDSV--HIKFLGI------------------------ 161
WP_242996699.1  114 SYVLVVEKEFAKTKAFEDFVISYNQAVEKLNQPEYLAEKLG---------------VQKEWIQDK--QIEFLPLELRQ-------------------- 174
WP_242872717.1  112 SYVLVVDRDFVETEAFADFVVSYNRAVEKLNDPEYLAEKLG---------------VEEDWLQGT--AVEFLPLQEVER------------------- 173
WP_016219572.1  119 SYVLVVDKDFAGTEAFADFIKSYNEAVRKLNRPAYLSERLG---------------VKESWLEDK--SIEFLTLEESEE------------------- 180
WP_005334056.1  202 SYVLIVDKEFEQTEAFRNFIESYNKAVEKLNDPFYLAEKLK---------------VSKEWVEEK--QIKFLPLEAKN-------------------- 262
WP_004608618.1  201 SYVLVVDKEFEKTEAFRDFIDSYNRAVSRLSNSDYLAWTMG---------------VKEEWLENK--NIKFLTLEGTGVD------------------ 263
WP_066572758.1  202 SYVLVVEKEFEQTEEFERFLKSYNRAASRLNDKGYLAEVLG---------------VEISWLDSK--NIKFLTLEETGED------------------ 264
WP_087258574.1  209 SYCLVVRDEIVDTPQFQTFLKYYNETAEALNDKANLQALYG---------------MDDAFWDSV--NLKFLYLPEAGQE------------------ 271
```
