## Supplemental Figure 7 for "Eight Unexpected Selenoprotein Families in ABC transport, in Organometallic Biochemistry in *Clostridium difficile* and other anaerobes, and in Methylmercury Biosynthesis"

methylmercury formation protein HgcA

```
WP_250697401.1     1 -------MKVQKTGKVEWSNVKLATPAGCGPAAGNGLEK----------------PPCUGP----RISAGGGA---IDETVPGFLGWLTTDAGRFPQVASELAWTDRLGG
WP_250697402.1     1 -------MKIREIRTITKAECAPVN-TGCVPTADNDPNK----------------PPCUGP----PTSAGGGT---IDETVPGFLSWLPTRAGRVPRIATELRFADRLGT
WP_250697418.1     1 -------MKITSIRRVTEADRTPAA-SACGDVSGDGQDK----------------PPCUGP----PTSAGGGA---ITDKVPGFIEWLETPAGRVPRITSKLGLKDHLGA
WP_250697419.1     1 ------MEKPVKMRQFKKIEVAKPKAAPGGCVPSADPDA----------------PPCUGP----RTSAGGGT---IDETVPGFEGWLDTPAGRVPRIRTALTVADHLGA
WP_252396620.1     1 ------MSEETQMRRFKKIEMKKPASSSCASELGANPDA----------------PPCUGP----PTSAGGGE---ITDQVPGFLGWVAAGSRRAPRISGSLTWADHWGA
WP_250697417.1     1 MHVASKTKEHYMSEQLKTCGEINAPSAGCGTPAG-ETDK----------------PPCUGP----RPPAGGGI---IDETVPGFTGWLETQAGKVPQISSDLRFDDLIGR
WP_250697413.1     1 -MVTVKEKIMERQTCGDESREKAEGTGASGCGGGAAGDA----------------PPCUGP----KTLAVGGE---INENVPGFVQWLATPAGRVPQVSSELGFADRLGA
WP_250697414.1     1 -------MTQKPLSAGGIRSVKPVEPCSSVSPEGPEIDK----------------PPCUGP----KTPAVSGS---IDENVPGFIRWLETLVGRVPVISSTLRFRDRAGA
WP_250697415.1     1 ----------MAYRVRKVTVIKQGRGGFRPGQAGADLAK----------------EPCUGP----KTPAVSGT---INENVPGFIRWHETPAGRVPIVSTEFRFADRLGA
WP_108102289.1     1 -----------------MKNRNGETSPCCCTEVEPHVER----------------PPCCGP----PATVGGGE---IDETVPGFLGWLSTTTGRVPRISSELTINDRFSA
WP_004098500.1     1 ----MEKG---------------PAAGGCSCSCGETQNSE---------------THCCNQ----RPMANIS-----EKPMPWIIGEIETSIGKIPQVSTEYASSDRFGS
WP_004626173.1     1 ----MENKSCCTSGNARLNTGDRKSVKIRPRLKAANKNCTGVS-------GQKDNTCCCGG----SDNCSSEPVTNYDRNIPWITGEIETAAGKIPQVATRLLRSDILGA
WP_020491038.1     1 ----MLYLGEISFD-PNQNN---RNANNKAELKIFDPQ-----------------PCCCG------DDSLAGSVTVYDKECGWITGEIKTPAGLVPQVSTNLTALDILGG
WP_010243111.1     1 ----MEDKRCCNSNKINSKN---KVAGNKPKIRVSSQSSSLPY-------TAEKGSCCCGG----SDQAEAEPITQYDKKDNWITGEIKTPRGKVPIVSTQLAFSDRLGG
WP_207655107.1     1 ----MGNNNCCNSENKLDKNAKVKIVRIKPKSIGSLTSSKG---------GKTKSSCCCGN----GSSASCEAITKYDLKEKWIIGEISTPAGMVPRVSTKLTFEDTWGT
WP_245693461.1     1 -------------MTETDPCCSPAPPGDSLPMAEACCPPPGS-------TALAMAEPCCGSAAEAVGPRWADRERPGYRIWPFVDEWIETPVGDVPVVSTRLGKGDVLGR
WP_155317484.1     1 -------------MTSCCPEEKNQVPRPDGTSHESTACGLKP----DFPILGKDAGACCGG---QAPETGSRHETPGYRTWGFVEDFVETPAGRIPRVHNHWNREDRTGT
WP_226860105.1     1 ------------------------------MQDDLMAFGERA-------ARDLGKPSCCG-----PSPARDAREMPGYRLCGYVDSFTETPAGKVPVVKTRLDGSDHLGG
WP_236890013.1     1 ------------------------------MQDDLMAFGEKA-------AGAVDEPCCCGP---APSPRSDGRELPGYKLCGFVASFMETPADTVPVVKTRLEREDHLGT
WP_022666783.1     1 -------------MDKTKQKKSEAPGPSGFSGRNLSILPSMP-------SQENSDLSCCGSG--VREPALPGDQKPGYALCSHVEDFLPTHAGRVPIIKTRLNREDLFST
WP_250697411.1     1 --------------MKGLD-DKPLTGTTPEPLEPLALMAPPD------VPARD-DAPCUGP---RPDPRAGVFEKPGYAVEPFVDGFARTAAGPVPRVRTYLRRDDRVGD
WP_250697410.1     1 -------------MEKQKK-EMQPLDLQPLDMAPLEFAPLEG------CPSQD-DAPCUGP---KPDPRAGVHEKPGYSMESYVDGFVDTPAGFVPRVRTRLSRRDMVGT
WP_250697412.1     1 -------------MNKSKHSDLGPVFVAPPDARPNVCGPQII------LGTPTGDAPCUGP---KPDPEAGVHERPGYVIEPYVDGFVETSAGPVPRVETAVSLRDKLGT
WP_200881740.1     1 -----------MSELKPPQDECCAAGAACCPPENPTALQPFT------MAAAS-DAVCCGE---PPGPRSNPDEKPGYELQHYVDSFLDSPMGQIPRVKTGLEPADFRGT
WP_252409333.1     1 -----------MDKLEKRDTELGKAEAIVPLVEGAFCMIRR--------PSGCGEEPGUGP---RPGPGFGPENKPGYTVCHFVRDFARTPVGHVPRVMSYMTPRDWAGS
WP_246298930.1     1 ----------MMPLTGKKRTTAAADPCCCSDAPGRGDAAASSCCDGSASGGEDSGQPCCGG---PQAPAASPMARPGYELHPFVETFRTTLAGPVPVVRTSLDRRDFRGT
WP_205232271.1     1 -----------MAGTKSGREEKDFSEGSVIELAEVQTAAAP-------------QEVCCGG---APPPKTNPLERPGYTLKPFVRAFIKDSRSDVALVKTELDFSDHIGA
WP_250697420.1     1 ---------------MS---LQPLTALRPAGNAGGCTTG--------------DDAPCUGP---KQQPRSGVHERPGYRLDPFVVDFLETAAGPVPQVATRPDRHDRLGA
WP_250697404.1     1 ---------------MQP--INPCPASPQPLP---LFAPS------------PPEAPCUGP---KPEPRSGVHERAGYSLCPFVTGFIDTPAGPVPQVTTTRSSRDLWGT
WP_250697403.1     1 ---------------MQ---SGPLPDLRASAP----PSGK------------ESE-PCUGP---PQQPRSGAHHRAGYALSPWVSGFIDSPLGPVPRVTTAPLWSDRLGT
WP_250697406.1     1 ---------------MQ---NTIHTQCSGRNETDMCGQGDHA------QKHFDNQAPCUGP---NAEPRAGVHEKAGYSIDAYVEGFIDTPVGPIPRVATRSQLRDRVGR
WP_250697405.1     1 ---------------MH---DGKMIELQKLQPKPVIPIAA------------EREAPCUGP---KQEPRAGVDQRAGYCIEPFVEQFMATSIGSVPKVFTIPSVRDRYET
WP_252405647.1     1 ---------------MSDALQQQIFPLLHTEQNKPGSPAPLQ------VPEEIPAGPCUGP---KPPPRSSGYARAGYTICPFVEGFTETPAGPVPRVATSLSIRDRLGT
WP_252405646.1     1 ---------------MNSFLQQHIEASP--VANRP------Q------IPPEVTGETCUGP---PSPPRSGGDEKAGYRLLPFVDRFIDTACGPVPVVATTLSRRDLLGT
WP_250697409.1     1 ---------------MNPGPQHHMNLQP--FPTLPMASPLAP------PADHGDAAPCUGP---KPDPRSGVHERPGYRIVPYVADFIATAVGPVPLVSTTLTWRDHLGT
WP_250697408.1     1 ---------------MTDQPQHFITPISAIKPLDSHVANPFS------PPTEPEAGPCUGP---ETDPRAGEHERAGFILAPFVAGFVTTPVGAVPQVATALTLRDHLGT
WP_250697407.1     1 ---------------MRSEPQQFITPIQDI---------PIQ------PPPEIDSGPCUGP---ETYPRAGEHEKAGYTIDPFVHGFIQTARGSVPQLKSELSGRDHLGT
WP_252396621.1     1 -----------MKTNSHEVTPLSSAGLSPSLLDEISVHDLPG------FPPPADNAPCUGP---KTESGFGVYDKPGYTISPFIESFYETPGGPVPRLRTHLSLRDHGGT
WP_252396623.1     1 -----------MNDLK----ELNDVMFNQPMMA----------------PSSQTEEICUGP---QPDPEAGVYERPGYDICGFVEEFMETPAGTVPRISSKIRFKDHFGT
WP_250697582.1     1 ------------MVSPAALTAKDFAAFTTNISCGTNGSPCGAG-DVIAQGAPADDAPCUGPR--NELPPEAAGERPGLRSWDFVSGWLQTPAGLVPRVRFDWAASDRLGA

WP_250697401.1    81 WKARWGIGRMSYLIPAGLYAVGQPTPADPVVVTANYKMSYDLVRRSLAGRNVWLLVLETFGINVWCAAGKGTFGTDELVRRIGATGLAQVVTHRRLLLPILGAPGVAAHE
WP_250697402.1    80 CKTRWGIGRMEYMVPPGLYAVGTPTPDDPVVVTANYKMSYDMVRRALAGRNVWLMVLETFGINVWCAAGKGTFGTDELVKRVAATGLATVVNHRHLLLPILGAPGVAAHE
WP_250697418.1    80 CKVRWGIGRMSYMVPPGLYAIGRPTPEDPVLVTANYKMSYDIIRHTLSERNLWLLVLETYGINVWCAAGKGTFGTGEVVRRVAATGLARVVNHRRLILPILGAPGVAAHQ
WP_250697419.1    82 CMARWGINRMDFLVPPGLYAIGAPTPEAPVLVTANYQMTYDLVRRALAGRNVWLLVLETFGINVWCAAGKGTFGTTELVRRVAESGLAKVVQHRRLTLPILGAPGVAAHE
WP_252396620.1    82 CLARWGINRMTFMVPPGLYAIGTPDAQAPVVVTANYKMSYDIVRRALSGRNVWLLVLETYGINVWCAAGKGTFGTGELVRRIEASGLAQAVSHRRLILPILGAAGVSAHR
WP_250697417.1    87 WKARWGIGRMSYLVPPGLYAIGRPTGSDQVVVTANYKMSFDIVRNELAGRNVWLLVLETHGINVWCAAGKGTFGTEELVHRLEATGLARIVTHRRLLLPILGAPGVAAHQ
WP_250697413.1    87 WKARWGIGRMSYLVPPGLYAIGDPGADDPVVVTANYKMSYDLVRRALAGRRVWLLVLETHGINVWCAAGKGTFGTVELVQRIWQTGLHQVVQHRLLLLPILGAPGVAAHV
WP_250697414.1    81 WKARWGIGRMSYLVPPGLYAIGTPGPADPVAVTANYKMSYDLVRQAMAGRDIWLLVLETYGINVWCAAGKGTFGTGELVRRLEASGLARVVTHRTILLPLLGAPGVKAQE
WP_250697415.1    78 WKARWGIGRMSYLVPPGLYAIGSPGPEDPVVVTANYKMSYDLVRRELSGRNVWFLVLETFGINVWCAAGKGTFGTDELVRRVGMVKLATVVKHRALILPILGAPGVAAHQ
WP_108102289.1    71 WKLRWGIGRMSAVLPPGLYALGTPEGQDPVVVTANYTMSYDLVRHALAGRNLWLLVLETFGVNVWCAAGKGTFGTEELVRRIEKTGLSKVVDHRLLLLPILGATGIEAFK
WP_004098500.1    68 WKARWGIGRIDYKIPPGLYGVGNPDADSLVLVSANYKMSFDRLRSELKGLNLWILVLDTKGINVWCAAGKGTFGTDELVNRINQVKLPEIVTHRTLVVPQLGAPGVSVHT
WP_004626173.1    96 WKVRWGIGRMNYKINPGVYAVGIPDRASPVLVTANYKLTFDALRKELSGMNVWILVLDTKGINVWCAAGKGTFGTRELVNRISRARLREIVEHKTLILPQLGAPGVSAHE
WP_020491038.1    80 WKARWGINRMNYKINPGLYAVGNPDEHSEVLVTANYKMTFDALRKELAGLQAWILVLDTKGINVWCAAGKGTFGTVELSSRIEKTRLARIVSHRTLLLPQLGAPGVSAHQ
WP_010243111.1    93 WKARWGINRMNYKINPGLYAVGIPDNTSPVLVTANYKLTFDALRKELADINAWILVLDTRGINVWCAAGKGTFGTREIVTRINKTRLHEIVSHRTLILPQLGAPGVSAHE
WP_207655107.1    94 WKVRFGINRMNYKINAGIYAVGTPDNTSPVLVTANYKLTFDSLRKELEGLNAWIMVLDTKGVNVWCSAGKGTFGTKELVNRISKVRIPQIVSHKTLILPQLGAPGISAHQ
WP_245693461.1    91 WAMRWGIGRDRYRIAPGLYAVGRPTPDSPVVVTANYKMTFDLVRKDLGGVNAWILVLETWGINVWCAAGKGTFGTDELVARVRKAGLDRVVSHRTLVVPQLGATGVAAHE
WP_155317484.1    91 LCARLGIGRDSYRIAPGLYGVGAADAASPVMVTANYKLSFDTLRTSLADTSAWILVLETHSINVWCAAGKGTFSTREVADRVKAAGLEKVVAHRRLILPQLSATGVAARQ
WP_226860105.1    69 LAVRCNLRRSRYAVAPGLYAVGSPSDTSPVLVTANYKLSFDAVRRALDGMDAWLLAADTRGVNVWCAAGKGTFSAGEVADRVKASRLAEVVTHRRLILPQLSASGVTARE
WP_236890013.1    71 LAARWNVGRSNYKVTPGLYAVGTPDKESPVLATANYKLSFDAVRRELSGINAWLLVVDTRGINVWCAAGKGTFSAEEVAEMVKRTHLEKVVSHRKVILPQLCANGVTARD
WP_022666783.1    89 FYVRCGIGRDDYRVAPGLYAVGKPDRESEVLVTANFKLTFDHVRKNLKEVDAWILVLDTWGVNVWCAAGKGTFSTGELVSRIETSCLDKVVDHRRVIVPQLGATGVSARA
WP_250697411.1    86 LRARLGTNRHDFKVVPGLYCVGEPDRTSPVLVTANYKLTFDTLRERLTSIDAWLLVVDTRGINVWCAAGKGLFTASEVAFSVNAVRLHQVVEHRELILPQLAATGVAARE
WP_250697410.1    87 ALARLGACRVNYKVVPGLYCVGDPTPDSPVLVTANYKLTFDAVRRELDGVDAWLLVADTRGINVWCAAGKGLFSTDEVVYSVRSARLDEVVSHRKLILPQLGATGVAAHH
WP_250697412.1    89 VKTRLGIGRYNYKVVPGLYCVGTPGPESPVLMTANYKLTFDALRKELGGLDAWILAADTRGINVWCAAGKGLFSTEEVVRRVNRSRLAEIVSHRELILPQLSATGIAAHQ
WP_200881740.1    90 IKARLGLRGTEYKIAPGLYAVGSPGPDSPVLVTANYKLTFDTLRRELTSFDGWLLVIDTRGINVWCAAGKTLFSTQEVVYRIQQCRLSKVVQHKKIILPQLAATGVSAVS
WP_252409333.1    89 AFARLGVIRDNYKVVPGLYCVGFPDEDSPVIVSANYKLTFDALRRELSGIDAWILVVDTRGINVWCAAGKNLFSSEEVARVVRAARLSEVVSHRRLILPQLAATGVGARK
WP_246298930.1    98 LGARLGYGRDGYRVAPGLYAVGRPGPDSPVLVTADYKLSFDALRRELSGIDAWILVLDTRGINVWCAAGKGTFSAAAVARLVEASGLARVVSHRRLIVPQLAAPGLDARD
WP_205232271.1    84 LLARLGINRDEYRIAPGLYGVGYPGQGSPVIVTANYKLTFDFVRRELAGLDCWILVLDTCGINVWCAAGKRTFSTEEIVYRIKETGLSERVNHRRVIVPQLGASGIAAHE
WP_250697420.1    76 FLTRLGPIRNHYKVVPGLYAVGTPTAEAPVLVTGNYKLSFDALRFSLTGIDAWILVVDTRGVNVWCAAGKGTFSTAEIVHSVKHTRLADFISHRRLIVPQLGATGVSASE
WP_250697404.1    76 IGARIGIIRRNYKVAPGLYAVGSPGNQSAVLVSANYKLSFDALRHQLGGLDAWLLVVDTRGINVWCAAGKGTFSTWEVIDCVQRYRLNEVVAHRELILPQLAAPGVSAQA
WP_250697403.1    73 IGARLGAIRNTYLVAPGIYGVGNPQADSPLLVTANYKLSFDQLRFQLAGIDAWILVIDTRGINVWCAAGKNTFSTAELVYSLDHYQLSRLLSRRRLIVPQLSAPALCAEE
WP_250697406.1    84 LFTRIGIIRDNYRISPGLYAVGTPHRDSPVLVTANYKLSFDLLRFELAEVDCWILVVDTRGINVWCAAGKGTFSTDEVVLMVKQTRIAELVSHNTLIVPQLCAGGVSSSL
WP_250697405.1    78 VLARVGTIRNTYAIAPGIYAVGSPTSTSPVLVSANYKLSFDTLRFHLAGFSAWILVLDTRGINVWCAAGKQTFSTQEIVMSLERYRVEQLVDHQQIIVPQLAAVAVSAHE
WP_252405647.1    87 LGARTGFTRNTYSITPGLYCVGKPGKDSPVLVTANYKLTFDALRRALPHLSLWILVVDTRGINVWCAAGKKTFSAEEVAYQVKSAKLDQVVAHRELLLPQLAASGVAARE
WP_252405646.1    79 IGARTGFTRDGYSVTPGLYCTGSPDENSPVLVTANYKLSFDSLRKELGSLSCWLLIIDTRGINVWCASGKGTFSAEEIAYQIQRCGLIHVVKHRELILPQLSANGVAAHN
WP_250697409.1    85 IGARSGYYRSNYKVIPGLYGIGRPGLDSPVLVTANYKLSFDALRQVLPGIDAWILVVDTRGINVWCAAGKKTFSTDEVAYQVKSTRLAEIVRHRELILPQLAAVGVAAGR
WP_250697408.1    87 IGARTGFVRNSYKITPGLYAIGQPSPEDPVIVTANYKLTFDAVRSSLATQNLWLLVVDTRGINVWCAAGKGTFSTEEVAYQVRKARLTEIVSHRKLILPQLAASGVAAIR
WP_250697407.1    78 IGVRTGILRNNYKVTPGLYAVGSPSADSPVVVTANYKLTFDIARRNLKVIDVWLLVIDTRGINVWCAAGKGTFSTAEIAYQVKHSDLAKVVNHRTLILPQLSATGVDCLR
WP_252396621.1    91 MFARLG-GRTNYKIVPGLYCVGTPNAGSEVLVTCNYKLTVDALRHDIEGVNAWILILDTHGINVWCAAGKKTFSTVEIIHRVAMSELKCIAPEATLILPQLAATGVNARE
WP_252396623.1    77 FLVRTGFIRTNYRVTPGVHAVGNPGPDSPVIVTANYKLTFDVVRRDLACTDVWLLVIDTRGINVWCAAGKSLFSTDEVIRKIRSSGLEKLVGHRSVILPQLGAPGVAAHE
WP_250697582.1    96 LAVRMGIGRDSYRIAPGLYALGEPDAQSPVIVTCNYKLTFDILRRDLAGHSCWLLVIETYGINVWCAAGKRSFSTAEVAARVRATGLERIVEHRRLVLPQLAAPGVAARK

WP_250697401.1   191 VAGLTGFTVRYATIRADDLPAYLDNGMVTTPAMRELTFTFRERLVLIPVEL-VLALKPTAAVCAVIFLLAALLGGV---PAG-------LWTLLAYAGAVFTGIAVAPLL
WP_250697402.1   190 VQKQTGFTVRYATIRAADLPAYLDNGMVTTPAMRELTFSLRERLVLIPVEL-VLALKSITLIGGLIFLAALILGGD---AAG-------IKALIAWIGSVLTGIVAGPLL
WP_250697418.1   190 VARRCGFSVSYAAIRAADLPEYLNNGMVTTPAMQQLTFTLYERLVLVPVEL-VMALKSVAIIAGVVLLLVAALGGL---TAG-------VTVLIAYVGAVVTGTAVGPLL
WP_250697419.1   192 VQKATDFSITYATIRAEDIPEYLDNGERTTPAMRELSFTLYERLVLVPVEI-VLVLKTLLVLGGGLYLVALLAGGS---GAA-------LTTVLGYLGAVFTGIAVAPVL
WP_252396620.1   192 VKQRSGFEVVYATVRADDLPAYLDQGGTATPAMRELRFNWYDRLVLIPVEI-VLAAKPVAPVAALLYLGARLFWGA---DAA-------AATALAFLGAVLTGIAAGPLL
WP_250697417.1   197 VAQRTGFSIVYATIRASDLPRFLDNGQVTTPDMGKLTFSLYERLVLVPVEV-MHSLVPTAVIILCLFLCGTLFGGA---TSG-------LQAAIAYLGATLAGLVATPLL
WP_250697413.1   197 IKQGTGFGVRYATIRATDLPEFLDNGMVTTPAMRELTFTLRERLVLVPVEL-VSSLPATLAIVAVL-----VAAGL---LAGS----PLAP-AAAYLGGVAAGAAVTPLL
WP_250697414.1   191 VRQRTGFTARFTTIRIEDLPLYLDNGQKATPAMGELTFTAGERLVLVPVEL-VSALKVSLPILAVLYLLAGMAYGF---TAPA----AFFP-LVVWLGGVIAGAVVTPVL
WP_250697415.1   188 VARRSGFRVHYATIRAADLPEYLDNGMVTTPEMRQLTFTTRERLALTPIEL-VTTGKWVVPFVALLVVLSIVPGGL---LSSP----TLVSSLVLFVGTVLAGAVAAPVL
WP_108102289.1   181 VEARSGFRVRYATIRVDDLPEYLDNGMVTTTAMREMTFFMRERLALVPVEL-VMSLKKLAAVSLLLCLVPALMGDP---GAG-------AWAVLAFLGAALGGIVVVPLL
WP_004098500.1   178 VLKLSGFKVVYGPVRAADIPAFISAGLKATAAMREVRFDMFDRLVLAPFEL-VSTIKPALILIAVLFLVNLTAGKE---GVDLRVVSHTLADFIPFLGAILTGTVLVPLF
WP_004626173.1   206 VTKQTGFKVVYGTVRAKDIRSFINSEMKATEEMRQVRFTVYDRLVLTPMEL-VGTFKISMLIFGILFLINLVARK-----------PFGVMDFCAYAGAVAAGCVITPIL
WP_020491038.1   190 VHKQTGFNVVYGPVRAWDIREFIASGRKATAEMRTVKFTAYDRLVLTPMEL-TGTFKISLMILGILFIFNLIRPG-----------AFGITDFYAYAGALIVGCVLTPVL
WP_010243111.1   203 VSTKIGFKVVYGPVRASDIKEFITLGMKATPEMRTVKFNTYDRLVLTPMEL-VNTFKICLMVLGLLFILNLIAVR-----------SFGVIDLYAYMGAILAGTVLTPVL
WP_207655107.1   204 VTKQSGFKIVYGPVRAEDLKEFLEAGMKATEEMREVRFNIIDRVVLTPIEF-LSTFKVAIIVFGIMFLLNLVAVN-----------PFGVIDFYAFMGALLAGCVITPIL
WP_245693461.1   201 VRRRCGFRVVYGPVRSRDLAAFLRAGMKASPNMRRVTFSLWERLVLTPVELRVILMRPSLALAALLLVLGGIGP---GGYSLAGALVRGGAAVGTLAAGVLAGAVAAPVL
WP_155317484.1   201 VKRLCGFEVVWGPVHARDIKAFLENGMKAAPAMRKVTFSLRERLELIPVEL-NHMGRPTLYLIPGLLVLSGLGT---GIFSVSAAVSRGSAAILAYVLAVATGAILSPLL
WP_226860105.1   179 MKTRCGFAADFGPVRAWDLPRFIEAGGCAEPAMRRVTFSLKERLVLSPVEL-SLTFKPLVWMLPVLFLLSGIGP---GVFSFSQALERGPVAWAALCTGILAGAVVTPAL
WP_236890013.1   181 VKKLCGFRAEFGPIRASDLPRFIKNGLKADEEMRRITFTFKERVELTPVEL-SLTGKPLLWLLPLLFLLSGIGP---DVFTFSQADHRGLAAWMAIFTGIVAGAVVTPAL
WP_022666783.1   199 VKKASGFRVVYGPIRARDIPRFLENRRKADPAMRRVTFPLYERFILTPVEI-KIALKPALITAIVLILLSGIGP---GVFSFSGAWERGMAAIAALVTGILAGGFVTPVL
WP_250697411.1   196 VERICGFKVLWGPIRARDLPAFLRNGNKADEAMRGVTFTLRERAALIPVEL-YQLRKPLFAAIPLLFLLSALGP---DLFSPPALWQRGISAVTATLVGALAGSVLVPLF
WP_250697410.1   197 VRKACGFEVAYGPVRAEDLSAYLAAGQRADEEMRSVTFTLKERAELVPVEL-FLIGKPLLLTVLAAFVLSGIGP---GFFSLSRAWERGVIASGITILGILAGCLMVPLL
WP_250697412.1   199 VKKICGFKVIYGPIRAADIPAFIQAGKTADQAMRTVTFTFRERAVLTPVEI-LLLRKALAVVIIATLFLSGFGP---GIFSMDAALSRGIPAVCATLFGILAGAVIVPFL
WP_200881740.1   200 VKKESGFEVLWGPIQARDIRQYVADGCQAGQAMRRVTFTLPERVVLIPVEL-SHLPKPTIWVLLAVFLLSGIGA---GLFSFNAAWSRGLIALVAYAVGILAGAVAAPIL
WP_252409333.1   199 VKKLCGFDVVFGPVRASDLQRYFENGSEATPDMREVTFTLAERAVLIPVEL-VLLWKTMLTALCVAFVLSGVGP---DVFSFSAAAERVGDVALALVAGVVSGAVVVPLL
WP_246298930.1   208 VRRRCGFTVMFGPVRAADLPAFLRAGNRAEPAMRRVTFTLSERLAVAPVEV-YHARKWLWWAALAALAVSGIGP---GVFSLAAVWERGVWLVAAILAGLVGGALVVPVF
WP_205232271.1   194 VRKRCGFRVVYGPVKAVALKAFLENDMQATEKMRTVTFTFGERFELIPVEF-YLFGRKVWWIFPLLFIISGFSP---DIFSPLQGWQRGILVSSAALWGGVCGMLIVPLL
WP_250697420.1   186 VKKGCGFSVIYGPIKADDLPAFLKQGESAGEAMRTVTFPLKERLVLIPVEL-YLQLKQLALIMVIGFVISGVSP---EIFSLSAAWRRGLLLFAATVIGMVCGSALVPAL
WP_250697404.1   186 VRKGCGFTVLFGPVRAADLPAYLGQGNRCDEAMREVTFPLADRAVLIPVEL-FLLAKPLLAMLILGFLLSGISP---TIFSLTAALERGEALLAATGLGMLGGAVLTPML
WP_250697403.1   183 VRRQSGFRVIFGPVRAADLPRFLAQGEVADEEMRTVTFSWQERAVLIPVEF-FLVLKPLAAILISCLLISGLGP---GFFSLEAVLSRGPLLLITTLAAICSGTVLVPLL
WP_250697406.1   194 VKKKSGFRIIFGPVQARDLPEFLERDNTASEGMRAVTFTLMERVELIPVEL-YLQLKTIGFVLIAGFLFAWIGP---HMFSLQAIIQRTLHLGMATLLGLLCGSVFVPIL
WP_250697405.1   188 VKKRSGFTVIFGPVRASALQEFVANGNQADEAMRTVTFSMAERMVLVPVEF-YLLSRQLLVILLVGVALSGIGP---DAFSAAAMFDRGGIFVIITVAAVFSGALVVPLF
WP_252405647.1   197 LKKLCGFRGRFGPIAAAMLPEFLKNGK-ADERMRTVTFSLTERAALIPLEI-CMLWKRLLAAAIVLFVLSGISP---QIFSLEAARSRGLILLLATLGAITAGAVLTPLL
WP_252405646.1   189 LHKLSGFSAKFGPIRASDLVLFLQRGK-ADEAMRSVTFSLAERAVLIPLEI-CMLWKQLAIACVLFFVLSAISP---EIFSLRAGLERFSLLIYATLTAVAAGAVVTPLL
```

```
WP_250697404.1  186 VRKGCGFTVLFGPVRAADLPAYLGQGNRCDEAMREVTFPLADRAVLIPVEL-FLLAKPLLAMLILGFLLSGISP---TIFSLTAALERGEALLAATGLGMLGGAVLTPML
WP_250697403.1  183 VRRQSGFRVIFGPVRAADLPRFLAQGEVADEEMRTVTFSWQERAVLIPVEF-FLVLKPLAAILISCLLISGLGP---GFFSLEAVLSRGPLLLITTLAAICSGTVLVPLL
WP_250697406.1  194 VKKKSGFRIIFGPVQARDLPEFLERDNTASEGMRAVTFTLMERVELIPVEL-YLQLKTIGFVLIAGFLFAWIGP---HMFSLQAIIQRTLHLGMATLLGLLCGSVFVPIL
WP_250697405.1  188 VKKRSGFTVIFGPVRASALQEFVANGNQADEAMRTVTFSMAERMVLVPVEF-YLLSRQLLVILLVGVALSGIGP---DAFSAAAMFDRGGIFVIITVAAVFSGALVVPLF
WP_252405647.1  197 LKKLCGFRGRFGPIAAAMLPEFLKNGK-ADERMRTVTFSLTERAALIPLEI-CMLWKRLLAAAIVLFVLSGISP---QIFSLEAARSRGLILLLATLGAITAGAVLTPLL
WP_252405646.1  189 LHKLSGFSAKFGPIRASDLVLFLQRGK-ADEAMRSVTFSLAERAVLIPLEI-CMLWKQLAIACVLFFVLSAISP---EIFSLRAGLERFSLLIYATLTAVAAGAVVTPLL
WP_250697409.1  195 LRKACGFNGIFGPIQAKDIPTYLHRNRQADEAMRTVTFNTGERAVLIPVEI-CLLWKPFLAATAVLFLLSGIGP---EVYSLSAAGTRGIQALIASLLAVLAGAVLTPLL
WP_250697408.1  197 LKDMTSFRAAFGPLRIADLPRYLSGSIDDFEQMRSITFTAKERLVLIPVEV-CMMYKQLALSILFVVLISGIGP---DIFSARMAISRTWQFILATGLAILAGAVITPLA
WP_250697407.1  188 LKSECGFRGVFGPVRLSDLPEFIANQLQADDRMRSVTFTLKERAELVPVEV-FLSIKPLVLLLLVMLPISGIGP---EFFSTNQAISRIIPFLLATLMGIFAGAVVTPLL
WP_252396621.1  200 VRKKTGFPTIFGPILSKDIPQFLANNLQADTAMRRVTFNMSERAVLIPVEI-TLLWKPIIIGLVLMLLVSGISP---SIFSFHAALTRGYAAFLAGVAGIVAGTVIVPLF
WP_252396623.1  187 VRKATGFKVKYGPVRSSDLKAFIENGLKTTDDMRAVSFTLKERAELIPIEI-VMAWRWILLALLLFVLLGGIGP---WGYDVHAAWSRGWDAFLTGLAGFLGGTALAPLL
WP_250697582.1  206 LRGLCGFGAVFGPVRSRDLPAFLGAGFTAEPSMRRAEFPLGERLTVALVEM-HAARKFMAWTLALCLVLAALGPAWKGGFSVMGLLAVGLGAFAMTLAGFVGGTFVTPAL

WP_250697401.1  290 LPWLPVRSFAVKGALAG---LAWSTGYYLLAGGGS---WSVPVAIAAFLTLPAVSAFYTLNFTGCTTFTSRSGVKKEMRLALPVMGGALAVSALLLVAGLFL--------- 385
WP_250697402.1  289 LPWLPGRSFAVKGGIAG---LGWSFVWFQLAGGAA---WGVPVSLAAFLALTAVSAFYTLNFTGCSTYTSRSGVKKEIRLALPAMGVALAVSLALVLAGRFL--------- 384
WP_250697418.1  289 LPWLPGKSFSVKGVVAG---LIWSGAFYLLADGPA---WGIPTTIALFLALPAVSAFHTLNFTGCSTYTSRSGVKKEMRIALPAMGGALLASVVLLLAGRFL--------- 384
WP_250697419.1  291 LPFIPGRSFAVKGALAG---IAFSAVFFALAGGAL---WSWPVQAASFLAFPAVSSFYTLNFTGCTTFTSRSGVKKEMRLGLPVLGGALLVSLLLVLAGALVY-------- 387
WP_252396620.1  291 LPLLPGRSFAVKGACAG---LLYCALLYALAAGVR---WGGAATAACFLALPAVSAFYTLNFTGCTTYTSKSGVKKEMRLGLPIMAGALGVALIIVIAGRLLV-------- 387
WP_250697417.1  296 LPWLPGKSFSIKGTVTG---VLWCILWYFAGNGAT---WGLPATIGAFLALPAVSAFYALNFTGCSTYTSRTGVKREMRRALPLMGGAVALGVLLFVAGKICVIC------ 394
WP_250697413.1  293 LPWLPTRSFAVKGVAAG---MAWSLLWYLLAGGSG---WDRTTAVAAFLCLPAVSSFYALNFTGSTPFTSRSGVKKEMRRSLPAMGLAVLAGIIVWLAGRFV--------- 388
WP_250697414.1  292 LPWLPTPSFAVKGALAG---LLLALLFAGLSATG-----RRLESAALFLVLPAVSAFFSLNFTGSTPFTSRSGVKKEMRISLPLMAVAVLAGVAAWVAGMLAPLYH---- 389
WP_250697415.1  290 LPWLPGKSFAVKGAVAG---CCWAVLFIGVGRGAT----GWLDAMALFLTVPAVTAFLTLNFTGSTPFTSRSGVKKEMRLGLPMMAVSLLAGISLWIAARFF--------- 384
WP_108102289.1  280 LPWLPCRAFSVKGAVVG---LVMVAAWYLSGISRD---ATVYAKVAAFVALPAVSAFHALNFTGCTPFTSRSGVKKEMRIALPVMGCAVLAGIVLLLIGLIPRQ------- 377
WP_004098500.1  284 LPYIPGPAFAWKGWLLG---LLWAGAY----VWYIASPLSGIHELFYFLILPPIASYLAMNFTGCSTYTSLSGVAKEMNFAIPAQIISAVGGIALLAGGLFLGF------- 380
WP_004626173.1  304 LPWIPGRAFAWKGWQAG---LLWAIALNFLNGWIGPEGLSLLRGVGYLLVLPSISAYYAMNFTGSSTYTSFSGVLKEMRLAIPAIILSLIPGIGLLLADSLIRF------- 404
WP_020491038.1  288 LPWIPGKAFAWKGWLLG---LLWAAAVNVLNGWPEELSYSWFRALGYLLVLPSISAFYAMNFTGSSTYTSFSGVMKEMKTAVPAIAVTICIGCILILLSSLFGL------- 388
WP_010243111.1  301 LPWIPGRAFAWKGWFLG---FIWAITVNILNGWPFSPSYSLIRLLGYLLILPSISSFYAMNFTGSSTYTSLSGVLKEMKIAIPIIIITISSGVLLILIDSFIKL------- 401
WP_207655107.1  302 LPWIPVRAFAAKGWIMG---LIWTIVVNMLNGWPAMPQYGVLKTIAYLLILPSVSSFYAMNFTGSSTYTSFSGVIKEMKIAVPAIAVSIGIGVVLLLINSFIYI------- 402
WP_245693461.1  308 LPWLPGRMFAVKGVWTG----LGAALVLAARFGTRLNGWGL---MALVLAVTAASSYCAMNFTGSTPFTSPSGVEAEMRRALPWQIGAAGLALVAWMVSAWSL-------- 403
WP_155317484.1  307 LPWLPGRAFAVKGAAIG----AAAGALLSAGFRGHLAWMEM---LALTLFIMAVSSYLAMNFTGSTPFTSPTGVEKEMRRAIPLQAAGSLAAAALWVGAGFTG-------- 402
WP_226860105.1  285 LPWIPGRAFSLKGCLAG----LACGFPFWVYYRQDLE-FGS--LLALILWLVAVSSYLAMNFTGATPFTSPTGVEYEMKRAIPAQAAAMVLGFILWVAAPFTG-------- 380
WP_236890013.1  287 LPWLPGRAFALKGGVAG----LASGLLFCAFYWHDFG-VGT--LGALLLWLTAISSYLAMNFTGSTPYTTPSGVEYEMKRAIPAQAIAVVVGLIIWLAAPFTG-------- 382
WP_022666783.1  305 LPYLPFRAFAPKGIIIG----SAASVLLLLMSTHETDSLSR--ILALFLFSTAVSSYLAMNFTGATPFTSPSGVEKEMKQFIPVQAGALAISLGLWIYSAF---------- 399
WP_250697411.1  302 LNRLPWRQFWPKGALVG----GAAGTLAALYLP--VHGWAD--PLALTLWATAVASWQAMNFTGSTPYTSPSGVEKEMRRGMPLQALAALAAAGLWLAGPFLG--------- 396
WP_250697410.1  303 LNRLPWREFWLKGALVG----GAAGGLAALCLSGSVSGFG---AAAMILWALAVGSFTAMNFTGSTPYTSPSGVEREMRRGIPVQAAAALIALALWLGAPFWG--------- 398
WP_250697412.1  305 LPHFPWREFWISGALAG----IVAGVANTWLFNGRLDAWE---SLSLMLWTGAVGSYLAMNFTGSTPYTSPSGVEKEMRRAIPAQVAAGLIAVAVWFAAPFFG--------- 400
WP_200881740.1  306 LPWIPGRAFALKGSLTG----VVAGLAVAVFFG-SRLSWVE--AAALISFTTAISSYLAMNFTGSTPYTSPSGVEKEMRRAIPLQAAVALIAVVAWVGAAFAG--------- 401
WP_252409333.1  305 LPWLPGTAFSVKGLWPG----VFAAAGLLTVMGMDEPMAGA----ALAALSVSVSSYMAMNFTGSTPFTSPSGVEREMRSAMPWQLGGLLVWAVLWVALRFAA--------- 399
WP_246298930.1  314 LPWLPGRAFAFKGAIAAGGLGLLTVLAQAAWAPAAVSSAGR---LAMVLVAGSLGSYLAMNFTGSTPFASPSGVEREMRRAMPLQAVGLVIAAILWIISAF---------- 411
WP_205232271.1  300 LPWIPGRAFSLKGALLA----MATTPAFVLVADLQVLET-----TALVLVFTSISSHLAMNFTGSTPYTSPSGVEKEMKRALPLQGLALAAGIVMWLAAPFVNI------- 394
WP_250697420.1  292 LPWLPGRQFWIKGLWPS----LLAGTALLLSTAVPPAPVDQ---IALLLWIVAVSSFQAMNFTGCTPYTSPSGVEFEMRRGVPLQILLAVAATLLWLASPFLNT------- 388
WP_250697404.1  292 LPWLPGRQFWLKGVWAG----LPAALGAWLLLAARLAAVEQ---VALISWVLSLSSYLAMNFTGSTPFTSPSGVEYEMRRGLPVQLAATCLALVLWLASPFLH-------- 387
WP_250697403.1  289 LPWLPGRQFWLKGLLPG----LLSGGLC-WLFAAPLAVVEG---LALLFWSLSIASFLAMNFTGSTPFTSPSGVEYEMKRGLVVQFALTGLALTLWLLAPFL--------- 382
WP_250697406.1  300 LPWLPCRQFWLKGLWPP----LLVALCWGIN-IYPHVTIEL---LSLSVWMSVFSSFQAMNFTGCTPYTSPSGVEYEMRRGIPVLALLAILALLVWLVAPFVD-------- 394
WP_250697405.1  294 LPWLPGRQFWIKGLFPG----LLVGLWCLWGCDTVLSDFEK---AAMALWSVAISSYLAMNFTGSTPFTSPSGVEYEMKRGLLVQCLCAAGALVLWLLSPFVI-------- 389
WP_252405647.1  302 LPWIPFRQFWLKGTVTG----SITAMLFLLGIQPAVNGPEK---LAIFLFITTCAAYLAMNFTGSTPFTSLSGVEKELRRGLPVQIGAAALALLIWLAGPFIP-------- 397
WP_252405646.1  294 LPWIPARQFWLKGAVVG----ALLAFLYLAAVAGRPNSFEL---LSMFLWICGCSSYLAMNFTGSTVFTSLSGVKKEMTRGLKFQLICTAAAVICWVAAPF---------- 387
WP_250697409.1  301 LPWIPSRQFWLKGALVG----GLTAMTAMAALLPTTWTADLPAAIALVLWAASVASYLAMNFTGSTPYTSLSGVGLEMRRGLAFQLGGTTLALGLWIWSAFA--------- 398
WP_250697408.1  303 LPWLPGRQFWLKGLSVS----IFAGLIFTSLSS--SGVTAT---LALALWILAVGSYLAMNFTGSTPYTSLSGVEAEMRKGLPVQIGLAALALLLWLVAPFITQA------ 398
WP_250697407.1  294 LRALPFRQFWLKGAFTS----LISALLFSMWSIPIAGSHDT---LALASWMLATGSFLAMNFTGSTPYTSLSGVEFEMRRGLPLQIALATIALVLWLISPFIRG------- 390
WP_252396621.1  306 LPWLPGRAFSIKGALTG----FGTGTIAAFMG--DSFLAGS----ALVLFSTAISTYLAMNFTGSTPFTSPSGVEKEMRYALPWIAGGFVLSVLLWIISGLSA-------- 398
WP_252396623.1  293 MSRLPGRMFWLHGALLG----AIAAACTAFMV-----ASGP----GPVLIGTAAGSFMMMNFTGSTPFTSPTGVEHEMKRGIPFQLVTALLGLALWIVMPFITGGA---- 385
WP_250697582.1  315 LRLLPGRAFAAKGLLAG------AAVGLPLAVLLARSVPEG---LAALSICAAFSSWFAMHYTGSTPFTSLSGVDREMRRYMPAQGLLAFLAVMLWLGWSWFGGPAAGGS 415
```
